# The Role of the Receptor for Advanced Glycation End-Products in Cancer: Evidence from a Systematic Review and Meta-Analysis

**DOI:** 10.64898/2026.02.13.705839

**Authors:** Michael B. Nelappana, Pawel Wityk, Catherine C. Applegate, Goodluck Okoro, Leszek Kalinowski, Iwona T. Dobrucki, Lawrence W. Dobrucki

**Author notes:** Authors contributed equally to the manuscript (Catherine C. Applegate, Goodluck Okoro). **Corresponding author:** Professor Lawrence W. Dobrucki, University of Illinois at Urbana-Champaign, 405 N. Mathews Ave, MC-251, Urbana, IL 61801. The authors declare no potential conflicts of interest.

## Abstract

The Receptor for Advanced Glycation End-products (RAGE) has been implicated in driving cancer growth, aggression, and metastasis through the fueling of chronic inflammation in the tumor microenvironment. This systematic review and meta-analysis summarize and analyze current clinical and preclinical data to provide insight into the relationship between RAGE and cancer, cancer grade, metastasis, patient survival, and cellular processes. A multi-database search was performed to identify original clinical and preclinical research studies examining RAGE expression in cancer. After screening and review, 53 clinical and 233 preclinical studies were included. Associations of RAGE with clinical cancer outcomes were estimated using odds ratio (OR) and associated 95% confidence intervals (CI). The meta-analysis found that RAGE expression was highly correlated with cancerous tissue when compared to controls; high-grade tumors; regional lymph node invasion; and was somewhat negatively associated with patient survival. In addition, meta-analysis estimates of preclinical studies found positive associations between RAGE expression/activation and cancer growth, metastatic potential, evasion of apoptosis, and activated NF-κB expression. This systematic review and meta-analysis is the first comprehensive study through which both preclinical and clinical research in all available cancer types are assessed for correlations with RAGE expression and activation, demonstrating that RAGE does indeed play a significant role in cancer progression and that further research is warranted.

## Introduction

Chronic inflammation associated with increased obesity has been identified as a key part of both the carcinogenic process and the development of more aggressive cancers^1^. In particular, the Western Diet has been strongly linked to the development of obesity, obesity-associated carcinogenesis, and the development of systemic chronic inflammation^2^. The Western Diet and its subsequent diet-induced pathologies have also been linked with increased cancer-associated mortality and morbidity^2,3^. Comprised largely of animal protein and high-carbohydrate and high-fat processed foods, the Western diet is also rich in dietary Advanced Glycation End-products (AGEs)^4^. AGEs are stable end-products formed endogenously and exogenously through the nonenzymatic glycation of proteins, lipids, and nucleic acids that can form toxic crosslinks with other molecules and bind to specific inflammatory receptors^5^.

The Receptor for Advanced Glycation End-products (RAGE) is a member of the immunoglobulin protein family of cell surface proteins found in a wide range of tissue types, with increased presence in lung tissue^6^. RAGE activation by AGEs stimulates PI3K-mediated activation of NF-κB that leads to a positive feedback loop of pro-inflammatory responses, including the increased expression of RAGE^7–9^. Inflammation in the tissue microenvironment has been linked to proliferation, apoptosis inhibition, treatment and immune resistance, angiogenesis, and epithelial-mesenchymal transition (EMT)^10,11^. Activation of RAGE can also be induced by a wide range of ligands, such as the S100 protein family and high mobility group box 1 protein (HMGB1), a damage-associated molecular pattern (DAMP) molecule released by injured or dying cells^9^. Activation of RAGE by proteins like HMGB1, released by cells that die during therapies such as radiation, suggests that RAGE may play a role in cancer treatment resistance. Our research group has looked to the AGE/RAGE as a potential target for diagnostic and therapeutic actions due to its critical role in the inflammation of the cancer microenvironment. We have published work investigating the use of a multimodal nanoparticle targeted at RAGE for imaging prostate cancer, and we found that not only did RAGE allow for high-specificity imaging of cancerous tissue by PET/CT, but also that RAGE expression had a strong positive correlation to Gleason score^12,13^.

Despite the body of studies demonstrating the important role the AGE/RAGE axis has been shown to play in cancer, only one systematic review or meta-analysis evaluating RAGE expression in clinical cancer specimens exists. This systematic review and meta-analysis was recently published by our group, assessing RAGE as a novel biomarker in prostate cancer, where we demonstrated a strong positive association between RAGE expression, cancerous tissue, and high-grade cancer^14^. As such, we wanted to expand the scope of that systematic review and meta-analysis to evaluate all cancers. We hypothesized that RAGE expression and activation is strongly correlated with clinical cancerous tissue as well as clinical tissue grading and preclinical cancer progression.

In this systematic review and meta-analysis, we assessed the relationship between RAGE expression in low– and high-grade cancer specimens and those observed in benign or adjacent normal tissues. We additionally expanded our systematic review to include clinical specimens of cancer that invaded regional lymph nodes and assessed the relationship between RAGE expression and patient survival. Furthermore, preclinical works investigating the relationship between RAGE activation and cancer growth, metastasis, apoptosis evasion, and activated NF-κB expression were assessed. The findings from this review provide new evidence that RAGE may serve as a biomarker not only for cancer diagnosis but also for distinguishing between benign tissue, low-grade (low-risk) cancers, and high-grade (high-risk) cancerous tissue.

## Materials and Methods

### Study selection criteria

This clinical meta-analysis was conducted in accordance with the Cochrane Handbook and the 2020 PRISMA guidelines for meta-analyses^15,16^. Studies that met the following criteria were included in the meta-analysis: (a) used validated cancer samples against appropriate control samples (benign tissue or adjacent normal tissues); (b) directly measured RAGE expression by validated techniques; (c) methodology was documented in replicable detail; (d) evaluated the relationship between RAGE expression and cancer; and (e) written in English.

Currently, no validated guidelines or tools exist for conducting systematic reviews and evaluating the validity and quality of mechanistic studies through meta-analyses. As part of an effort to utilize a cohesive and standardized set of guidelines for systematically reviewing and pooling evidence from preclinical studies, this systematic review was conducted in accordance with the framework outlined by the World Cancer Research Fund (WCRF) International and the University of Bristol (UoB)^17^. In addition, care was taken to follow the PRISMA reporting guidelines as closely as possible^16^. Preclinical studies that met the following criteria were included in this systematic review: (a) examined RAGE expression in a validated cancer cell line or animal model; (b) quantitatively evaluated the direct relationship between RAGE expression and cancer outcomes quantitatively through cell culture studies utilizing validated growth or functional assays or through animal studies utilizing validated *in vivo* or *postmortem* assessments; (c) includes RAGE-targeted treatment, activation, or intervention; (d) methodology was documented in replicable detail; (e) included appropriate controls, and (f) was written in English.

### Literature search

To reduce the risk of publication bias, we included grey literature in our search strategy. As such, we conducted (up to April 13, 2024) a comprehensive literature search of PubMed, Web of Science, and Scopus using a combination of the following keywords and their variants: cancer, carcinoma, receptor for advanced glycation end products, advanced glycation end product(s). Titles and abstracts of studies identified by the keyword search were screened against the study selection criteria. Potentially relevant manuscripts were retrieved for evaluation of the full text. We also conducted a reference list search (backward search) and cited reference search (forward search) from manuscripts meeting the study selection criteria. Studies identified through this process were further screened and evaluated using the same criteria until no further relevant studies were found. An updated search was performed between April 13^th^, 2024, and July 11^th^, 2025, to include recently published studies. Four authors (MN, CA, GO, PW) individually determined the inclusion/exclusion of all studies retrieved in full text, and discrepancies were resolved through discussion.

### Data extraction and quality assessment

The following information was extracted from each study for the clinical meta-analysis: name of the first author; year of publication; number of cases, controls, and total number of participants in the study; methodological details describing RAGE expression measurements; patient survival, histological grade, lymph nodal invasion, and study type. A number of cases (designated RAGE positive by the authors of the individual studies) within both the total number of cancer cases and controls were used to calculate the odds ratio (OR) and 95% confidence intervals (CI) that were subsequently used to perform the meta-analysis. For the secondary outcomes, a grade of 1-2 or description of well to moderate differentiation was considered to be low-grade cancer, while a grade of 3-4 or description of poor differentiation was considered to be high-grade cancer; a grade of Nx-N0 on the TNM system was considered to have no lymph nodal invasion, while a grade of N1+ was considered to be indicative of lymph nodal invasion; and survival was assessed through end-point patients who were still alive.

The quality assessment (QA) of each study was performed using the Newcastle-Ottawa Scale, which is a validated scale for non-randomized cohorts in a meta-analysis^18^. This tool judges the literature based on the following three categories: selection of cases and controls, comparability of studies, and exposure of the main variable (RAGE). We regarded scores of 1-3, 4-5, and 6-8 as low, moderate, and high quality, respectively. QA scores were used to measure the strength of the evidence provided by each study but were not used to determine the inclusion of studies.

For preclinical studies, data extraction was performed using the recommendations set forth by the WCRF/UoB framework as a guide^17^. The following information was extracted from each cell culture study: names of cell lines; whether cell lines were established patient-derived tumor cell lines or freshly isolated primary cells; whether cell lines were authenticated; culture conditions; treatment regime (dose and length of treatment), if any; details of laboratory procedures; RAGE-related outcomes analyzed (growth, metastatic potential, apoptosis evasion, and activated NF-κB expression); results of RAGE-related outcomes; sample size and standard deviation (SD) or standard error of the mean (SEM); statistical test conducted; and *p*-values. The following information was extracted from each animal study: animal model used; names of cell lines grafted or inoculated; treatment regime (dose and length of treatment), if any; details of laboratory procedures; RAGE-related outcomes analyzed (tumor growth and metastasis); results of RAGE-related outcomes; sample size and SD or SEM; statistical test conducted; and p-values.

There is a lack of validated QA tools to evaluate the risk of bias associated with preclinical studies involving cells and experimental animals. QA of cell culture studies included in this review was performed using adapted criteria recommended by the WCRF/UoB framework and other published recommendations (score range: 0-8; a score of 0 was assigned for each parameter not fulfilled or not reported)^17,19^. Based on the score, studies were rated as low (0-3), moderate (4-6), or high (7-8) quality. QA of animal studies was performed using the SYstematic Review Centre for Laboratory animal Experimentation (SYRCLE) risk of bias tool adapted from the established and validated Cochrane tool for human study risk of bias assessment^20,21^. Risk of bias was determined to be “high,” “low,” or “unclear.” Total scores were not evaluated using the SYRCLE tool to avoid inappropriate weighting of each category. QA scores were utilized to provide a measure for the strength of the evidence and to determine if a risk of bias was present for each study and were not used to determine the inclusion of studies. Conclusions based on whether the included studies supported the biological plausibility of the causal pathway being investigated were based on the QA.

### Statistical analysis

STATA/IC version 18.0 (StataCorp LP, College Station, TX) was utilized to analyze the data. OR and 95% CI were used as a measure of the effect size for all studies. Heterogeneity among studies was assessed using the *I*^2^ statistic based on Cochran’s *Q*^22^. Random (DerSimonian-Laird) effects models were applied due to high *I*^2^ values (≥50%), indicative of increased study heterogeneity. Potential publication bias was assessed using visual assessment of funnel plots, and funnel plot asymmetry was evaluated using Harbord’s test for small study effects, which is a modified version of Egger’s linear regression test^23–25^. We also performed sensitivity “leave-one-out” analyses to evaluate whether the pooled results could differ if a single study at a time was excluded. Figures were generated through Python scripts from the STATA/IC analysis.

The extreme degree of heterogeneity between the methodologies and outcome measures of preclinical studies makes conducting a true statistical analysis via a meta-analysis exceedingly difficult. In lieu of a true meta-analysis, effect estimates for each study outcome were calculated through the generation of albatross plots. An albatross plot, as described by Harrison *et al.,* scatters the *p*-values of each study according to their sample size and according to the observed direction of the effect on the outcome (positive or negative)^26^. In the absence of exact *p*-values provided, the most conservative *p*-value was assigned to that outcome (e.g., if given *p*<0.05, set *p*=0.05). Non-statistically significant *p*-values without an exact given value were assigned the most liberal *p*-value to that outcome (e.g., if given *p*>0.05 or ns, set *p*=0.5). The contour lines extending over the plot represent estimated effect sizes (represented as standardized mean difference, SMD) to allow for the estimation of the magnitude of effects for individual studies. Visual inspection of the albatross plots determined an overall estimated standardized effect for each outcome, which represents the strength of the associations using the absolute values for the calculated beta-coefficients. As such, a larger beta coefficient represents a larger standardized effect on that outcome. To provide additional information, meta-analyses of *p*-values were conducted for each outcome using the weighted Z-test^27,28^. The weight used was 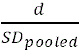, where *d* was equal to the direction of the effect on the outcome (positive = 1; negative = –1) and *SD_pooled_* was the pooled standard deviation of the study. Albatross plots were generated using Python scripts. A *p*-value of less than 0.05 was considered statistically significant for all analyses.

### Data availability statement

This systematic review and meta-analysis were not registered under any program. The data generated in this study are available within the article and its supplementary data files. Additional information pertaining to literature search results and data extraction are available from the authors upon request.

## Results

### Literature search

In total, 10,329 studies evaluating RAGE in cancer were identified from the library search engines. This process is summarized in **Figure 1**. After removing duplicates and screening abstracts for pertinent information, 417 studies remained and were evaluated by full-text review. Manuscripts found not to meet inclusion criteria did not measure RAGE expression in association with the listed outcomes. Two hundred and twenty-six (226) manuscripts were found to meet the inclusion criteria and were included in the final analysis: 52 studies measuring clinical expression of RAGE in cancer^29–80^; 160 studies measuring *in vitro* effects of RAGE expression on cancer growth, metastatic potential, apoptosis evasion, and activated NF-κB expression^A^; and 73 studies measuring *in vivo* effects of RAGE expression on tumor growth and metastasis^B^. Clinical studies were further stratified according to the measured RAGE-dependent outcomes: RAGE expression’s association with cancerous tissue was evaluated in 38 studies^C^; with histological grading in 29 studies^D^; with lymph node metastasis in 19 studies^E^; and with patient survival in 9 studies^F^. Cell culture studies were further stratified according to the measured RAGE-dependent outcomes: 124 studies evaluated cancer cell proliferation^G^; 95 studies evaluated invasion and/or migration of cancer cells^H^; 32 studies evaluated apoptosis evasion of cancer cells^I^; and 24 studies evaluated expression of activated/phosphorylated NF-κB in cancer cells^J^. Animal studies were further stratified according to the measured RAGE-dependent outcomes: 69 studies evaluated tumor volume, weight, and/or other growth assessments^K^; and 25 studies evaluated metastatic incidence, number, tumor burden, and/or other assessments of metastasis^L^.

**Figure 1.**
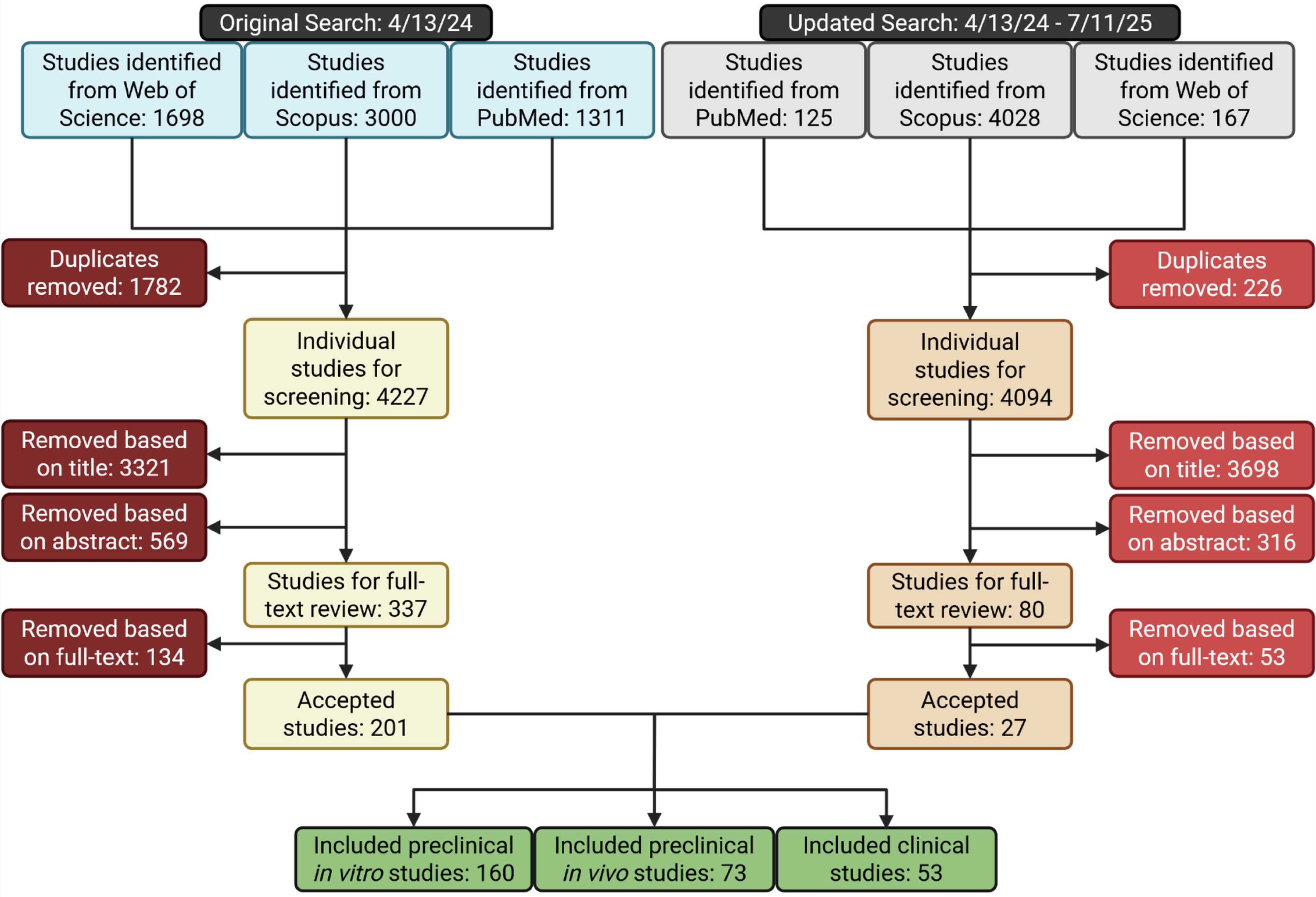
Schematic of workflow for identifying relevant articles. Studies were identified through keyword search in the listed databases on 4/13/24. Papers were filtered for duplicates before being screened by title and abstracts. Full-text review was performed using the stated inclusion criteria and accepted studies then underwent data extraction. An update to the paper search was performed on 7/11/25 and resulting papers underwent the same process. Created with BioRender.com

### Study characteristics

All 52 clinical studies were retrospective case-control studies examining the associations of RAGE expression in cancer samples. Fourteen (14) semi-quantitatively assessed the association of RAGE expression with cancer samples vs. adjacent normal tissue^29,31,37,49,50,55,56,67,68,72,74,78–80^, 25 with high grade vs. low grade cancerous tissue^29,33–35,37,38,41,42,45,50,51,53,55,56,61,65–68,72–74,78–80^, 19 with invasion of regional lymph nodes^33–35,42,45,49–51,53,54,65–68,72–74,79,80^, and 9 with patient survival^47,51,52,54,59,65,67,68,80^. Furthermore, 26 quantitatively assessed expression of RAGE with cancer samples vs. adjacent normal tissue^30,32,35,36,38–40,42–44,46,48,54,58–60,63,64,68–71,75–78^, and 4 with high grade vs. low grade cancerous tissue^40,57,58,62^. Study characteristics are summarized in **Supplementary Table S1**. The total number of study participants included was 13,915 and the total number of cancer cases reported was 7,868, though some participants in either sum were counted twice if counted in two or more semi-quantitative assessments. All semi-quantitative studies reported incidence data as positive vs. negative RAGE expression, enabling calculation of ORs and corresponding 95% CIs. All quantitative studies reported incidence data as means and standard deviations, enabling calculation of SMDs

Quality scores (QA) were assigned to each article using the criteria outlined by the Newcastle–Ottawa scales for case-control studies. The average score for the studies was 6.1±1.6, the lowest score being 2 (low quality) and the highest being 8 (high quality) on the 8-point scale. Seven studies were considered low quality, two studies were considered to be of moderate quality, and 43 studies were considered to be high quality. QA scores for individual studies are provided in **Supplementary Table S2**.

Cell culture study characteristics and results are summarized in **Supplementary Table S3**. 92 cell culture studies identified used a single verified cancer cell line in their analyses^M^, 67 used multiple verified cancer cell lines^N^, and one generated their own unverified cancer cell line^214^. In addition, five studies used cell lines from two different organs of origin^137,140,150,152,190^ and one from three different organs of origin^96^, making 167 unique reports. One hundred and fifty-two (152) reports confirmed RAGE expression in association with measured outcomes with the remaining 15 directly treating RAGE via AGE-RAGE axis activators, inhibitors, or transfection. One hundred and thirty-one (131) reports directly modulated the AGE-RAGE axis to examine its effects on cancer cells, 97 examined effects on metastatic potential, 33 examined effects on apoptosis, and 28 examined effects on activated NF-κB expression.

QA of cell culture studies was carried out using criteria adapted from the WCRF/UoB framework for evaluating mechanistic studies. The criteria used to evaluate the quality of the cell culture studies were vague, resulting in 74 cell culture studies considered to be high quality (7-8) and 87 cell culture studies considered to be of moderate quality (5-6). The average score for the studies was 6.5±0.76, the lowest score being 5 (moderate quality) and the highest being 8 (high quality). QA scores for individual studies are provided in **Supplementary Table S4**.

Animal study characteristics and results are summarized in **Supplementary Table S5**. Of the 87 animal models used, 73 were xenografts on verified cancer cell lines^O^, nine were transgenic animals or induced-cancer models^189,210,228,239,245–247,249,251^, and five were xenografts of unverified cancer cell lines^214,230,236,240,252^. Seventy-nine (79) reports directly modulated the AGE-RAGE axis to examine its effects on tumor growth and 27 examined effects on metastatic growth.

QA of animal studies were carried out using criteria adapted from the SYRCLE risk of bias tool. The SYRCLE tool resulted in a largely unclear risk of bias for all but one study which had a high risk of bias. QA assessments for individual studies are provided in **Supplementary Table S6**.

All accepted studies were graphed to provide an understanding of the landscape and frequency of publishing research related to RAGE and cancer (**Supplementary Figure S1**).

### RAGE expression in clinical cancer samples

Fourteen (14) clinical studies semi-quantitatively evaluated the expression of RAGE in cancer specimens compared with adjacent normal tissue samples, mainly using immunohistochemistry (IHC). A random-effects meta-analysis was performed based on the DerSimonian-Laird approach to account for between-study heterogeneity (*I*^2^=88.65%, Cochran’s *Q p*=0.00). Meta-analysis of these studies confirmed the OR was 3.49 (95% CI: 1.53-7.95) indicating a strong likelihood for cancerous tissue to express RAGE compared with benign tissues (**Figure 2**). A trend was noticed wherein gastrointestinal and respiratory system cancers were either not being associated with or negatively associated with RAGE expression. To further investigate this, these cancers were grouped and designated as ‘nulli-RAGE’ to indicate their negative/null association with clinical cancers and all other cancers were stratified as ‘pro-RAGE’. Meta-analysis of the grouped studies confirmed that the nulli-RAGE OR was 0.60 (95% CI: 0.13-2.84), indicating no association between RAGE expression and cancerous tissue and that the pro-RAGE OR was 8.54 (95% CI: 4.35-16.79), indicating a very high likelihood for cancerous tissue to express RAGE compared with adjacent normal tissue.

**Figure 2.**
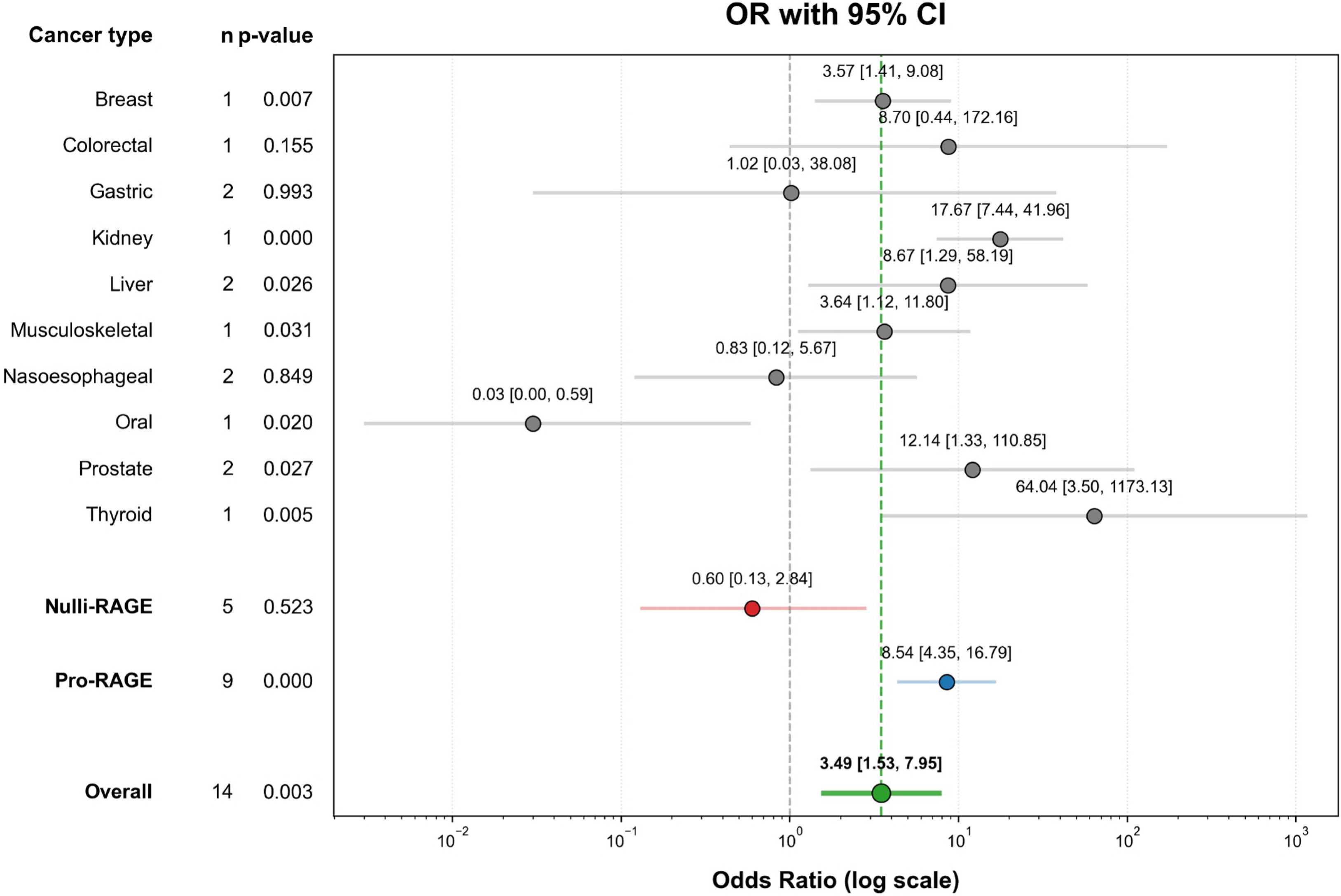
Forest plot of receptors for advanced glycation end-products (RAGE) expression in cancer. These associations were indicated as odds ratio (OR) estimates with a corresponding 95% confidence interval (CI). Vertical dashed grey line indicates OR of 1 or null association. Results are indicated by grey circles, indicating OR estimate, and horizontal grey lines, indicating 95% CI. Vertical dashed green line and green dot and horizontal line indicate the overall OR estimate. Stratified results for the nulli-RAGE (red dot and horizontal line) and pro-RAGE (blue dot and horizontal line) groups are also indicated.

Funnel plots were used to visually assess for publication bias in the full meta-analysis (FMA) as well as the stratified RAGE categories (**Supplementary Figure S2**). As such, we examined the studies for small-study effects using Harbord’s test and found no significant effect in the FMA (*p*=0.8685) and pro-RAGE stratification (*p*=0.3953), however significant effect was found in the nulli-RAGE stratification (*p*<0.0001) indicating small-study effects. Furthermore, leave-one-out sensitivity analyses consistently demonstrated elevated ORs for FMA, ranging from 2.90-4.53 (**Supplementary Figure S3A**), and pro-RAGE, ranging from 7.03-10.14 (**Supplementary Figure S3B**), and demonstrated consistently reduced ORs for nulli-RAGE, ranging from 0.32-0.93 (**Supplementary Figure S3C**).

### RAGE expression in high– vs. low-grade cancer

Twenty-five (25) clinical manuscripts additionally evaluated the expression of RAGE in high-vs. low-grade cancer specimens. Data were reported as histological grade and/or as designated by the individual study; low-grade was considered to be grade 1-2 and/or designated as “low-grade”, and high-grade cancer was considered to be grades 3-4 and/or designated as “high-grade”. Study heterogeneity continued to be present (*I*^2^=64.58%, Cochran’s *Q p*=0.00), so a random-effects DerSimonian-Laird model was performed. FMA confirmed an OR of 1.55 (95% CI: 1.03-2.35) indicating a likelihood for high-grade cancerous tissue to express RAGE compared with low-grade cancerous tissues (**Figure 3**). A disparity was again found between the stratified cancer types with the nulli-RAGE OR (0.88; 95% CI: 0.45-1.69) indicating no associations between RAGE expression and cancer grade and the pro-RAGE OR (2.22; 95% CI: 1.31-3.75) indicating a strong likelihood for high-grade cancerous tissue to express RAGE compared with low-grade cancers.

**Figure 3.**
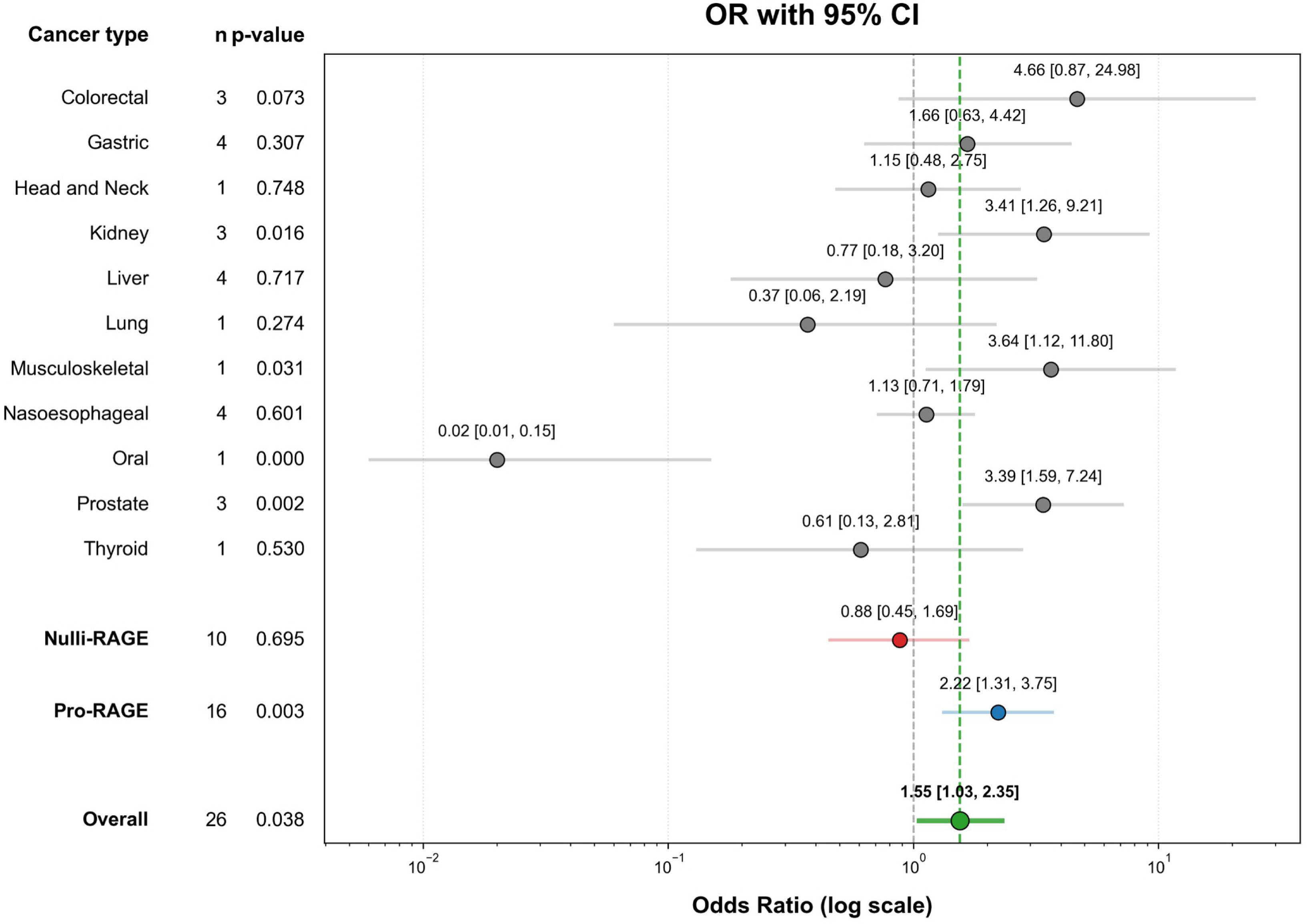
Forest plot of RAGE expression in high-grade cancer. These associations were indicated as odds ratio (OR) estimates with a corresponding 95% confidence interval (CI). *Vertical dashed grey* line indicates OR of 1 or null association. Results are indicated by *grey circles*, indicating OR estimate, and horizontal *grey* lines, indicating 95% CI. Vertical *dashed green* line and *green dot and horizontal line* indicate the overall OR estimate. Stratified results for the nulli-RAGE *(red dot and horizontal line)* and pro-RAGE *(blue dot and horizontal line)* groups are also indicated.

Funnel plots for the FMA as well as the stratified RAGE categories (**Supplementary Figure S4**) were inspected visually. The Harbord’s test for all three groups found no significant small-study effects ((FMA; *p*=0.2042) (pro-RAGE; *p*=0.3142) (nulli-RAGE; *p*=0.5689)). Furthermore, leave-one-out sensitivity analyses demonstrated slightly elevated ORs for FMA (1.43-1.72), elevated ORs for pro-RAGE (1.94-2.45), and demonstrated somewhat reduced ORs for nulli-RAGE (0.82-1.28) (**Supplementary Figure S5**).

### RAGE expression in lymph-invading cancers

To assess the relationship between expression of RAGE and cancer cases that invaded regional lymph nodes, 19 clinical manuscripts were additionally evaluated. Data were reported as N0-3; regional lymph node invasion was considered to be N1-3, and no regional lymph node invasion was considered to be N0. A random-effects DerSimonian-Laird model was performed due to study heterogeneity (*I*^2^=77.34%, Cochran’s *Q p*=0.00). FMA confirmed an OR of 2.72 (1.76-4.20), indicating a strong likelihood for cancer cases that invaded regional lymph nodes to express RAGE compared with cancer cases that did not (**Figure 4**). The disparity between the stratified cancer types was clear with the nulli-RAGE OR (1.38; 95% CI: 0.88-2.17) indicating no associations between RAGE expression and regional lymph node invasion and the pro-RAGE OR (5.13; 95% CI: 3.21-8.19) indicating a very high likelihood for cancer cases that invaded regional lymph nodes to express RAGE compared with cancer cases which did not.

**Figure 4.**
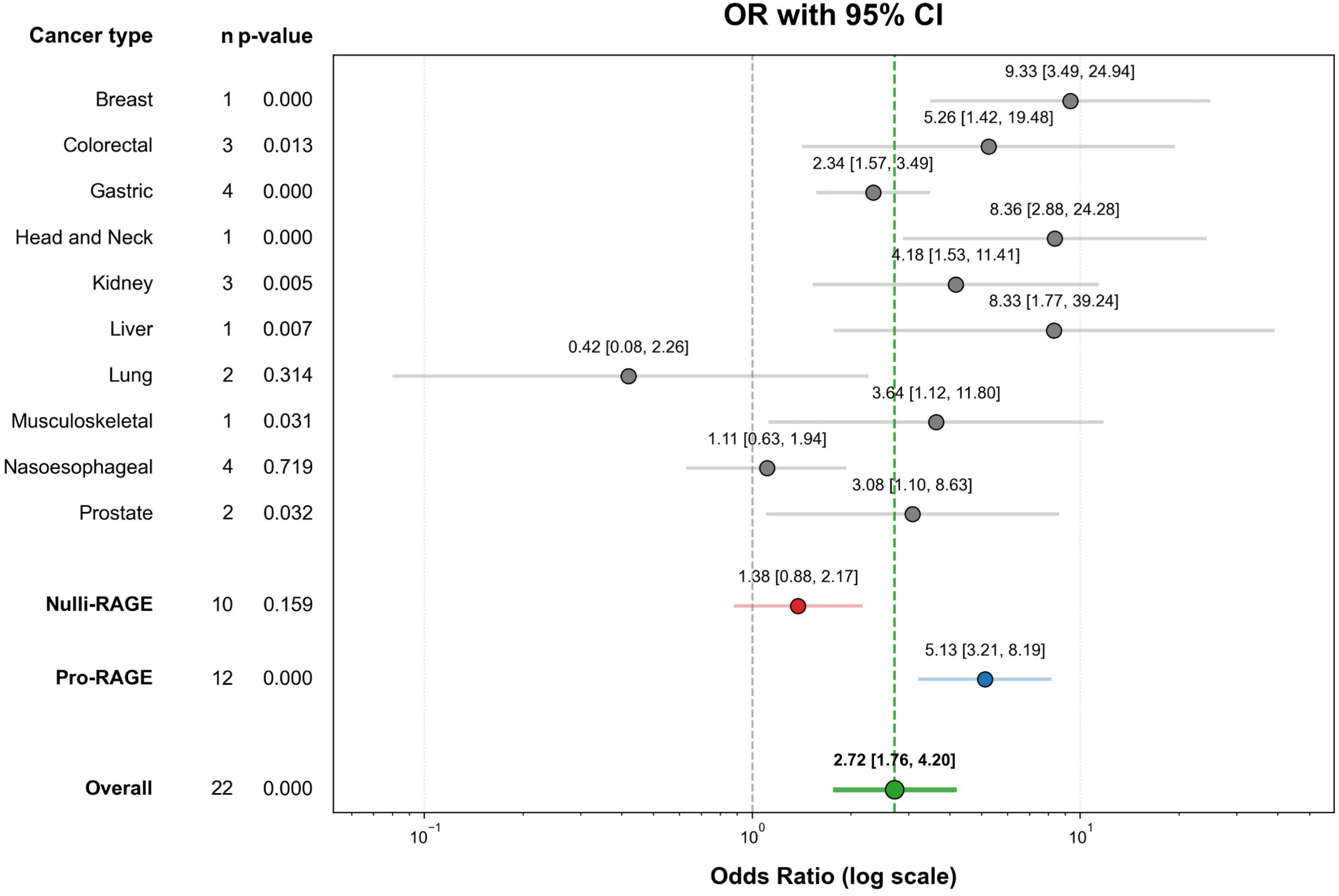
Forest plot of RAGE expression in cancers that invaded regional lymph nodes. These associations were indicated as odds ratio (OR) estimates with a corresponding 95% confidence interval (CI). Vertical dashed grey line indicates OR of 1 or null association. Results are indicated by grey circles, indicating OR estimate, and horizontal grey lines, indicating 95% CI. Vertical dashed green line and green dot and horizontal line indicate the overall OR estimate. Stratified results for the nulli-RAGE (red dot and horizontal line) and pro-RAGE (blue dot and horizontal line) groups are also indicated.

Funnel plots for the FMA as well as the stratified RAGE categories (**Supplementary Figure S6**) were inspected visually, and while slightly asymmetrical, our search strategy attempted to minimize these effects by including unpublished (“grey”) literature. The Harbord’s test for all three groups found no significant small-study effects ((FMA; *p*=0.5579) (pro-RAGE; *p*=0.8272) (nulli-RAGE; *p*=0.8846)). Furthermore, leave-one-out sensitivity analyses demonstrated consistently elevated ORs for FMA, ranging from 2.43-2.97, and pro-RAGE, ranging from 4.20-6.00, and demonstrated slightly elevated ORs for nulli-RAGE, ranging from 1.27-1.53 (**Supplementary Figure S7**).

### RAGE expression and patient survival

Finally, for clinical samples, the association of RAGE expression and patient survival was evaluated using 9 clinical manuscripts. Data were reported as patients; living patients were considered as cases, and deceased patients were considered as controls for the purposes of calculating an OR. Study heterogeneity continued to be present (*I*^2^=86.58%, Cochran’s *Q p*=0.00), so a random-effects DerSimonian-Laird model was performed. FMA confirmed an OR of 0.70 (0.37-1.33), indicating no associations between RAGE expression and patient survival (**Supplementary Figure S8**). The stratified cancer types, however, elucidated an effect with the nulli-RAGE OR (1.12; 95% CI: 0.51-2.43) continuing to indicate no associations between RAGE expression and patient survival, but the pro-RAGE OR (0.42; 95% CI: 0.27-0.68) indicated a high likelihood for deceased patients to express RAGE compared with patients who survived.

Funnel plots for the FMA as well as the stratified RAGE categories (**Supplementary Figure S9**) were inspected visually, and were found to be slightly asymmetrical but, as mentioned above, minimized through the inclusion of “grey” literature. When examining the three groups for small-study effect with the Harbord’s test, FMA (*p*=0.0358) and nulli-RAGE (p=0.0072) were found to have significant small-study effects and pro-RAGE (*p*=0.0720) was found to be close to a significant small-study effect. Furthermore, leave-one-out sensitivity analyses demonstrated consistently reduced ORs for FMA, ranging from 0.59-0.82 (**Supplementary Figure S10A**), strongly reduced ORs for pro-RAGE, ranging from 0.32-0.52 (**Supplementary Figure S10B**), and demonstrated mildly elevated ORs for nulli-RAGE, ranging from 0.86-1.49 (**Supplementary Figure S10C**).

### Effect of RAGE on cancer cell growth potential

One hundred and twenty-four (124) of the 160 cell culture studies evaluated the association between AGE-RAGE axis activation or expression and cancer cell growth potential, as assessed by cell viability and proliferation assays. Despite large variation in the methodologies used, the types of treatments performed, and the doses utilized, 324 of the 371 (87.33%) results using cancer cell lines reported increased AGE-RAGE axis activation or expression resulting in increased cancer cell growth potential. To maintain consistency, gastrointestinal and respiratory cancers were again categorized as nulli-RAGE cancers, and all other cancer types were categorized as pro-RAGE for all preclinical studies. Two hundred and sixty-seven (267) of 301 (88.70%) pro-RAGE results and 57 of 70 (81.43%) nulli-RAGE results also reported that increased AGE-RAGE axis activation or expression resulted in increased cancer cell growth potential. An in-depth description of individual study results can be found in **Supplementary Table S3**.

An albatross plot (**Figure 5A**) was generated to integrate the data, and visual inspection of the plot provided an estimated standardized effect between AGE-RAGE axis activation or expression and cancer cell growth potential. The overall SMD range, representing a wide range of effects, was –17.89 to 500.96. Sixty-four results showed no effects of RAGE on cancer cell growth with an SMD below ±0.97. Eight results showed mild negative effects, and 41 results showed mild positive effects of RAGE on cancer cell growth, with an SMD between ±0.97 and ±1.94. Two results showed moderate negative effects, and 43 showed moderate positive effects of RAGE on cancer cell growth, with an SMD between ±1.94 and ±2.91. Eighteen results showed strong negative effects, and 195 results showed strong positive effects with an SMD greater than ±2.91.

**Figure 5.**
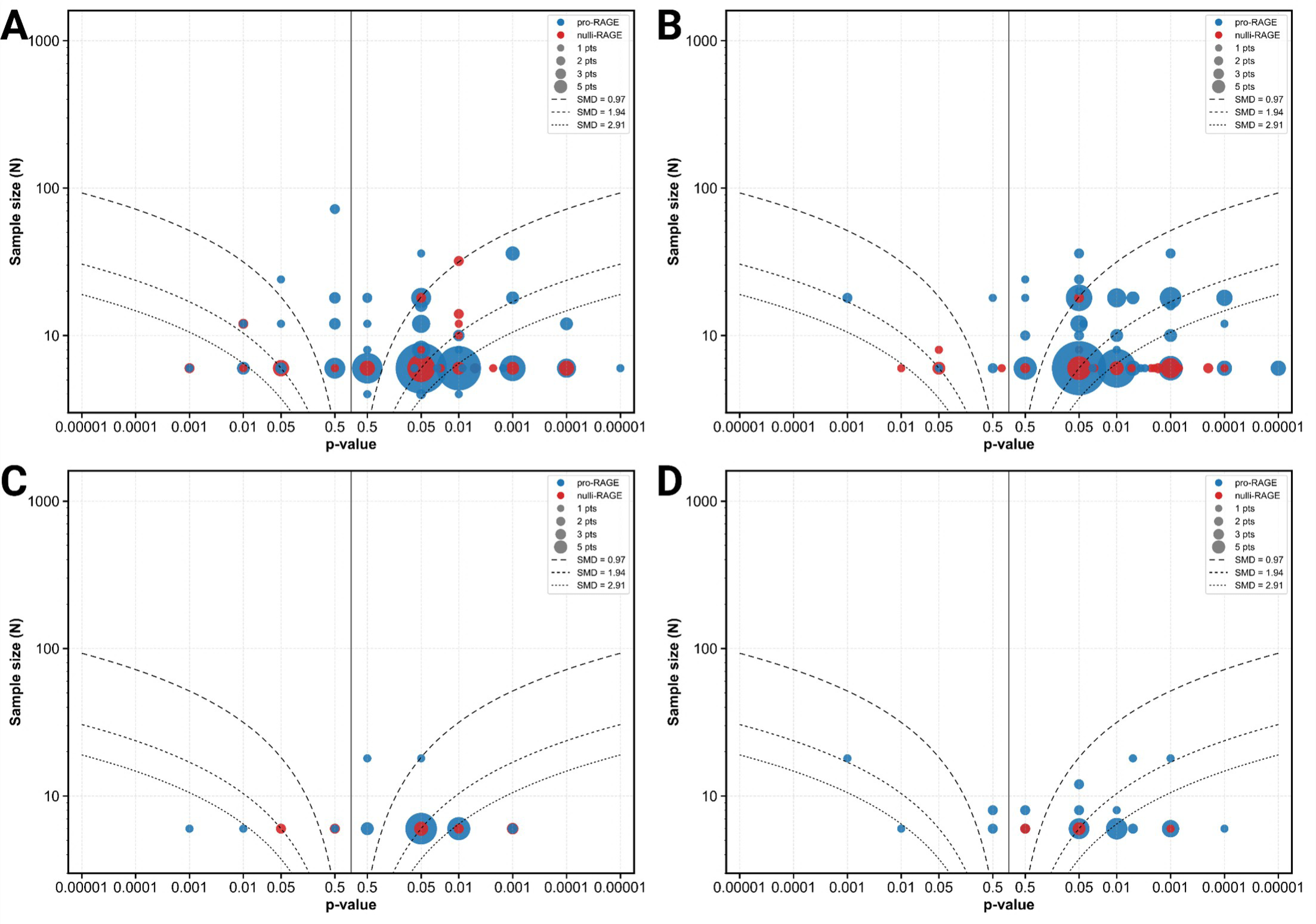
Albatross plot of cell culture studies measuring AGE-RAGE axis association and modulation with. (**A**) cancer cell growth potential, measured by growth assays; (**B**) cancer metastatic potential, measured by invasiveness and migration assays; (**C**) cancer apoptosis evasion, measured by molecular assays; and (**D**) cancer activated NF-kB expression, measured by luminescent assays or Western blots. Each point represents data and its size is correlated to number of overlapping points, with the effect estimate (represented as a p-value), plotted against the total given sample size (n) included within each study. The directionality of the p-values correlate to negative associations with RAGE for the left and positive associations with RAGE for the right. Contour lines are standardized mean differences (SMD). Stratification for nulli-RAGE and pro-RAGE cancers was done by color, as described in the legends of each panel. Non-exact p-values reported were plotted as stated in the manuscript (e.g., if p < 0.05, plotted p = 0.05; if p> 0.05, plotted p =0.5) as an estimate.

While these results indicate an association between AGE-RAGE axis activation or expression and cancer cell growth potential, a statistical assessment was needed due to the wide range of effects. A weighted Z-test was used to assess the statistical significance of FMA (*p*=4.8 x 10^−6^), pro-RAGE (*p*=2.95 x 10^−5^), and nulli-RAGE (*p*=2.12 x 10^−3^). Further weighted Z-tests assessing each individual organ of origin can be found in **Supplementary Table S7**. The overall results demonstrate that AGE-RAGE axis expression and activation encourage cancer cell growth in all cancer types, with a higher significance among pro-RAGE cancers.

### Effect of RAGE on cancer cell metastatic potential

Ninety-five (95) of the 160 cell culture studies evaluated the association between AGE-RAGE axis activation or expression and metastatic potential, as assessed by migration and invasion assays. Three hundred and twenty-two (322) out of 337 (95.54%) results using cancer cell lines, despite substantial variation in methodologies; treatment types; and doses among experiments, showed that increased AGE-RAGE axis activation or expression was associated with higher cancer cell metastasis potential. A majority of pro-RAGE results (269 of 277; 97.11%) and nulli-RAGE results (53 of 60; 88.33%) also reported that increased AGE-RAGE axis activation or expression resulted in increased cancer cell metastatic potential. In depth description of individual study results can be found in **Supplementary Table S3**.

To visually examine the data and provide an estimated standardizes effect of AGE-RAGE axis activation or expression on cancer cell metastatic potential, an albatross plot was generated (**Figure 5B**). The SMD range spanned –8.00 to 85.60, a wide range of effects. Thirty (30) results showed no effects of RAGE on cancer cell metastatic potential with an SMD below ±0.97. Forty one (41) results showed mild positive effects with an SMD between ±0.97 and ±1.94. Two results showed moderate negative effects, and 44 showed moderate positive effects with an SMD between ±1.94 and ±2.91. Six results showed strong negative effects and 214 results showed strong positive effects with an SMD greater than ±2.91.

Weighted Z-tests assessed the statistical significance of the implied association between AGE-RAGE axis activation or expression and cancer cell metastatic potential shown in the albatross plot. Clear significance was found in the FMA(*p*=7.62 x 10^−12^) and pro-RAGE (*p*=4.79 x 10^−15^) datasets and no significance was found in the nulli-RAGE (*p*=0.109) dataset. Each individual organ of origin was also assessed via weighted Z-tests (**Supplementary Table S7**). The overall results demonstrate that AGE-RAGE axis expression and activation encourage cancer cell metastatic potential in pro-RAGE cancers but not in nulli-RAGE cancers.

### Effect of RAGE on cancer cell apoptosis evasion

Thirty-two (32) of the 160 cell culture studies evaluated the association between AGE-RAGE axis activation or expression and apoptosis evasion, as assessed by molecular assays such as flow cytometry, Western blots, and luminescent assays. Sixty-two (62) of the 69 (89.86%) results using cancer cell lines reported increased AGE-RAGE axis activation or expression resulted in increased cancer cell apoptosis evasion, despite large variation in the experimental methodologies used. Ninety-four point fifty-four percent (94.54%; 52/55) pro-RAGE results and 71.43% (10/14) nulli-RAGE results also reported that increased AGE-RAGE axis activation or expression resulted in increased cancer cell apoptosis evasion. An in-depth description of individual study results can be found in **Supplementary Table S3**.

As above, an albatross plot (**Figure 5C**) was generated to integrate the data, and visual inspection of the plot provided an estimated standardized effect between AGE-RAGE axis activation or expression and cancer cell apoptosis evasion. The subsequent overall SMD range of –11.76 to 38.45 represents a wide range of effects. Three results showed no effects of RAGE on cancer cell apoptosis evasion with an SMD below ±0.97. Five results demonstrated mild positive effects with an SMD between ±0.97 and ±1.94. Twelve showed moderate positive effects with an SMD between ±1.94 and ±2.91. Six results showed strong negative effects, and 43 results showed strong positive effects with an SMD greater than ±2.91.

While these SMDs strongly indicate an positive association between AGE-RAGE axis activation or expression and cancer cell apoptosis evasion, a weighted Z-test had to be performed to assess the statistical significance of FMA (*p*=1.84 x 10^−14^), pro-RAGE (*p*=4.26 x 10^−13^), and nulli-RAGE (*p*=3.53 x 10^−5^). Further weighted Z-tests assessing each organ of origin can be found in **Supplementary Table S7**. The overall results demonstrate that AGE-RAGE axis expression and activation encourage cancer cell apoptosis evasion in all cancer types, with a much higher significance among pro-RAGE cancers.

### Effect of RAGE on cancer cell activated NF-κB expression

Of the 160 cell culture studies accepted, 32 studies evaluated the association between AGE-RAGE axis activation or expression and activated NF-κB expression, as assessed by luciferase reporters or Western blots. They reported, despite large variation in the methodologies utilized, 54 of the 60 (90%) positive correlations between AGE-RAGE axis activation or expression and increased cancer cell NF-κB expression. Forty-seven (47) of 53 (88.67%) pro-RAGE results and 7 of 7 (100%) nulli-RAGE results also reported that increased AGE-RAGE axis activation or expression resulted in increased cancer cell NF-κB expression (**Supplementary Table S3**).

An albatross plot (**Figure 5D**) was created to combine the data, and visual inspection of this plot suggested an estimated standardized effect linking AGE-RAGE axis activation or expression with cancer cell–activated NF-κB levels. With a span of –17.75 to 49.60, the overall SMD indicates a wide range of effects. Five results showed no effects of RAGE on cancer cell-activated NF-κB expression with an SMD below ±0.97. Two results showed mild negative effects, and nine results showed mild positive effects, with an SMD between ±0.97 and ±1.94. Seven showed moderate positive effects with an SMD between ±1.94 and ±2.91. Three results showed strong negative effects, and 34 results showed strong positive effects with an SMD greater than ±2.91.

While a positive association between AGE-RAGE axis activation or expression and cancer cell-activated NF-κB expression was strongly indicated by these results, a statistical assessment was still necessary to quantitatively assess significance. A weighted Z-test was used for the FMA (*p*=1.21 x 10^−5^), pro-RAGE (*p*=1.53 x 10^−4^), and nulli-RAGE (*p*=4.80 x 10^−4^) data sets. Further weighted Z-tests assessing each organ of origin can be found in **Supplementary Table S7**. The overall results show that AGE-RAGE axis expression and activation promote activated NF-κB expression across all cancer types.

### Effect of RAGE on in vivo tumor growth

Sixty-nine (69) of the 73 animal studies evaluated the association between AGE-RAGE axis activation or expression on tumor growth, as assessed by tumor volume, weight, bioluminescence, or count. One hundred and forty-three (143) out of 158 (90.51%) results, while varying in methodology, treatment, and doses, showed that increased activation or expression of the AGE-RAGE axis led to enhanced tumor growth. Most of pro-RAGE (125/132; 94.70%) and nulli-RAGE (18/26; 69.23%) results reported that increased AGE-RAGE axis activation or expression resulted in increased tumor growth. In depth description of individual study results can be found in **Supplementary Table S5**.

A comprehensive albatross plot (**Figure 6A**) was constructed to synthesize the data, and a visual examination of the plot yielded an estimated standardized effect size relating to AGE-RAGE axis activation or expression and tumor progression. The overall SMD range of –2.52 to 25.65 represents a wide range of effects. Twenty (20) results showed no effects of RAGE on tumor growth with an SMD below ±0.69. Seven results showed mild negative effects, and 29 results showed mild positive effects with an SMD between ±0.69 and ±1.38. Three results showed moderate negative effects and 33 showed moderate positive effects with an SMD between ±1.38 and ±2.07. One result showed strong negative effects and 65 results showed strong positive effects with an SMD greater than ±2.07.

**Figure 6.**
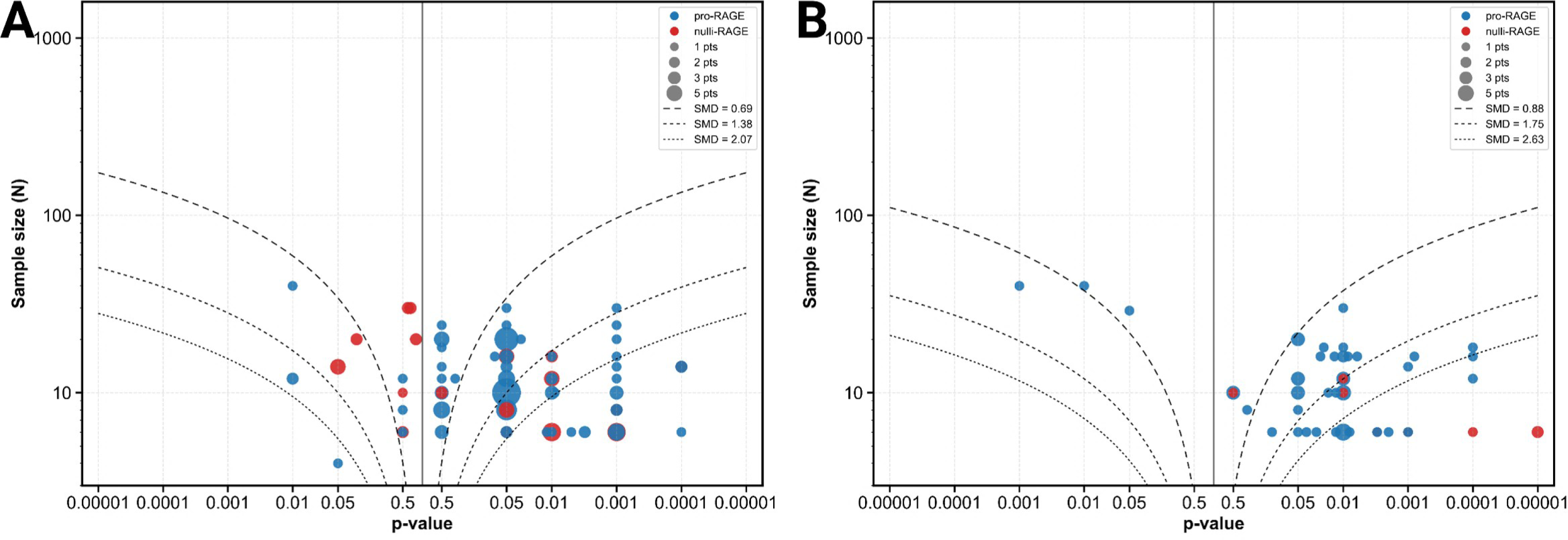
Albatross plot of animal studies measuring AGE-RAGE axis association and modulation with. (**A**) tumor growth, measured by tumor size, bioluminescence, or count; and (**B**) in vivo metastasis, measured by metastatic tumor burden or number of metastases. Each point represents data and its size is correlated to number of overlapping points, with the effect estimate (represented as a p-value), plotted against the total given sample size (n) included within each study. The directionality of the p-values correlate to negative associations with RAGE for the left and positive associations with RAGE for the right. Contour lines are standardized mean differences (SMD). Stratification for nulli-RAGE and pro-RAGE cancers was done by color, as described in the legends of each panel. Non-exact p-values reported were plotted as stated in the manuscript (e.g., if p < 0.05, plotted p = 0.05; if p> 0.05, plotted p =0.5) as an estimate.

These results strongly suggested an association between AGE-RAGE axis activation or expression and tumor growth, but a quantitative statistical assessment was missing. To address this, a weighted Z-test was employed to evaluate the statistical significance of FMA (p=1.66 x 10^−10^), pro-RAGE (*p*=9.59 x 10^−9^), and nulli-RAGE (*p*=1.60 x 10^−7^). Further weighted Z-tests assessing each individual organ of origin can be found in **Supplementary Table S8**. The overall results show that AGE-RAGE axis expression and activation promote tumor growth in all cancer types, with a higher significance in pro-RAGE cancers.

### Effect of RAGE on in vivo metastasis

The association between AGE-RAGE axis activation or expression and tumor metastasis was evaluated in 25 of the 73 animal studies, as assessed by metastatic tumor burden or number of metastases in relevant organ. Despite large variation in the methodologies used, the types of treatments performed, and the doses utilized, 59 of the 62 (95.16%) results using animal models reported increased AGE-RAGE axis activation or expression resulted in increased metastasis. Data showing that increased AGE-RAGE axis activation or expression resulted in increased metastasis was reported in 51 of 54 (94.44%) pro-RAGE results and 8 of 8 (100%) nulli-RAGE results. In depth description of individual study results can be found in Supplementary Table S5.

As performed in the other preclinical analyses, an albatross plot (**Figure 6B**) was generated and visually inspected to provide an estimated standardized effect between AGE-RAGE axis activation or expression and metastasis. Representing a wide range of effects, the overall SMD range was from –4.46 to 9.48. Nine results showed no effects of RAGE on metastasis with an SMD below ±0.88. 14 results showed mild positive effects with an SMD between ±0.88 and ±1.75. One result showed moderate negative effects and eight showed moderate positive effects with an SMD between ±1.75 and ±2.63. Two results showed strong negative effects and 28 results showed strong positive effects with an SMD greater than ±2.63.

A positive association between AGE-RAGE axis activation or expression and metastasis was indicated by these results. A weighted Z-test was used to quantitatively assess the statistical significance of FMA (*p*=1.46 x 10^−13^), pro-RAGE (*p*=1.09 x 10^−11^), and nulli-RAGE (*p*=2.42 x 10^−8^). Further weighted Z-tests assessing each individual organ of origin can be found in **Supplementary Table S8**. The overall results demonstrate that AGE-RAGE axis expression and activation encourages metastasis in all cancer types, with a higher significance in pro-RAGE cancers.

### Quantitative assessments of RAGE expression in clinical samples

Twenty-six (26) clinical studies quantitatively assessed RAGE expression between cancer samples and adjacent normal tissue, and 4 clinical studies did the same between high-grade and low-grade cancerous tissue, all as assessed by molecular analyses such as Western blots and ELISAs. Twenty-two (22) of the 37 (59.46%) cancer vs. adjacent normal tissue results reported increased RAGE expression in cancerous tissue, and 5 of the 6 (83.33%) high-grade vs. low-grade results reported increased RAGE expression in high-grade cancerous tissue. Fifteen (15) of 16 (93.75%) pro-RAGE results and 7 of 21 (33.33%) nulli-RAGE results also reported increased RAGE expression in cancerous tissue. In addition, 5 of 5 (100%) pro-RAGE results and 0 of 1 (0%) nulli-RAGE results reported increased RAGE expression in high-grade cancerous tissue. In depth description of individual study results can be found in **Supplementary Table S1**.

To assess these non-case-control clinical datasets, albatross plots (**Supplementary Figure S11**) were generated and used to provide an estimated standardized effect between RAGE expression and clinical cancer vs. adjacent normal tissue and between RAGE expression and clinical cancer grading. For clinical cancer samples, the overall SMD range of –36.71 to 15.95 represents a wide range of effects. One result showed mild positive effects of RAGE expression on clinical cancer samples with an SMD between ±0.22 and ±0.43. One result showed moderate positive effects with an SMD between ±0.43 and ±0.65. 15 results showed strong negative effects and 20 results showed strong positive effects with an SMD greater than ±0.65.

A quantitative statistical assessment was performed to assess the estimated positive association between RAGE expression and clinical cancer samples. Weighted Z-tests were performed on the FMA (*p*=0.620), pro-RAGE (*p*=1.60 x 10^−5^), and nulli-RAGE (*p*=0.046; negative association) datasets. Further weighted Z-tests assessing each individual organ of origin can be found in **Supplementary Table S9**. The overall results demonstrate that RAGE expression was significantly positively associated with clinical pro-RAGE cancer samples and significantly negatively associated with clinical nulli-RAGE cancer samples.

For clinical cancer grading, the overall SMD range of –0.66 to 35.43 represents a wide range of effects. One result showed no effects of RAGE expression on clinical cancer grading with an SMD below ±0.10. One result showed moderate positive effects with an SMD between ±0.25 and ±0.50. One result showed strong negative effects and three results showed strong positive effects with an SMD greater than ±0.50.

While these results indicate an association between RAGE expression and clinical cancer grading, a statistical assessment was needed due to the wide range of effects. Weighted Z-tests were used to assess statistical significance of FMA (*p*=0.001), pro-RAGE (*p*=9.96 x 10^−4^), and nulli-RAGE (*p*=0.500). Further weighted Z-tests assessing each individual organ of origin can be found in **Supplementary Table S9**. The overall results demonstrate that RAGE expression was significantly associated only with clinical pro-RAGE cancer grading.

## Discussion

In this systematic review and meta-analysis of RAGE in clinical cancer samples, we demonstrated that there is a high prevalence of RAGE expression in cancerous tissues compared with adjacent normal tissue, and that RAGE can be used as a biomarker to differentiate between high– and low-grade cancers and between cancers that do or do not invade regional lymph nodes. Furthermore, we identified that gastrointestinal and respiratory system cancers, designated as nulli-RAGE, appear to either have null or negative associations between RAGE expression and cancerous tissue; cancer grade; regional lymph node invasion; and patient survival. The remaining cancers, designated as pro-RAGE, had greater associations in all measured outcomes, including patient survival, where high prevalence of RAGE expression was linked with a higher likelihood for patient mortality.

We further expanded this systematic review to include preclinical studies that investigated the association between RAGE and pro-tumorigenic effects in cancer. While no consistent strategies currently exist to perform meta-analyses in preclinical studies that have used various unique methodologies, we adapted the guidelines from the WCRF/UoB recommendations to perform a pseudo meta-analysis of these data^17^. Critically, results from this analysis show consistently strong positive associations between RAGE and cancer growth potential, metastatic potential, apoptosis evasion, activated NF-κB expression, tumor growth, and metastasis across all evaluated studies.

Our results demonstrate for the first time that RAGE expression is elevated in most cancers, with an overall OR of 3.49 when compared to adjacent normal tissue, with a strong elevation in pro-RAGE cancers (OR: 8.54) and no significant change in nulli-RAGE cancers (OR: 0.60). This expands upon our previous work in assessing RAGE as a novel biomarker for prostate cancer and found that RAGE expression was elevated when compared to benign prostatic hyperplasia (OR: 11.33) and in high-grade cancer when compared to low-grade prostate tumors (OR: 2.50)^14^. Our results also showed that high-grade and regional lymph node-invading pro-RAGE cancers were much more likely to express RAGE compared with low-grade and non-lymph node-invading pro-RAGE cancers and all nulli-RAGE cancer clinical outcomes, suggesting RAGE could be used as a biomarker to differentiate between gradations of cancer and to assess metastatic likelihood.

RAGE expression appears to be well-situated to serve as a biomarker to assess when indolent, localized pro-RAGE cancers undergo a phenotypic switch to aggressive, high-grade, invasive cancers and pose a risk to patient survival. This is supported by the body of evidence identified in preclinical cancer cell and animal studies that indicates RAGE’s significant role in promoting cancer growth, metastasis, evasion of apoptosis, and activation of the key inflammatory marker, NF-κB. Cell culture studies strongly linked AGE-RAGE axis expression and activation with cancer cell growth potential, apoptosis evasion, and activated NF-κB expression in all cancer types and metastatic potential in pro-RAGE cancers. This AGE-RAGE axis expression and activation was assessed through a variety of methods, such as activation with RAGE ligands such as HMGB1 and AGEs; transfection with full length-RAGE, truncated RAGE isoforms, or over-expressing variants; inhibition via RAGE-targeted pharmacological agents such as FPS-ZM1 or TTP488; or direct RAGE inhibition through antibodies, siRNA, or other methods^83,135,175,184,185,193,197^.

Animal studies consistently demonstrated that expression and activation of the AGE-RAGE axis was directly linked to tumor growth and metastasis. This was also evaluated using a wide range of methods, such as implantation in RAGE-null mice; treatment with anti-AGE-RAGE axis therapies such as glycyrrhizin, ALT-711, or RP7; modulation of AGEs such as S100A4, HMGB1, or hi-AGE diet; or genetic modulation of RAGE expression through transfection or silencing^32,96,152,234,239,245,246,253^.

These preclinical results support the body of literature that shows that the AGE-RAGE axis’ signaling pathways such as PI3K/Akt and NF-κB support an inflammatory microenvironment which in turn may induce tumorigenesis, support tumor advancement, and initiate therapy resistance^7–9,255^. Furthermore, the results of this study indicate that the AGE-RAGE axis engages with various hallmarks and enabling characteristics of cancer, namely, “resisting cell death” through increasing apoptosis evasion; “sustaining proliferative signaling” through increasing growth potential and tumor growth; “activating invasion and metastasis” through increasing metastatic potential *in vitro* and *in vivo*; and “tumor-promoting inflammation” through increasing NF-κB activation^11,256,257^.

Due to the high association identified between RAGE and cancerous tissue, RAGE expression could be used as a prognostic tool to identify and monitor localized tumors. Longitudinal monitoring of RAGE expression via repeated biopsies are not clinically feasible, as exampled by prostate cancer biopsies which which can be painful and has significant risk of false-negatives^258^. To address this issue, our research group has developed a multimodal imaging strategy to non-invasively quantify RAGE expression in tissues, through the use of a RAGE-targeted nanoparticle or RAGE-antagonistic peptide probe, which has demonstrated consistent utility in imaging RAGE in prostate cancer at various stages of its progression.^12,13^. This imaging platform has the potential to impact the current diagnostic and therapeutic landscape of prostate and breast cancer treatment, and perhaps other solid tumors, by enabling clinicians to use PET-CT or optical imaging tools to non-invasively and longitudinally monitor tumors. Increases in RAGE expression over time would indicate cancer progression and an increase in metastatic potential, providing a necessary criterion that will help determine clinical therapeutic response.

Strengths of this study include its novelty in comprehensively analyzing all available cancer studies involving RAGE expression and activation and in identifying RAGE as a potent biomarker for cancer outcomes, particularly in pro-RAGE cancers, with these associations remaining strong following leave-one-out sensitivity analysis. OR values for pro-RAGE cancerous tissue remained above 7.0 in all scenarios, demonstrating a robust association between RAGE expression and cancerous tissues. OR values for pro-RAGE cancer grading remained above 1.9 in all scenarios, demonstrating an association between RAGE expression and high-grade cancer. OR values for regional lymph node invasion in pro-RAGE cancers remained above 4.2 in all scenarios, demonstrating a robust association between RAGE expression and regional lymph node invasion. OR values for pro-RAGE cancer patient survival remained below 0.5 in all scenarios, demonstrating a negative association between RAGE expression and patient survival. Bias and sensitivity analyses were all supportive of these results, which lends confidence to the strength of the associations between RAGE expression and pro-RAGE cancer.

Limitations of this study include high complexity and high intra-study heterogeneity due to the small number of accepted clinical studies for each organ of origins with most having below three reports for all outcomes, and only nine unique reports for patient survival. Only liver and prostate in clinical tumor grading and colorectal, gastric, kidney, and nasoesophageal in clinical tumor grading and lymph invasion have 3 or more reports. Small-study effects, as measured by Harbord’s test, were seen in all patient survival analyses and in the nulli-RAGE cancerous tissue analysis. However, random-effects models were run to account for this heterogeneity when examining both clinical and preclinical outcomes. Although preclinical studies demonstrated consistently positive and robust associations with AGE-RAGE activation and expression and preclinical outcomes, albatross plots can only be visually used as an estimation of the overall but not the absolute effect. Despite this, the fact that most examined studies showed positive associations with preclinical outcomes demonstrates the high potential for RAGE to serve as a biomarker that potently contributes to pro-RAGE tumorigenesis.

In addition, results from this systematic review and meta-analysis are clinically meaningful, as dietary modification to minimize intake of dietary AGEs may provide cancer patients with a simple lifestyle-change adjuvant treatment to existing therapies. Dietary reduction of AGEs has been found to reduce AGE-RAGE axis activation and expression and induce significant changes in the outcomes of other inflammatory-related disease states such as chronic kidney disease and diabetes^259,260^. As such, results from this meta-analysis demonstrating the effects of RAGE expression and AGE-RAGE axis activation on cancer can be immediately translatable to clinical practice. Further work will be needed to understand how existing and future targeted anti-RAGE axis therapies such as Alagebrium and Azeliragon, both of which have been investigated in clinical trials for other disease states and were found to be safe but lacked efficacy in treating cardiovascular or Alzheimer’s disease respectively, may work to prevent cancer progression in both preclinical and clinical models, which our team is actively pursuing^261–263^.

## Conclusions

This systematic review and meta-analysis of clinical and preclinical studies demonstrated a robustly positive association between RAGE and cancer outcomes. The results of this study provide novel insight into a potentially potent biomarker for cancer, immediately actionable evidence for clinical recommendations for dietary modification to improve cancer outcomes, and new avenues for adjuvant cancer therapies.

## List of Abbreviations

AGEs: Advanced Glycation End-products
RAGE: Receptor for Advanced Glycation End-products
EMT: Epithelial-mesenchymal transition
HMGB1: High Mobility Group Box 1 Protein
DAMP: Damage-associated molecular pattern
WCRF: World Cancer Research Fund
UoB: University of Bristol
OR: Odds ratio
CI: Confidence intervals
QA: Quality Assessment
SD: Standard deviation
SEM: Standard error of the mean
SYRCLE: SYstematic Review Centre for Laboratory animal Experimentation
nulli-RAGE: Cancer types with negative or null associations with RAGE expression
pro-RAGE: Cancer types with positive associations with RAGE expression
FMA: Full meta-analysis

## Footnotes

A 32-34,45,47,51,53,56,66,69,70,73,74,76-78,81-224

B 32,45,57,59,81,83,96,110,113,116,119,122,139,149,150,152,160,164,170-172,179,180,183,184,186,188,189,192,196,197,200,203, 205,207,208,210,213-217,220,225-254

C 29-32,35-40,42-44,46,48-50,54-56,58-60,63,64,67-72,74-80

D 29,33-35,37,38,40-42,45,50,51,53,55-58,61,62,65-68,72-74,78-80

E 33-35,42,45,49-51,53,54,65-68,72-74,79,80

F 47,51,52,54,59,65,67,68,80

G 45,47,53,56,66,69,70,73,74,76,78,81-87,89,90,92-96,99,101-105,107,108,110,111,113-115,117-124,126,128-131,136-138,140, 141,143-147,149-154,156-159,161,163-179,181,182,184,185,187,188,190-198,200-205,207,208,210-213,215-217,219,221-224

H 32-34,51,53,56,69,70,73,74,77,81,84,85,88,91,92,95-100,103,105-109,111,112,115-117,119,121,122,125,129-135,138-140,142, 143,145,146,148-153,155-158,160-162,164,165,169,170,173,175,176,178,180,183,184,186,187,191,192,195,197,199-201,205,212-215,218-220,222,223

I 32,69,73,81,86,87,89,94,101,103,104,106,127,130,131,136,149,151,156,157,166,167,171,177,188,205,206,209,211,212,222,223

J 70,76,81,96,132,140,143,150,153,155,161,162,176,178,189,191,193,197,198,206,207,209,210,218

K 32,45,57,59,81,83,96,110,113,116,119,122,139,149,150,152,160,164,170-172,179,180,184,188,189,192,196,197,200,203,205,207, 208,210,213-217,220,225-233,235-249,251-254

L 59,81,116,119,139,150,164,170,180,183,184,186,188,189,197,205,210,229,230,233,234,243,245,250,252

M 33,51,53,56,69,70,73,76,78,82-84,86-88,90-92,94,97-99,102,105-108,110,111,115,119,121-125,127-130,132,134-136,138,139, 142,143,145-148,151,153,154,157-160,163,165-171,174,176-179,181,183,184,191,193-196,198-200,202,204,210,212,213,217,218, 220,221

N 32,34,45,47,66,71,74,77,81,85,89,93,95,96,100,101,103,104,109,112-114,116-118,120,126,131,133,137,140,141,144,149,150,152, 155,156,161,162,164,172,173,175,180,182,185-190,192,197,201,203,205-209,211,215,216,219,222,224

O 32,45,57,59,81,83,96,110,113,116,119,122,139,149,150,152,160,164,170-172,179,180,183,184,186,188,189,192,196,197,200,203, 205,207,208,210,213,215-217,220,225-235,237,238,240-245,248,250,253,254

