## Supplemental Table Legends for "The Role of the Receptor for Advanced Glycation End-Products in Cancer: Evidence from a Systematic Review and Meta-Analysis"

**Supplementary Table S1.** Characteristics of clinical studies included in the meta-analysis.

**Supplementary Table S2.** Quality assessment of included clinical studies.

**Supplementary Table S3.** Characteristics of cell culture studies included in the meta-analysis.

**Supplementary Table S4.** Quality assessment of included cell culture studies (adapted from WCCRF/UoB recommendations) [17].

**Supplementary Table S5.** Characteristics of animal studies included in the meta-analysis.

**Supplementary Table S6.** Quality assessment of included animal studies (SYRCLE tool) [21].

**Supplementary Table S7.** Table of cell culture studies *p*-values. *p*-values are assumed for positive association with exception for *negative or †null association between AGE-RAGE axis modulation and cancer cell outcomes.

**Supplementary Table S8.** Table of animal studies *p*-values. *p*-values are assumed for positive association with exception for *negative or †null association between AGE-RAGE axis modulation and animal cancer outcomes.

**Supplementary Table S9.** Table of clinical studies *p*-values. *p*-values are assumed for positive association with exception for *negative or †null association between RAGE expression and clinical cancer outcomes.
