## Supplemental Table 1 for "The Role of the Receptor for Advanced Glycation End-Products in Cancer: Evidence from a Systematic Review and Meta-Analysis"

| Author, Year | Organ of Origin | Control | Subjects (n) | Method for RAGE | OR (95% CI)[1] | SMD (SE)[2] | *p*-value[3] |
|---|---|---|---|---|---|---|---|
| Aboushousha, 2018 [29] | Liver | Non-tumor, G1 | Non-tumor=36, HCC G1=10 HCC G2+3=23 | IHC | C/C: 25.00 (6.75-92.59); Histo: 0.25 (0.03-2.41) | N/A | **<0.0001; 0.3820** |
| Akkus, 2021 [30] | Prostate | BPH | BPH=40, Localised PCa = 46 | IHC | N/A | C/C: 0.6138 ± 0.2214 | **<0.010** |
| Amornsupak, 2017 [31] | Breast | Benign Lesions | benign=31, tumor=96 | IHC | C/C: 3.57 (1.32-10.68) | N/A | **0.0066** |
| Dai, 2023 [32] | Breast | non-TNBC | non-TNBC=42, TNBC=42 | RNA-seq | N/A | C/C: 0.2251 ± 0.2189 | **0.001** |
| Deng, 2017 [33] | Gastric | Well diff., N0 | Well Diff=69, Poor Diff=91, N0=70, N1=90 | IF | Histo: 1.14 (0.60-2.17); Mets: 2.09 (1.09-3.99) | N/A | **0.679; 0.025** |
| Deng, 2017 [34] | Colorectal | Well diff., N0 | Well Diff=69, Poor Diff=91, N0=70, N1=90 | IF | Histo: 1.14 (0.60-2.17); Mets: 2.09 (1.09-3.99) | N/A | **0.679; 0.025** |
| Guo, 2015 [35] | Kidney | G1, N0, tumor edge | G1=6, G2-4=74, N0=64, N1-2=16, tumor edge=7, tumor=7 | IF; Flow Cytometry | Histo: 2.77 (0.48-16.11); Mets: 2.82 (0.82-9.67) | C/C: 9.2676 ± 2.0902 | **0.3959; 0.1576; <0.010** |
| Guo, 2019 [36] | Kidney | PANT | PANT=12, tumor=12 | qRT-PCR | N/A | C/C: 0.9771 ± 0.4358 | **<0.05** |
| Hermani, 2005 [37 | Prostate | BPH, LG | BPH=48, tumor=75, LG=24, HG=32 | IHC | C/C: 40.89 (9.20-181.82); Histo: 2.56 (0.84-7.76) | N/A | **<0.0001; 0.1059** |
| Hiwatashi, 2008 [38] | Liver | Normal Liver, Well diff. | Liver=6, HCC=36, Well Diff=10, Poor Diff=38 | IHC, qRT-PCR | Histo: 0.48 (0.11-2.12) | C/C: 1.2828 ± 0.4645 | **0.4780; <0.01** |
| Hofmann, 2004 [39] | Lung | PANT | PANT=15, tumor=43 | RT-PCR | N/A | C/C: -4.6819 ± 0.5394 | **<0.0001** |
| Ishiguro, 2005 [40] | Prostate | PANT, tumor, Well diff. | PANT= 43, tumor=43, refractory=13, Well diff.=8, Poor diff.=14 | qRT-PCR | N/A | C/C: 0.3023 ± 0.2169; Histo: 0.7726 ± 0.3254; Histo: -0.6586 ± 0.4562 | **<0.05; ns; ns** |
| Ito, 2014 [41] | Liver | Well diff. | Well diff.=13, Poor diff.=52 | IHC | Histo: 0.37 (0.07-1.88) | N/A | **0.3146** |
| Jing, 2008 [42] | Lung | Well diff., N0, PANT | Well Diff=20, Poor Diff=12, N0=13, N1=19; tumor=12, PANT=12 | IHC, RTFQ-PCR | Histo: 0.37 (0.06-2.19); Mets: 0.14 (0.02-0.85) | C/C: -1.1794 ± 0.4479; - 8.0727 ± 1.3249 | **0.4224; 0.0383; 0.025; <0.05** |
| Jing, 2008 [42] | Esophageal | Well diff., N0, PANT | Well Diff=26, Poor Diff=3, N0=9,N1=20; tumor=12, PANT=12 | IHC, RTFQ-PCR | Histo: 1.06 (0.08-13.33); Mets: 0.93 (0.18-4.90) | C/C: 1.1950 ± 0.4489; 1.4478 ± 0.4667 | **1.000; 1.000; 0.009; <0.05** |
| Jing, 2010 [43] | Lung | PANT | PANT=28, tumor=28 | IHC | N/A | C/C: -1.7795 ± 0.3187 | **0.022** |
| Jing, 2010 [44] | Lung | PANT | PANT=14, tumor =14 | Western Blotting | N/A | C/C: -36.7101 ± 5.2571 | **<0.05** |
| Jing, 2010 [44] | Esophageal | PANT | PANT=18, tumor=18 | Western Blotting | N/A | C/C: 15.9481 ± 2.0058 | **<0.05** |
| Khoo, 2023 [45] | Prostate | Gleason<7, N0 | G<7=22, G>7=19, N0=56, N1=16 | IHC | Histo: 4.57 (1.22-17.16); Mets: 1.91 (0.59-6.21) | N/A | **0.0296; 0.3925** |

| | | | | | | | |
|---|---|---|---|---|---|---|---|
| Khorramdelazad, 2015 [46] | Bladder | PANT | PANT=17, tumor=17 | qRT-PCR | N/A | C/C: 3.0865 ± 0.5232 | **<0.05** |
| Kobayashi, 2007 [47] | Lung | Dead | Dead=105, Living=77 | qRT-PCR | Survival: 1.50 (0.82-2.75) | N/A | **0.2225** |
| Korwar, 2012 [48] | Breast | PANT | PANT=2, tumor=7 | Western Blot, ELISA | N/A | C/C: 10.3478 ± 3.2129; 2.0211 ± 1.0060 | **<0.005; <0.05** |
| Kuniyasu, 2003 [49] | Colorectal | non-neoplastic mucosa, Duke's B | mucosa=3, tumor=119, Duke's B=54, Duke's C=47 | IHC | C/C: 8.70 (0.44-172.16); Mets: 18.58 (6.84-50.48) | N/A | **0.3295; <0.0001** |
| Landesberg, 2008 [55] | Oral | PANT, tumor, Well-moderately diff. | PANT=12, tumor=38, well-moderately diff=14, mod.-poor diff=24 | IHC | C/C: 0.03 (0-0.59); Histo: 0.02 (0-0.15) | N/A | **0.0032; <0.0001** |
| Li, 2022 [50] | Esophageal | PANT, Well diff., N0 | PANT=80, tumor=80, well diff=18, mod.-poor diff=62, N0=26, N1=34 | IHC | C/C: 2.15 (1.14-4.05); Histo: 0.72 (0.25-2.11); Mets: 0.85 (0.33-2.19) | N/A | **0.026; 0.6007; 0.8128** |
| Li, 2024 [51] | Head and Neck | Gx/G0, N0, Dead | Gx/G0=66, G1/G2=42, N0=88, N1=20, Dead=25, Alive=83 | IHC | Histo: 1.15 (0.48-2.75); Mets: 8.36 (2.88-24.28); Survival: 0.22 (0.08-0.57) | N/A | **0.8249; 0.0001; 0.0036** |
| Liang, 2025 [52] | Kidney | Dead | Dead=490, Living=40 | RNA seq | Survival: 0.40 (0.20-0.81) | N/A | **0.0128** |
| Lin, 2012 [53] | Kidney | G1, N0 | G1=10, G2-4=110, N0=115, N1=5 | IHC | Histo: 2.17 (0.58-8.12); Mets: 8.78 (0.47-162.54) | N/A | **0.3209; 0.2352** |
| Lin, 2022 [54] | Lung | Normal Lung, N0, Dead | Normal Lung=111, tumor=111, N0=348, N1-3=171, Dead=384, Alive=94 | Unknown for C/C, IHC | Mets: 0.81 (0.56-1.17); Survival 2.71 (1.67-4.38) | C/C: -6.5813 ± 0.3422 | **0.3043; <0.0001; <0.0001** |
| Medapati, 2015 [56] | Thyroid | PANT, PTC+FTC | PANT=9, tumor=58, PTC+FTC=48, UTC=10 | IHC | C/C: 64.04 (3.50-1173.13); Histo 0.61 (0.13-2.81) | N/A | **0.0001; 0.5778** |
| Muoio, 2023 [57] | Breast | Stage I | Stage I=475, Stage II/III=915 | RAGE mRNA Expression | N/A | Histo: 0.0062 ± 0.0565 | **<0.001** |
| Nankali, 2016 [58] | Breast | PANT, I/II TNM, | PANT=20, tumor=25, TNM I/II=17, TNM >II=8 | RT-PCR | N/A | C/C: -1.5365 ± 0.3441; Histo: 35.4275 ± 5.4185 | **0.045; <0.001** |
| Nasser, 2015 [59] | Breast | Normal Breast, Dead | Normal Breast=10, tumor=50, Dead=153, Alive=355 | RAGE expression from microarrays | Survival: 0.60 (0.38-0.96) | C/C: 1.907 ± 0.3901 | **0.0345; <0.05** |
| Qian, 2019 [66] | Colorectal | Well Diff, N0 | Well Diff=32, Mod.-Poor=34, N0=34, N1=32 | IHC | Histo: 6.19 (2.02-18.97); Mets 4.18 (1.45-12.02) | N/A | **0.0011; 0.0112** |
| Qie, 2015 [67] | Kidney | PANT, High Diff., N0, Dead | PANT=80, tumor=80, high diff.=31, medium/low diff.=49, N0=38, N1=42, Dead=77, Alive=3 | IHC | C/C: 17.67 (7.44-41.96); Histo: 14.00 (1.63-120.41); Mets: 9.26 (1.08-79.21); Survival: 0.20 (0.02-2.49) | N/A | **<0.0001; 0.0047; 0.0240; 0.2741** |

| | | | | | | | |
|---|---|---|---|---|---|---|---|
| Rahimi, 2017 [60] | Ovaries | PANT | PANT=222, tumor=222 | qRT-PCR | N/A | C/C: 1.2629 ± 0.1040 | **<0.001** |
| Sasahira, 2005 [61] | Colorectal | Mild Atypia | Mild=36, Moderate-Severe=60 | IHC | Histo: 19.43 (4.28-88.26) | N/A | **<0.0001** |
| Sasahira, 2007 [62] | Oral | TNM I/II | TNM I/II=9, TNM III/IV=11 | ELISA | N/A | Histo: 0.9563 ± 0.4793 | **ns** |
| Sinduja, 2024 [63] | Oral | Submucous fibrosis | Submucous fibrosis=16, tumor=17 | ELISA | N/A | C/C: -1.8932 ± 0.4267 | **<0.05** |
| Tafani, 2010 [64] | Breast | Normal Breast | Normal Breast=8, tumor=8 | Western Blot | N/A | C/C: 5.8056 ± 1.2636 | **<0.05** |
| Tateno, 2009 [65] | Esophageal | Well diff., N0, Dead | Well diff.=73, Mod.-Poor diff.=135, N0=72, N1=136, Dead=121, Alive=87 | IHC | Histo: 1.46 (0.83-2.60); Mets: 0.88 (0.50-1.55); Survival: 2.32 (1.32-4.09) | N/A | **0.2451; 0.7741; 0.0048** |
| Wang, 2015 [68] | Gastric | PANT, well diff., N0, Dead | PANT=69, tumor=180, well diff.=18, mod.-poor=162, N0=46, N1=134, Dead=105, Alive=75; PANT=30, tumor=30 | IHC, qRT-PCR | C/C: 6.32 (3.27-12.22); Histo: 4.92 (1.67-14.51); Mets: 2.16 (1.09-4.26); Survival: 0.50 (0.27-0.92) | C/C: 0.9320 ± 0.2727 | **<0.001** |
| Wang, 2020 [69] | Lung | Normal Lung | Normal Lung=26, tumor=24 | RAGE expression from dataset | N/A | C/C: -5.3191 ± 0.6199 | **<0.01** |
| Wang, 2021 [70] | Nasopharynx | Benign Inflammation | Benign=15, tumor=15 | Western Blotting | N/A | C/C: 0.8718 ± 0.3843 | **<0.05** |
| Wu, 2018 [71] | Lung | PANT | PANT=194; tumor=527 | RAGE expression from dataset | N/A | C/C: -6.057 ± 0.4368; -9.9753 ± 0.4886; -11.7293 ± 0.8805; -6.0740 ± 0.3427 | **<0.00001; <0.00001; <0.00001; <0.00001** |
| Xu, 2013 [72] | Gastric | T1, N0 | T1=7, T2-3=33, N0=16, N1=24 | IHC | Histo: 0.33 (0.04-3.12); Mets: 5.00 (1.17-21.39) | N/A | **0.6521; 0.0367** |
| Xu, 2013 [73] | Gastric | PANT, T1, N0 | PANT=40, tumor=40, T1=7, T2-3=33, N0=40, N1=40 | IHC | C/C: 0.16 (0.06-0.44); Histo: 2.08 (0.35-12.32); Mets: 5.00 (1.17-21.39) | N/A | **0.0004; 0.6770; 0.0722** |
| Yang, 2015 [74] | Liver | PANT, Stage 1/2, N0 | PANT=75, tumor=75, S1/2=52, S3-4=23, N0=52, N1=23 | IHC | C/C: 3.56 (1.81-6.97); Histo: 4.89 (1.29-18.53); Mets: 8.33 (1.77-39.24) | N/A | **0.0003; 0.0166; 0.003** |
| Yang, 2023 [75] | Lung | Normal Lung | Normal Lung=59, LUAD=515, Normal Lung=51, LUSC=501 | RAGE mRNA expression from dataset | N/A | C/C: -6.2525 ± 0.2305; -7.3397 ± 0.2659 | **<0.001; <0.001** |
| Yaser, 2012 [76] | Liver | Para-neoplastic | Para-neopl. = 10, neopl.=10, para-neopl.=5, neopl.=5 | qRT-PCR, Western Blot | N/A | C/C: 6.7723 ± 1.2605; 3.1631 ± 1.0801 | **<0.01; <0.01** |
| Yin, 2018 [77] | Breast | PANT | PANT=5, tumor=5 | Western Blot | N/A | C/C: 7.6596 ± 2.2124 | **<0.0001** |

| | | | | | | | |
|---|---|---|---|---|---|---|---|
| Zhang, 2016 [78 | Bone | Peritumor, Stage 1/2 | Peritumor=65, tumor=65, peritumor=27, tumor=27, Stage 1/2=34, Stage 3=31, | IHC, Western Blot | C/C: 3.01 (1.46-6.20); Histo: 3.64 (1.12-11.80) | C/C: 3.8821 ± 0.4725 | **0.0043; 0.0321; <0.001** |
| Zhao, 2014 [79] | Prostate | BPH, Gleason<7, N0 | BPH=30, tumor=85, Gleason<7=28, Gleason>7=22, N0=47, N1=38 | IHC | C/C: 4.25 (1.75-10.32); Histo: 4.00 (0.75-21.22); Mets: 5.47 (1.45-20.66) | N/A | **0.002; 0.1535; 0.0079** |
| Zhou, 2014 [80] | Nasopharynx | Chronic nasopharyngitis, Stage 1/2, N0, Dead | Chronic Nasopharyngitis=60, tumor=30, Stage 1/2=22, Stage 3/4=38, N0=30, N1=30, Dead= 30, Alive=30 | IHC | C/C: 0.30 (0.12-0.75); Histo: 0.64 (0.19-2.14) ; Mets: 3.33 (1.00-11.14); Survival: 0.19 (0.05-0.74) | N/A | **0.0117; 0.5599; 0.0840; 0.0194** |

1 Odds ratio (OR) and 95% confidence interval (CI) calculated from case-control outcomes

2 Standardized Mean Difference (SMD) and Standard Error (SE)

3 Bolded p-values indicate significance (p<0.05)

Abbreviations: receptor for advanced glycation end-products (RAGE), hepatocellular carcinoma (HCC), immunohistochemistry (IHC), benign prostatic hyperplasia (BPH), prostate cancer (PCa), triple-negative breast cancer (TNBC), immunofluorescence (IF), paired adjacent normal tissue (PANT), low grade (LG), high grade (HG), quantitative reverse transcription polymerase chain reaction (qRT-PCR), real-time fluorescent quantitative polymerase chain reaction (RTFQ-PCR), enzyme-linked immunoabsorbent assay (ELISA), papillary thyroid cancer (PTC), follicular thyroid cancer (FTC), anaplastic/undifferentiated thyroid cancer (UTC), lung adenocarcinoma (LUAD), lung squamous cell carcinoma (LUSC)
