## Supplemental Table 2 for "The Role of the Receptor for Advanced Glycation End-Products in Cancer: Evidence from a Systematic Review and Meta-Analysis"

| Source Author, Year | Selection | | | | Comparability[5] | Exposure | | | Total[9] |
|---|---|---|---|---|---|---|---|---|---|
| | Definition[1] | Representative[2] | Selection[3] | Definition[4] | | Ascertainment[6] | Method[7] | Technique[8] | |
| Aboushousha, 2018 [29] | 1 | 1 | 1 | 1 | 1 | 1 | 1 | 1 | 8 |
| Akkus, 2021 [30] | 1 | 1 | 1 | 0 | 1 | 1 | 1 | 1 | 7 |
| Amornsupak, 2017 [31] | 1 | 1 | 1 | 0 | 1 | 1 | 1 | 1 | 7 |
| Dai, 2023 [32] | 1 | 0 | 1 | 0 | 0 | 0 | 0 | 1 | 3 |
| Deng, 2017 [33] | 1 | 1 | 1 | 0 | 0 | 1 | 1 | 1 | 6 |
| Deng, 2017 [34] | 1 | 1 | 1 | 1 | 0 | 1 | 1 | 1 | 7 |
| Guo, 2015 [35] | 1 | 1 | 1 | 0 | 0 | 1 | 1 | 1 | 6 |
| Guo, 2019 [36] | 1 | 1 | 1 | 1 | 0 | 1 | 1 | 1 | 7 |
| Hermani, 2005 [37] | 1 | 0 | 1 | 0 | 0 | 1 | 0 | 1 | 4 |
| Hiwatashi, 2008 [38] | 1 | 1 | 1 | 1 | 1 | 1 | 1 | 1 | 8 |
| Hofmann, 2004 [39] | 1 | 1 | 1 | 1 | 0 | 1 | 1 | 1 | 7 |
| Ishiguro, 2005 [40] | 1 | 1 | 1 | 1 | 0 | 1 | 1 | 1 | 7 |
| Ito, 2014 [41] | 1 | 1 | 1 | 1 | 1 | 1 | 1 | 1 | 8 |
| Jing, 2008 [42] | 1 | 1 | 1 | 1 | 0 | 1 | 1 | 1 | 7 |
| Jing, 2010 [43] | 1 | 1 | 1 | 1 | 0 | 1 | 1 | 1 | 7 |
| Jing, 2010 [44] | 1 | 1 | 1 | 1 | 0 | 1 | 1 | 1 | 7 |
| Khoo, 2023 [45] | 1 | 1 | 1 | 1 | 0 | 1 | 1 | 1 | 7 |
| Khorramdelazad, 2015 [46] | 1 | 1 | 1 | 1 | 0 | 1 | 1 | 1 | 7 |
| Kobayashi, 2007 [47] | 1 | 1 | 1 | 1 | 0 | 1 | 1 | 1 | 7 |
| Korwar, 2012 [48] | 1 | 1 | 1 | 1 | 0 | 1 | 1 | 1 | 7 |
| Kuniyasu, 2003 [49] | 1 | 1 | 1 | 1 | 1 | 1 | 1 | 1 | 8 |
| Landesberg, 2008 [55] | 1 | 1 | 1 | 1 | 0 | 1 | 1 | 1 | 7 |
| Li, 2022 [50] | 1 | 1 | 1 | 1 | 0 | 1 | 1 | 1 | 7 |
| Li, 2024 [51] | 1 | 1 | 1 | 0 | 0 | 1 | 1 | 1 | 6 |
| Liang, 2025 [52] | 1 | 0 | 1 | 1 | 0 | 0 | 0 | 1 | 4 |
| Lin, 2012 [53] | 1 | 1 | 1 | 0 | 0 | 1 | 1 | 1 | 6 |
| Lin, 2022 [54] | 1 | 0 | 1 | 1 | 0 | 0 | 0 | 0 | 3 |
| Medapati, 2015 [56] | 1 | 1 | 1 | 1 | 0 | 1 | 1 | 1 | 7 |
| Muoio, 2023 [57] | 1 | 0 | 0 | 0 | 0 | 0 | 0 | 1 | 2 |
| Nankali, 2016 [58] | 1 | 1 | 1 | 1 | 0 | 1 | 1 | 1 | 7 |
| Nasser, 2015 [59] | 1 | 0 | 1 | 1 | 0 | 0 | 0 | 0 | 3 |
| Qian, 2019 [66] | 1 | 1 | 1 | 0 | 0 | 1 | 1 | 1 | 6 |
| Qie, 2015 [67] | 1 | 1 | 1 | 1 | 0 | 1 | 1 | 1 | 7 |

| | | | | | | | | | |
|---|---|---|---|---|---|---|---|---|---|
| Rahimi, 2017 [60] | 1 | 1 | 1 | 1 | 0 | 1 | 1 | 1 | 7 |
| Sasahira, 2005 [61] | 1 | 1 | 1 | 0 | 0 | 1 | 1 | 1 | 6 |
| Sasahira, 2007 [62] | 1 | 1 | 1 | 0 | 0 | 1 | 1 | 1 | 6 |
| Sinduja, 2024 [63] | 1 | 1 | 1 | 0 | 0 | 1 | 1 | 1 | 6 |
| Tafani, 2010 [64] | 1 | 1 | 1 | 0 | 1 | 1 | 1 | 1 | 7 |
| Tateno, 2009 [65] | 1 | 1 | 1 | 0 | 0 | 1 | 1 | 1 | 6 |
| Wang, 2015 [68] | 1 | 1 | 1 | 1 | 0 | 1 | 1 | 1 | 7 |
| Wang, 2020 [69] | 1 | 0 | 0 | 0 | 0 | 0 | 0 | 1 | 2 |
| Wang, 2021 [70] | 1 | 1 | 1 | 0 | 0 | 1 | 1 | 1 | 6 |
| Wu, 2018 [71] | 1 | 0 | 1 | 0 | 0 | 0 | 0 | 1 | 3 |
| Xu, 2013 [72] | 1 | 1 | 1 | 0 | 0 | 1 | 1 | 1 | 6 |
| Xu, 2013 [73] | 1 | 1 | 1 | 1 | 0 | 1 | 1 | 1 | 7 |
| Yang, 2015 [74] | 1 | 1 | 1 | 1 | 1 | 1 | 1 | 1 | 8 |
| Yang, 2023 [75] | 1 | 0 | 0 | 0 | 0 | 0 | 0 | 1 | 2 |
| Yaser, 2012 [76] | 1 | 1 | 1 | 1 | 1 | 1 | 1 | 1 | 8 |
| Yin, 2018 [77] | 1 | 1 | 1 | 1 | 0 | 1 | 1 | 1 | 7 |
| Zhang, 2016 [78] | 1 | 1 | 1 | 1 | 0 | 1 | 1 | 1 | 7 |
| Zhao, 2014 [79] | 1 | 1 | 1 | 0 | 0 | 1 | 1 | 1 | 6 |
| Zhou, 2014 [80] | 1 | 1 | 1 | 0 | 0 | 1 | 1 | 1 | 6 |

1Indicates that cases are independently validated for cancer by a trained histopathologist; 2cases are from a representative population or drawn from the same community; 3control of appropriate negative control (benign, low-grade, non-regional lymph invading, deceased patient); 4control of adjacent normal tissue; 5specified as secondary disease states or infections; 6cancer samples were attained from medical records; 7collection method of cancer was included; 8approved identification technique was used to measure RAGE expression; 9total: minimum equals 1 star; maximum equals 8 stars.
