## Supplemental Table 3 for "The Role of the Receptor for Advanced Glycation End-Products in Cancer: Evidence from a Systematic Review and Meta-Analysis"

| Author, Year | Cell Line(s) | Organ of Origin | Culture Conditions* | Treatment/ Dose(s) | RAGE Expression | Growth Potential | Metastatic Potential | Apoptosis Evasion | NF-κB Expression |
|---|---|---|---|---|---|---|---|---|---|
| Abe, 2004 [184] | G361 | Skin | Standard | AGE2 & AGE3 (1mg/mL); anti-RAGE ab | Confirmed RAGE expression in G361 cell line | AGE2 and AGE3 increased cell proliferation (p<0.05; 0.01); inhibition of RAGE by anti-RAGE ab reduced proliferation in both groups (p<0.005) | AGE2 and AGE3 increased migration and invasion in both groups (p<0.005) | N/A | N/A |
| Alka, 2024 [207] | Panc01, Panc02 | Pancreas | Standard | Azeliragon (3μM); S100P & S100A2 & HMGB1 (0.1μM) | Confirmed RAGE expression in cell lines | S100P, S100A2, and HMGB1 increased cell proliferation (p<0.05; 0.01); inhibition of RAGE by azeliragon reduced proliferation (p<0.001; 0.0001); inhibition of RAGE by azeliragon negated ligand-induced proliferation (p<0.01; 0.05) | N/A | N/A | S100P, S100A2, and HMGB1 increased NF-kB (p<0.05; 0.01; 0.001); inhibition of RAGE by azeliragon reduced NF-kB (p<0.01; 0.001); inhibition of RAGE by azeliragon negated ligand-induced NF-kB (p>0.05; p<0.01; 0.001) |
| Amornsupak, 2022 [135] | MDA-MB-231 | Breast | Standard | rHMGB1 | RAGE silencing (shRNA) reduced RAGE expression | N/A | HMGB1 increased migration and invasion (p<0.05) | N/A | N/A |
| Arrè, 2024 [141] | Caco-2; SW620 | Colorectum | Standard | RAGE inhibitor | Confirmed RAGE expression in Caco-2 and SW620 | RAGE inhibition didn't affect cell proliferation (p>0.05) | N/A | N/A | N/A |
| Arumugam, 2005 [196] | BxPC-3 | Pancreas | Standard | anti-RAGE ab (1μg) | Confirmed RAGE expression in BxPC-3 cell line | RAGE inhibition reduced cell proliferation (p<0.05) | N/A | N/A | N/A |
| Arumugam, 2006 [81] | Panc01, BxPC-3 | Pancreas | Standard | S100P (100nM); cromolyn (100μM) | Confirmed RAGE expression and reduction with cromolyn | Cromolyn reduced S100P-induced cell proliferation (p<0.001) | Cromolyn reduced S100P-induced invasion (p<0.001) | Cromolyn didn't induce apoptosis evasion (p>0.05) | Cromolyn reduced S100P-induced NF-kB expression (p<0.001; 0.005) |
| Arumugam, 2012 [197] | MOH, HPAF II, Mpanc-96, BxPC-3 | Pancreas | Standard | S100P (2μM); siRAGE; RAP (10μM) | Confirmed RAGE expression and reduction with RAGE siRNA | RAP and RAGE siRNA reduced cell proliferation (p<0.05) | RAP and RAGE siRNA reduced migration (p<0.05) | N/A | RAP reduced S100P-induced NF-kB expression (p<0.05) |
| Bao, 2015 [82] | PC-3 | Prostate | Standard | AGEs (200 μg/mL); siRAGE | Confirmed RAGE expression and reduction with RAGE siRNA | RAGE silencing reduced cell proliferation (p<0.01) | N/A | N/A | N/A |
| Bartling, 2005 [83] | NCI-H358 | Lungs | Standard | flRAGE vector | Inserted flRAGE expression into cell line | flRAGE expression reduced cell proliferation (p<0.05) | N/A | N/A | N/A |
| Bassi, 2008 [84] | T98G | Brain | Standard | HMGB1 (50nM); anti-RAGE ab (10 μg/mL) | Confirmed RAGE expression in T98G cell line | Inhibition of RAGE by antibody negated HMGB1-induced proliferation (p<0.01) | Inhibition of RAGE by antibody negated HMGB1-induced migration (p<0.01) | N/A | N/A |
| Bhawal, 2005 [85] | LMF4, HSC3, patient-derived (KKp, KKm) | Oral | Standard | RAGE antisense (3μM) | Confirmed RAGE expression and reduction with RAGE antisense in cell lines | RAGE inhibition reduced cell proliferation in the metastatic lines (p=0.0217; 0.0023) and didn't in localized lines (p>0.05) | RAGE inhibition reduced migration and invasion in the metastatic lines (p<0.001) and didn't in localized lines (p>0.05) | N/A | N/A |
| Chang, 2023 [161] | JJ012, SW1353 | Musculoskeletal | Standard | CML (50; 100μM) | Confirmed RAGE expression and increased with CML | CML didn't increase RAGE viability (p>0.05) | CML increased migration and invasion (p<0.05) | N/A | CML increased NF-kB expression (p<0.05) |

| | | | | | | | | | |
|---|---|---|---|---|---|---|---|---|---|
| Chang, 2024 [162] | MG63, U2OS | Musculoskeletal | Standard | CML (50µM); anti-RAGE ab (10 µg/mL) | Confirmed RAGE expression and increased with CML | N/A | CML increased migration and invasion (p<0.05); Inhibition by anti-RAGE antibody negated CML-induced migration and invasion (p<0.05) | N/A | CML increased NF-kB expression (p<0.05) |
| Chen, 2014 [86] | HCT116 | Colorectum | Standard | AGEs-BSA (200; 400 mg/L); Glucose (25mM); anti-RAGE ab (10 µg/mL) | Confirmed RAGE expression and increased with AGEs-BSA and decreased with anti-RAGE ab | AGEs-BSA and AGEs-BSA+Glucose increased cell proliferation (p<0.05); inhibition by anti-RAGE antibody negated AGEs-BSA-induced cell proliferation (p<0.05) | N/A | AGEs-BSA+Glucose increased apoptosis evasion (p<0.05) | N/A |
| Chen, 2020 [170] | A549 | Lungs | Standard | RAGE-high cell line | Confirmed RAGE expression in RAGE-high lines | RAGE expression reduced cell proliferation (p<0.01) | RAGE expression reduced migration (p<0.01) | N/A | N/A |
| Chen, 2022 [87] | MiaPaCa-2 | Pancreas | Standard | 4-AAQB (5µM) | Confirmed RAGE expression and decreased with 4-AAQB | 4-AAQB decreased cell viability (p<0.001) | 4-AAQB decreased migration and invasion (p<0.01; 0.001) | 4-AAQB decreased apoptosis evasion (p<0.05) | N/A |
| Chen, 2022 [151] | MiaPaCa-2 | Pancreas | Standard | Lucidone (50µM) | Confirmed RAGE expression and decreased with lucidone | Lucidone decreased cell viability (p<0.01) | N/A | Lucidone decreased apoptosis evasion (p<0.01) | N/A |
| Chen, 2024 [186] | HCCLM3, RBE | Liver | Standard | siS100A6 | Treated cells with silencing RAGE ligand (siS100A6) | n/A | siS100A6 decreased migration (p<0.05) | N/A | N/A |
| Cho, 2024 [191] | A549 | Lungs | Standard | Cigarette smoke extract (CSE; 1.5%); siS100A8/9; siRAGE | Confirmed RAGE expression, increased with CSE, decreased with siS100A8/9 and siRAGE | Inhibition of AGEs and RAGE with siRNA negated CSE-induced cell viability (p<0.01; 0.001) | Inhibition of AGEs and RAGE with siRNA negated CSE-induced migration and invasion (p<0.05) | N/A | Inhibition of RAGE with siRNA negated CSE-induced NF-kB expression (p<0.05); inhibition of AGEs with siRNA didn't negate CSE-induced NF-kB expression (p>0.05) |
| Choi, 2011 [142] | SCC7 | Head and Neck | Standard | HMGB1 (2000 ng/mL); nifedipine (100µM) | Confirmed RAGE expression in cell lines | N/A | HMGB1 increased migration (p<0.0002) and inhibition of HMGB1 by nifedipine negated increased migration (p<0.0007) | N/A | N/A |
| Chung, 2015 [88] | N87 | Stomach | Standard | siRAGE; rhHMGB1 (10 µg/mL); HMGB1 transfection | Confirmed RAGE expression, increased with rHMGB1, decreased with siRAGE | N/A | Inhibition of RAGE with siRNA negated HMGB1-induced migration and invasion (p<0.05) | N/A | N/A |
| Dahlmann, 2014 [175] | HCT116, SW620, DLD-1 | Colorectum | Standard | rS100A4 (5 µg/mL); RAGE-OE; rsRAGE; anti-RAGE ab | Confirmed low RAGE expression in parental cells and increase with RAGE-OE | S100A4 and RAGE-OE doesn't increase cell proliferation (p>0.05) | S100A4 and RAGE-OE increased migration and invasion (p<0.05; 0.001; 0.005); inhibition of RAGE negated RAGE-OE and S100A4-induced migration and invasion (p<0.05; 0.005) | N/A | N/A |
| Dai, 2023 [32] | MDA-MB-231, BT549 | Breast | Standard | RP7 (50µM) | Confirmed RAGE expression in both cell lines | N/A | RP-7 inhibits migration and invasion (p<0.01) | RP7 inhibits apoptosis evasion (p<0.01) | N/A |

| | | | | | | | | | |
|---|---|---|---|---|---|---|---|---|---|
| Deng, 2017 [33] | SGC7901 | Stomach | Standard | AGEs (100 µg/mL); anti-RAGE | Confirmed RAGE expression, increased with AGEs, decreased with anti-RAGE | N/A | AGEs increased migration and invasion (p<0.05); inhibition of RAGE negated AGEs-induced migration and invasion (p<0.05) | N/A | N/A |
| Deng, 2017 [34] | LoVo, SW620 | Colorectum | Standard | AGEs (200 µg/mL); anti-RAGE (5 µg/mL) | Confirmed RAGE expression, increased with AGEs, decreased with anti-RAGE | N/A | AGEs increased migration and invasion (p<0.05); inhibition of RAGE negated AGEs-induced migration and invasion (p<0.05) | N/A | N/A |
| Dhumale, 2015 [168] | MCF-7 | Breast | Standard | Quercetin (50µM) | Confirmed RAGE expression, decreased with quercetin | Quercetin reduced cell viability and proliferation (p<0.05) | N/A | N/A | Quercetin reduced NF-kB expression (p<0.05) |
| Elangovan, 2012 [216] | LNCaP, DU145 | Prostate | Standard | rHMGB1 (1 µg/mL); shRAGE | Confirmed RAGE expression, decreased with shRNA | Inhibition of RAGE by shRNA negated HMGB1-induced cell proliferation (p<0.01) | N/A | N/A | N/A |
| El-Far, 2018 [140] | C6 | Brain | Standard | HMGB1 (1 µg/mL); AGE-BSA (1 µg/mL); papaverine (20 µM) | Treated cells with RAGE ligands (HMGB1, AGEs-BSA) | N/A | N/A | N/A | HMGB1 and AGEs-BSA increased NF-kB expression (p<0.05; 0.01); RAGE inhibition by papaverine negated RAGE-ligand-induced NF-kB expression (p<0.01) |
| El-Far, 2018 [140] | HT1080 | Musculoskeletal | Standard | flRAGE vector, papaverine (20 µM) | Confirmed no RAGE expression in parental cells and increased with flRAGE vector | RAGE inhibition by papaverine decreased cell proliferation (p<0.01) | RAGE inhibition by papaverine decreased migration and invasion (p<0.01) | N/A | N/A |
| Fuentes, 2007 [169] | SW480 | Colorectum | Standard | S100P (100; 1000 nM); AmphP (100 µM) | Confirmed RAGE expression in SW480 cell line | S100P increased cell proliferation (p<0.05; 0.01); Blocking of RAGE-AGE interation by AmphP negates S100P-induced cell proliferation (p<0.01) | S100P increased migration (p<0.05) | N/A | N/A |
| Gao, 2024 [89] | A549, H1299 | Lungs | Standard | Isoflurane, FPS-ZM1 (0.5 nM) | Confirmed RAGE expression, increased with isoflurane, decreased with FPS-ZM1 | Inhibition of RAGE negates isoflurane-induced cell viability decrease (p<0.001) | N/A | Inhibition of RAGE doesn't negates isoflurane-induced apoptosis (p>0.05) | N/A |
| Geicu, 2020 [90] | C2BBe1 | Colorectum | Standard | AGEs-Csn (200 µg/mL); anti-RAGE IgG | Confirmed RAGE expression, increased with AGEs-Csn, decreased with anti-RAGE IgG | AGEs-Csn increased cell proliferation and viability (p<0.05; 0.001); inhibition of RAGE didn't impact AGEs-Csn-induced cell proliferation and viability (p>0.05) | N/A | N/A | AGEs-Csn didn't affect NF-kB expression (p>0.05); inhibition of RAGE reduced NF-kB expression (p<0.05) |
| Ghavami, 2008 [190] | MDA-MB-231 | Breast | Standard | S100A8/A9 (10 µg/mL); siRAGE; RAGE blocking ab | Confirmed RAGE expression, increased with S100A8/A9, decreased with siRAGE and RAGE blocking ab | S100A8/A9 increased cell proliferation (p<0.05); inhibition of RAGE with siRAGE and RAGE blocking ab negated S100A8/A9-induced cell proliferation (p<0.05) | N/A | N/A | N/A |

| | | | | | | | | | |
|---|---|---|---|---|---|---|---|---|---|
| Ghavami, 2008 [190] | SHEP | Brain | Standard | S100A8/A9 (10 µg/mL); RAGE blocking ab | Confirmed RAGE expression, increased S100A8/A9, decreased with RAGE blocking ab | S100A8/A9 increased cell proliferation (p<0.05); inhibition of RAGE with RAGE blocking ab negated S100A8/A9-induced cell proliferation (p<0.05) | N/A | N/A | N/A |
| Gurumayum, 2024 [159] | HepG2 | Liver | Standard | Methylglyoxal (2.5 mM) | Confirmed RAGE expresion, increased with Methylglyoxal | Methylglyoxal decreased cell viability (p<0.001) | N/A | N/A | N/A |
| Guzmán, 2019 [206] | Panc-1, Mia-PaCa-2 | Pancreas | Standard | Curcumin (30µM); scalarin (10 µg/mL) | Confirmed RAGE expression, decreased with curcumin and scalarin | N/A | N/A | Scalarin and curcumin didn't affect apoptosis evasion (p>0.05) | Scalarin and curcumin didn't affect NF-kB expression (p>0.05) |
| Han, 2023 [204] | HeLa | Cervix | Standard | RAGE KO and KO 2 | Confirmed RAGE expression, decreased after genetic KO | RAGE KO decreased cell viability (p<0.0001) | N/A | N/A | N/A |
| He, 2018 [137] | HeLa | Cervix | Standard | Ethyl pyruvate (10 mM), glycyrrhizin (1 mM), HMGB1 KO | Confirmed RAGE expression, decreased with HMGB1 KO | Glycyrrhizin, ethyl pyruvate, and HMGB1 KO decreased cell proliferation (p>0.05; <0.01) | N/A | N/A | N/A |
| He, 2018 [137] | HT29, HCT116 | Colorectum | Standard | Ethyl pyruvate (10 mM), glycyrrhizin (1 mM), HMGB1 KO | Confirmed RAGE expression, decreased with HMGB1 KO | Glycyrrhizin, ethyl pyruvate, and HMGB1 KO decreased cell proliferation (p>0.05; <0.01) | N/A | N/A | N/A |
| Herwig, 2016 [91] | A375 | Lungs | Standard | hRAGE transfection, sRAGE, S100A4 | Confirmed RAGE expression in A375 cell line | N/A | hRAGE transfection and S100A4/hRAGE transfection increased migration (p<0.05); sRAGE didn't affect migration (p>0.05) | N/A | N/A |
| Herwig, 2016 [143] | A375 | Lungs | Standard | hS100A4, hRAGE transfection, anti-RAGE ab, sRAGE, S100A4 (3000 ng/mL) | Confirmed RAGE expression, increased with hRAGE and hS100A4 | hS100A4 increased cell proliferation (p<0.05) | hS100A4 increased migration (p<0.05); RAGE transfection and inhibition decreased migration (p<0.05; >0.05); inhibition of RAGE negated S100A4-induced migration and invasion (p<0.05; >0.05) | N/A | hS100A4 increased NF-kB expression (p<0.05) |
| Hou, 2018 [185] | A549, LLC | Lungs | Standard | S100A4 (1 µg/mL), FPS-ZM1 (75nM) | Confirmed RAGE expression, decreased with FPS-ZM1, increased with S100A4 | FPS-ZM1 decreased cell viability (p<0.05); S100A4 negated FPS-ZM1-induced cell viability (p<0.05) | N/A | N/A | N/A |
| Hsu, 2020 [166] | MiaPaCa-2 | Pancreas | Standard | Pterostilbene (50µM) | Confirmed RAGE expression, decreased pterostilbene | Pterostileben decreased cell viability (p<0.001) | N/A | Pterostilbene decreased apoptosis evasion (p<0.01) | N/A |

| | | | | | | | | | |
|---|---|---|---|---|---|---|---|---|---|
| Huang, 2018 [136] | SW480 | Colorectum | Standard | Oxaplatin (10µM); capecitabine (10µM); irinotecan (5µM); pcDNA3.1-wtRAGE; shRAGE | Confirmed RAGE expression, increased with wtRAGE, decreased with shRAGE | RAGE transfection negates chemtherapeutic-induced cell viability reduction ($p<0.05$) | N/A | RAGE silencing increases chemtherapeutic-induced apoptosis ($p<0.05$) | N/A |
| Huang, 2018 [218] | LoVo | Colorectum | Standard | anti-RAGE ab (5 µg/mL) | Treated cells with anti-RAGE antibody | N/A | Inhibition of RAGE decreased invasion ($p<0.01$) | N/A | Inhibition of RAGE decreased NF-kB expression ($p<0.05$; 0.01) |
| Huttunen, 2002 [183] | HT1080 | Musculoskeletal | Standard | Amphoterin inhibitor (500µM) | Confirmed RAGE binding to amphoterin inhibitor | N/A | Amphoterin inhibitor decreased migration (0.00001) | N/A | N/A |
| Ichikawa, 2011 [189] | MC38, Caco-2 | Colorectum | Standard | S100A8/A9 (10 µg/mL); anti-RAGE ab | Confirmed RAGE expression, increased with S100A8/A9, decreased with anti-RAGE ab | N/A | N/A | N/A | S100A8/A9 increased NF-kB expression ($p<0.005$); inhibition of RAGE by anti-RAGE ab negated S100A8/A9-induced NF-kB expression ($p<0.05$; 0.01) |
| Inada, 2019 [92] | T98G | Brain | Standard | HMGB1 (1 µg/mL); papaverine (300µM) | Confirmed RAGE expression in T98G cell line | Papaverine negated HMGB1-induced cell proliferation ($p<0.05$) | Papaverine negated HMGB1-induced migration ($p<0.01$) | N/A | N/A |
| Inada, 2019 [93] | U87MG, T98G | Brain | Standard | Papaverine (30µM) | Used RAGE-HMGB1 interaction inhibitor | Papaverine decreased cell proliferation ($p<0.05$; 0.001) | N/A | N/A | N/A |
| Ishibashi, 2013 [154] | MCF-7 | Breast | Standard | AGEs-BSA (100 µg/mL); anti-RAGE ab (5 µg/mL) | Confirmed RAGE expression, increased with AGEs-BSA | AGEs increased cell proliferation ($p<0.01$); inhibition by anti-RAGE antibody negated AGEs-induced cell proliferation ($p<0.01$) | N/A | N/A | N/A |
| Ji, 2022 [222] | Eca-109, TE-1 | Esophageal | Standard | oeHMGB1, verbascoside (50µM; 100µM) | Confirmed RAGE expression, decreased with verbascoside, increased with oeHMGB1 | Verbascoside decreased cell viability ($p<0.05$); oeHMGB1 negated verbascoside-induced cell viability decrease ($p<0.05$) | Verbascoside decreased migration and invasion ($p<0.05$); oeHMGB1 negated verbascoside-induced migration and invasion decrease ($p<0.05$) | Verbascoside decreased apoptosis evasion ($p<0.05$); oeHMGB1 negated verbascoside-induced apoptosis evasion decrease ($p<0.05$) | N/A |
| Jia, 2020 [157] | HeLa | Cervix | Standard | mir-218 | Confirmed RAGE expression, decreased with mir-218 | miR-218 decreased cell viability ($p<0.05$) | miR-218 decreased migration and invasion ($p<0.05$) | miR-218 decreased apoptosis evasion ($p<0.05$) | N/A |
| Jiang, 2025 [160] | 4T1 | Breast | Standard | FPS-ZM1; 2f-LIGRL-NH2 | Treated cells with RAGE inhibitor | N/A | FPS-ZM1 decreased migration ($p<0.0001$) | N/A | N/A |
| Jin, 2011 [193] | KYSE180 | Esophageal | Standard | S100A14 (10 µg/mL; 20 µg/mL); dnRAGE; AmphP (5µM); siRAGE (50nM); RAGE transfection | Confirmed RAGE expression, decreased with RAGE siRNA, dnRAGE, and AmphP, increased with RAGE transfection and S100A14 | S100A14 increased cell viability ($p<0.05$); inhibition of RAGE by dnRAGE, AmphP, and siRAGE negated S100A14-induced cell viability ($p<0.05$) | N/A | N/A | S100A14 didn't affect NF-kB expression ($p>0.05$); RAGE transfection and S100A14 increased NF-kB expression ($p<0.05$) |

| | | | | | | | | | |
|---|---|---|---|---|---|---|---|---|---|
| Jin, 2025 [94] | PEO-14 | Ovaries | Standard | siRAGE; ASA (3mM); rhAPE1/Ref-1 (5 µg/mL) | Confirmed RAGE expression, increased with ASA, decreased with siRAGE | siRAGE and siRAGE/ASA didn't affect cell viability (p>0.05); silencing RAGE negated ASA+ rhAPE1/Ref-1-induced viability decrease (p<0.01) | N/A | ASA + rhAPE1/Ref-1 reduced apoptosis evasion (p<0.001) | N/A |
| Jing, 2015 [164] | TE-11, Eca-109 | Esophageal | Standard | miR-185 | Confirmed RAGE expression, decreased with miR-185 | miR-185 didn't affect RAGE expression (p>0.05) | miR-185 decreased migration and invasion (p<0.001) | N/A | N/A |
| Jube, 2012 [95] | REN, Phi | Mesotheliom a | Standard | anti-RAGE ab, HMGB1 (100 ng/mL) | Confirmed RAGE expression in both cell lines | RAGE inhibition reduces cell viability and induces cytotoxicity (p<0.05) | RAGE inhibition decreases migration (p<0.05); RAGE inhibition decreases and HMGB1 doesn't increase invasion (p<0.05) | N/A | N/A |
| Kang, 2010 [208] | Panc02, Panc2.03 | Pancreas | Standard | shRAGE, pUNO1-RAGE | Confirmed RAGE expression in both cell lines, decreased with shRAGE, increased with plasmid-RAGE transfection | RAGE silencing increases chemotherapeutic-induced cytotoxicity (p<0.05); RAGE overexpression decreases chemotherapeutic-induced cytotoxicity (p<0.05) | N/A | Silencing RAGE increases chemotherapeutic-induced apoptosis (p<0.05) | N/A |
| Kang, 2011 [209] | Panc02, Panc2.03 | Pancreas | Standard | shRAGE, GEM (100nM) | Confirmed RAGE expression in both cell lines, decreased with shRAGE | N/A | N/A | RAGE silencing decreased apoptosis evasion (p<0.05) | RAGE silencing decreased NF-kB expression (p<0.05); RAGE silencing synergistically decreased NF-kB expression with GEM (p<0.05) |
| Kang, 2012 [202] | Panc02 | Pancreas | Standard | shRAGE, pUNO1-RAGE | Confirmed RAGE expression, decreased with shRAGE, increased with pUNO1-RAGE | RAGE silencing decreased cell proliferation (p<0.05); re-expression of RAGE after RAGE silencing increased cell proliferation (p<0.05) | N/A | N/A | N/A |
| Kang, 2014 [174] | Panc02 | Pancreas | Normoxia, Hypoxia | shRAGE | Confirmed RAGE expression, increased in hypoxic conditions | RAGE silencing didn't affect cell viability in normoxia (p>0.05) and increased cell viability in hypoxia (p<0.05) | N/A | N/A | N/A |
| Kang, 2014 [203] | Panc02, Panc2.03 | Pancreas | Standard | shRAGE, HMGB1 (10 µg/mL) | Confirmed RAGE expression, decreased with shRAGE, increased with HMGB1 | RAGE silencing decreased cell proliferation even when stimulated with HMGB1 (p<0.05) | N/A | N/A | N/A |
| Kataoka, 2012 [187] | Saos-2, Hu09 | Musculoskele tal | Standard | S100A7 (1 µg/mL; 10 µg/mL); siRAGE | Confirmed RAGE expression in both cell lines, decreased with siRAGE | S100A7 didn't impact cell proliferation (p>0.05) | S100A7 didn't impact migration (p>0.05); S100A7 increased invasion (p<0.05); RAGE silencing decreased migration and invasion (p<0.05) | N/A | N/A |
| Khoo, 2023 [45] | LNCaP, 22rV1, DU145, PC3 | Prostate | Standard | GA-AGEs, anti-RAGE ab | Confirmed RAGE expression in all cell lines | RAGE inhibition decreased cell proliferation (p<0.05) | N/A | N/A | N/A |

| | | | | | | | | | |
|---|---|---|---|---|---|---|---|---|---|
| Kim, 2025 [96] | A549, NCI-H23, NCI-H60, NCI-H596 | Lungs | Standard | HMGB1 (0.4µM); HMGB1-peptide (300µm); rotenone (2µM) | Confirmed RAGE expression in all cell lines, increased with HMGB1, decreased with HMGB1-peptide | HMGB1-peptide decreased cell viability (p<0.0001); HMGB1 negated rotenone-induced cell viability (p<0.05); HMGB1-peptide negated HMGB1-induced cell viability (p<0.0001) | HMGB1-peptide decreased migration (p<0.001); HMGB1 increased migration (p<0.05); HMGB1 negated rotenone-induced migration (p<0.05) | N/A | HMGB1-peptide decreased NF-kB expression (p<0.05) |
| Kim, 2025 [96] | HCT-116 | Colorectum | Standard | HMGB1 (0.4µM); HMGB1-peptide (300µm); rotenone (2µM) | Confirmed RAGE expression, increased with HMGB1, decreased with HMGB1-peptide | HMGB1-peptide decreased cell viability (p<0.0001); HMGB1 negated rotenone-induced cell viability (p<0.05); HMGB1-peptide negated HMGB1-induced cell viability (p<0.0001) | HMGB1-peptide decreased migration (p<0.0001); HMGB1 increased migration (p<0.01 | N/A | N/A |
| Kim, 2025 [96] | AsPC-1 | Pancreas | Standard | HMGB1 (0.4µM); HMGB1-peptide (300µm); rotenone (2µM) | Confirmed RAGE expression, increased with HMGB1, decreased with HMGB1-peptide | HMGB1-peptide decreased cell viability (p<0.0001); HMGB1 negated rotenone-induced cell viability (p<0.05); HMGB1-peptide negated HMGB1-induced cell viability (p<0.0001) | HMGB1-peptide decreased migration (p<0.0001); HMGB1 increased migration (p<0.05) | N/A | N/A |
| Kinoshita, 2019 [97] | B16-BL6 | Lungs | Standard | S100A8/A9 (100 ng/mL); exRAGE-Fc (1 µg/mL) | Confirmed RAGE expression in B16-BL6 cell lines | N/A | exRAGE-Fc negated S100A8/A9-induced invasion (p<0.01) | N/A | N/A |
| Ko, 2014 [98] | SAS | Oral | Standard | RAGE RNAi (20nM) | Confirmed RAGE expression, decreased with RAGE RNAi | N/A | RAGE silencing decreased migration (p=0.002) | N/A | N/A |
| Kobayashi, 2007 [47] | A549, LK-1 | Lungs | Standard | flRAGE, esRAGE | Treated cell lines with flRAGE and esRAGE transfection | flRAGE didn't increase cell proliferation in A549 (p>0.05) and decreased cell proliferation in LK-1 (p<0.05); esRAGE decreased cell proliferation (p<0.05) | N/A | N/A | N/A |
| Kuniyasu, 2002 [99] | MKN28 | Stomach | Standard | Anti-sense RAGE oligodeoxynucleotide (3µM) | Confirmed RAGE expression increase with sense-ODN and decrease with anti-sense-ODN | Anti-sense RAGE ODN didn't impact cell proliferation (P>0.05) | Anti-sense RAGE ODN decreased invasion (p<0.0001) | N/A | N/A |
| Kuniyasu, 2003 [100] | Colo320, WiDR | Colorectum | Standard | Anti-sense RAGE S-ODN (3µM) | Confirmed RAGE expression, increase with sense-ODN, and decrease with anti-sense-ODN | N/A | Anti-sense RAGE S-ODN decreased migration and invasion (p<0.0001; 0.0036) | N/A | N/A |
| Kwak, 2017 [215] | MDA-MB-231, MDA-MB-231-4175, 4T1, 67NR, DT28 | Breast | Standard | oeRAGE, RAGE sh(66,10,12) FPS-ZM1 (1µM; 25µM), siRAGE | Confirmed RAGE expression, increased with oeRAGE, decreased with RAGE shs, used RAGE inhibitor (FPS-ZM1) | RAGE overexpression sometimes increased cell proliferation (p<0.05; >0.05); RAGE silencing and inhibition sometimes decreased cell proliferation (p<0.05; >0.05) | RAGE silencing and inhibition mostly decreased invasion (p<0.05; >0.05) | N/A | N/A |
| Lai, 2021 [101] | Jurkat, HL-60 | Blood | Standard | rHMGB1 (50 ng/mL); shRAGE | Confirmed RAGE expression, decreasd with shRAGE | RAGE silencing negated HMGB1-induced cell viability (p<0.01) | N/A | RAGE silencing negated HMGB1-induced apoptosis evasion (p<0.01) | N/A |

| | | | | | | | | | |
|---|---|---|---|---|---|---|---|---|---|
| Lan, 2019 [167] | MiaPaCa-2 | Pancreas | Standard | siRAGE, quercetin (50μM) | Confirmed RAGE expression, decreased with siRAGE and quercetin | RAGE silencing didn't affect cell viability (p>0.05); RAGE silencing increased chemotherapeutic effects on cell viability (p<0.05); quercetin decreased cell viability (p<0.05) | N/A | RAGE silencing and quercetin with chemotherapeutics increased apoptosis (p<0.05) | N/A |
| Lata, 2014 [147] | MCF-7 | Breast | Standard | siRAGE, 17α-ethinyl-estradiol (10nM) | Confirmed RAGE expression, decreased with siRAGE, increased with 17-EE | RAGE silencing didn't affect cell viability (p>0.05); RAGE silencing negated 17-EE-induced cell viability (p<0.05) | N/A | N/A | N/A |
| Lee, 2015 [102] | MDA-MB-231 | Breast | Standard | RAGE KD, ASA (5mM), ac-APE1/Ref-1, APE1/Ref-1 | Confirmed RAGE expression, increased with ASA | RAGE knockdown removed ASA, Ac-APE1/Ref-1, APE1/Ref-1 reductions on cell viability (p<0.05; >0.05) | N/A | N/A | N/A |
| Lee, 2018 [217] | MDA-MB-231 | Breast | Standard | oeRAGE, RAGE KD, rhAc-APE1 (1μg) | Confirmed RAGE expression, increased with oeRAGE, showed ac-APE1/Ref-1 interacted with RAGE | Ac-APE1 with oeRAGe decreased cell viability (p<0.01); Ac-APE1 with RAGE KD decreased cell viability (p<0.05) | N/A | N/A | N/A |
| Li, 2012 [103] | GBC-SD, SGC-996 | Gall bladder | Standard | Ethyl pyruvate (80mM) | Confirmed RAGE expression, decreased with EP | Ethyl pyruvate decreased cell proliferation (p<0.05) | Ethyl pyruvate decreased invasion (p<0.01) | Ethyl pyruvate decreased apoptosis evasion (p<0.01) | N/A |
| Li, 2018 [172] | Bel7402, HCCLM3 | Liver | Standard | shRAGE, sorafenib (10μM) | Confirmed RAGE expression, decreased with shRAGE and sorafenib | RAGE silencing increased sorafenib-induced cell viability decrease (p<0.001) | N/A | N/A | N/A |
| Li, 2020 [104] | SiHa, CaSki | Cervix | Standard | FPS-ZM1 (1μM; 8μM); GFP-RAGE transfection; RAGE-KD | Confirmed RAGE expression, decreased with RAGE KD, increased with GFP-RAGE transfection | RAGE inhibition decreased cell proliferation (p<0.05; 0.01; 0.001); RAGE overexpression increased cell proliferation (p<0.01) | N/A | RAGE inhibition decreased apoptosis evasion (p<0.05; 0.01); RAGE overexpression increased apoptosis evasion (p<0.05) | N/A |
| Li, 2020 [211] | SW480, T84 | Colorectum | Standard | oeRAGE, shRAGE, scutellarein (40μM; 80μM), oeCDC4/RAGE | Confirmed RAGE expression, decreased with scutellarein and shRAGE, increased with oeRAGE | RAGE overexpression negated scutellarein and oeCDC4-induced cell proliferation decrease (p<0.05); RAGE silencing increases the scutellarein and oeCDC4-induced cell proliferation decrease (p<0.05) | N/A | RAGE overexpression reduces scutellarein and oeCDC4-induced apoptosis (p<0.05); RAGE silencing increases scutellarein-induced apoptosis (p<0.05) | N/A |
| Li, 2024 [51] | SCC25 | Head and Neck | Normal media; MLKL-FD conditioned media | ISG15, FPS-ZM1 (10μM) | Confirmed RAGE expression, decreased with FPS-ZM1 | N/A | RAGE activation increased migration and invasion (p<0.001); inhibition of RAGE with FPS-ZM1 reversed ISG15- and MLKL-FD media-induced migration and invasion (p<0.001) | N/A | N/A |
| Li, 2024 [224] | SK-BR-3, MDA-MB-231 | Breast | Standard | NGR1 (400μM) | Confirmed RAGE expression, decreased with NGR1 | NGR1 decreased cell viability (p<0.0001) | N/A | N/A | N/A |

| | | | | | | | | | |
|---|---|---|---|---|---|---|---|---|---|
| Liang, 2011 [146] | SW480 | Colorectum | Standard | shRNA-4 | Confirmed RAGE expression, decreased with shRNA-4 | RAGE silencing didn't affect cell viability (p>0.05) | RAGE silencing decreased migration (p<0.01) | N/A | N/A |
| Liao, 2018 [105] | A549 | Lungs | Standard | High glucose (25mM); anti-RAGE ab | Confirmed RAGE expression, increased with high glucose, decreased with anti-RAGE ab | RAGE inhibition with high glucose media increased cell proliferation (p<0.05) | RAGE inhibition with high glucose media increased migration (p<0.05) | N/A | N/A |
| Liao, 2023 [117] | A549 | Lungs | Standard | RAGE transfection, gefitinib (50μM) | Confirmed RAGE expression with RAGE transfection | RAGE transfection decreased gefitinib-induced loss of cell viability (p<0.01) | N/A | RAGE transfection increased apoptosis evasion (p<0.001) | N/A |
| Lin, 2012 [53] | OS-RC-2 | Kidneys | Standard | HMGB1 (50nM); siRAGE | Confirmed RAGE expression, decreased with siRAGE | RAGE silencing didn't affect cell viability (p>0.05) | RAGE silencing didn't affect migration and invasion (p>0.05); RAGE silencing negated HMGB1-induced migration and invasion (p<0.05) | N/A | N/A |
| Lin, 2020 [221] | MiaPaCa-2; MiaPaCa-2(GEM-resistant) | Pancreas | Standard | siRAGE (25nM), gemcitabine (0.5μM) | Confirmed RAGE expression, decreased with siRAGE, increased with GEM-resistant line | RAGE silencing didn't impact cell viability or proliferation in parental or GEM-resistant lines (p>0.05); RAGE silencing increased tumor cell susceptibility to GEM in both lines (p=0.0085; 0.0252; 0.0015) | N/A | N/A | N/A |
| Liu, 2025 [106] | SCC-9 | Oral | Standard | Berberine (40μM) | Confirmed RAGE expression, decreased with berberine | N/A | RAGE inhibition by berberine decreased migration (p<0.001) | RAGE inhibition decreased apoptosis evasion (p<0.001) | N/A |
| Madhavan, 2021 [107] | HeLa | Cervix | Standard | vRAGE, RAGEWT transfection | Confirmed RAGE expressions after transfection | RAGE expression increased cell proliferation (p<0.001; 0.0001) | RAGE expression increased migration and invasion (p<0.01; 0.001) | N/A | N/A |
| Magna, 2023 [173] | MDA-MB-231-4175; 4T1 | Breast | Standard | FPS-ZM1 (1μM); TTP488 (1μM) | Treated cell lines with RAGE inhibitors | RAGE inhibition mostly didn't affect cell viability (p>0.05; <0.05) | RAGE inhibition decreased migration and invasion (p<0.01; 0.001; 0.0001) | N/A | N/A |
| Matou-Nasri, 2017 [108] | MCF-7 | Breast | Standard | MG-BSA-AGEs (200 μg/mL) | Confirmed RAGE expression, increased with MG-BSA-AGEs | MG-BSA-AGEs sometimes increased cell viability and proliferation (p>0.05; <0.05) | MG-BSA-AGEs didn't affect migration (p>0.05) | N/A | N/A |
| Medapati, 2015 [56] | FTC236 | Thryoid | Standard | rhS100A4 (500nM); AGE-BSA (10 μg/mL); siRAGE | Confirmed RAGE expression, decreased with siRAGE | rhS100A4 didn't affect cell viability or proliferation (p>0.05; =0.067) | rhS100A4 and AGE-BSA increased migration (p<0.05; 0.005); RAGE silencing didn't impact rhS100A4- and AGE-BSA-induced migration (p>0.05) | N/A | N/A |

| | | | | | | | | | |
|---|---|---|---|---|---|---|---|---|---|
| Meghnani, 2014 [176] | WM115 | Skin | Standard | RAGE transfection, oeRAGE, siRAGE | Confirmed RAGE expression, increased with oeRAGE, decreased with siRAGE | RAGE transfection decreased cell proliferation (p<0.05); RAGE silencing increased cell proliferation (p<0.05) | RAGE transfection increased migration and invasion (p<0.01); RAGE overexpression increased migration (p<0.05) and didn't affect invasion (p>0.05); RAGE silencing decreased migration (p<0.05) | N/A | RAGE transfection decreased NF-kB expression (p<0.01) |
| Méndez, 2018 [109] | MDA231i, BT549, Hs578T | Breast | Standard | shHMGB1, rHMGA1, sRAGE, anti-HMGA1 ab | Confirmed RAGE expression, increased with rHMGA1, confirmed binding to HMGA1 | Silencing and stimulating HMGA1 didn't affect proliferation (p>0.05) | anti-HMGA1 antibody didn't affect invasion (p>0.05); sRAGE reduced migration in the presence of HMGA1 (p=0.0413) | N/A | N/A |
| Menini, 2018 [198] | MiaPaCa-2 | Pancreas | Standard | CML-HSA (50 µg/mL); RAP (10µM) | Confirmed RAGE expression, increased with CML-HSA | RAP negated AGE-HSA-induced cell proliferation (p<0.001) | N/A | N/A | RAP negated AGE-HSA-induced NF-kB expression (p<0.001) |
| Mitsui, 2019 [220] | PK-8 | Pancreas | Standard | exRAGE-Fc decoy (1 µg/mL) | Treated with a decoy RAGE receptor | N/A | exRAGE-Fc decreased migration (p<0.001) | N/A | N/A |
| Nakamura, 2017 [179] | G361 | Skin | Standard | BSA-AGEs (1 mg/mL); RAGE aptamer (100nM) | Treated with RAGE ligand and inhibitor | BSA-AGEs increased cell proliferation (p<0.01); RAGE-aptamer negated BSA-AGEs-induced cell proliferation (p<0.01) | N/A | N/A | N/A |
| Nam, 2021 [156] | 786-O, A498 | Kidneys | Standard | AGE4 (100 µg/mL; 800 µg/mL) | Confirmed RAGE expression, increased with AGE4 | AGE4 increased cell proliferation (p<0.001; 0.0001) | AGE4 increased migration (p<0.05) | AGE4 increased apoptosis evasion (p<0.05; 0.01) | N/A |
| Ojima, 2014 [110] | G361 | Skin | Standard | AGE-BSA (100 µg/mL); AGE-aptamer (2µM) | Confirmed RAGE expression, decreased with AGE-aptamer | AGE-BSA increased cell proliferation (p<0.01); AGE-aptamer negated AGE-BSA-induced cell proliferation (p<0.01) | N/A | N/A | N/A |
| Pan, 2018 [149] | PC9, L78 | Lungs | Standard | lncRAGE-OE | Confirmed RAGE expression, down-regulated with lncRAGE-OE | lncRAGE-OE decreased cell proliferation (p<0.01) | lncRAGE-OE decreased migration (p=0.026); lncRAGE-OE didn't affect invasion (p=0.732) | lncRAGE-OE decreased apoptosis evasion (p<0.05; 0.001) | N/A |
| Pan, 2022 [111] | MDA-MB-231 | Breast | Standard | AGEs-BSA (20µg/mL; 100µg/mL) | Confirmed RAGE expression, increased with AGEs-BSA | AGEs-BSA decreased cell viability (p<0.05) | AGEs-BSA increased migration and invasion (p<0.05) | N/A | AGEs-BSA decreased NF-kB expression (p<0.05) |
| Popa, 2014 [112] | MelJuSo, SK-Mel28 | Skin | Standard | RAGE siRNA #1, #2 | Confirmed RAGE expression in both cell lines | N/A | RAGE silencing reduces migration in MelJuSo (p<0.05; 0.01) and didn't affect SK-Mel28 (p>0.05) | N/A | N/A |
| Pujals, 2023 [180] | MDA-MB-231, BT549, Hs578T | Breast | Standard | shRAGE, FPS-ZM1 (30µM), TTP488 (2µM), HMGA1-Mab-21,42,47 | Confirmed RAGE expression, decreased with shRAGE | N/A | RAGE silencing decreased invasion and migration (p<0.05; 0.001); RAGE inhibitors and anti-RAGE ligand antibodies decreased invasion (p<0.05; 0.01; 0.001) | N/A | N/A |

| | | | | | | | | | |
|---|---|---|---|---|---|---|---|---|---|
| Qian, 2019 [66] | HCT116, SW480 | Colorectum | Standard | HMGB1 (100 ng/mL), siRAGE | Confirmed RAGE expression, decreased with siRAGE | HMGB1 increased cell proliferation (p<0.05); RAGE silencing negated HMGB1-induced cell proliferation (p<0.05) | N/A | N/A | N/A |
| Qiao, 2016 [113] | Bel-7402, Bel-7404 | Liver | Standard | AGER-sh12 | Confirmed RAGE expression, decreased with AGER-sh12 | RAGE silencing decreased cell proliferation and viability (p<0.01) | N/A | N/A | N/A |
| Ray, 2020 [152] | A549 | Lungs | Standard | LPA (5µM); shRAGE | Confirmed RAGE expression, decreases with shRAGE | RAGE silencing didn't affect cell proliferation (p>0.05); RAGE silencing negated LPA-induced cell proliferation (p<0.001) | RAGE silencing negated LPA-induced invasion (p<0.001) | N/A | N/A |
| Ray, 2020 [152] | MDA-MB-231, MCF-7 | Breast | Standard | LPA (5µM); shRAGE | Confirmed RAGE expression, decreases with shRAGE | RAGE silencing didn't affect cell proliferation (p>0.05); RAGE silencing negated LPA-induced cell proliferation (p<0.05; 0.01) | RAGE silencing didn't affect invasion (p>0.05); RAGE silencing negated LPA-induced invasion (p<0.001) | N/A | N/A |
| Reeb, 2015 [188] | THJ-11T, THJ-16T | Thryoid | Standard | S100A8 (20 µg/mL), anti-S100A8 ab (1 µg/mL), shS100A8, FPS-ZM1 (100µM) | Confirmed binding between RAGE and S100A8, treated cells with RAGE inhibitor | S100A8 increased cell proliferation in THJ-11T (p<0.05) and didn't in THJ-16T (p>0.05); anti-S100A8 antibody decreased cell proliferation in both lines (p<0.05; 0.01); FPS-ZM1 decreased cell proliferation in both lines (p<0.01; 0.0001) | N/A | Anti-S100A8 antibody decreased apoptosis evasion (p<0.05) | N/A |
| Rehbein, 2008 [158] | H358 | Lungs | Standard | S100P, oeRAGE | Treated with RAGE ligand and transfected with overexpressing RAGE plasmid | Overexpressing RAGE increased S100P-induced cell proliferation (p<0.01) | Overexpressing RAGE negated S100P-induced migration (p<0.05) | N/A | N/A |
| Ren, 2021 [200] | HSC-4 | Oral | Standard | RAGE-KD | Confirmed RAGE expression, decreased with RAGE-KD | RAGE-KD decreased proliferation (p<0.0001) | RAGE-KD decreased invasion (p<0.0001) | N/A | N/A |
| Riuzzi, 2007 [171] | TE671 | Musculoskeletal | Standard | flRAGE transfection, anti-HMGB1 (2.5 µg/mL) | Confirmed flRAGE transfection | RAGE expression decreased cell proliferation (p<0.01); blocking RAGE-HMGB1 with antibody increased cell proliferation (p<0.01) | N/A | RAGE transfection decreased apoptosis evasion (p<0.01) | N/A |
| Ruma, 2016 [153] | B16-BL6 | Skin | Standard | rS100A8/A9 (1000 ng/mL); wtRAGE, CT-RAGE | Confirmed RAGE expression, rS100A8/A9 bound to RAGE but didn't increase expression, confirmed transfections | wtRAGE transfection didn't affect cell proliferation (p>0.05); rS100A8/A9 and wtRAGE transfection mostly increased cell proliferation (p<0.001; >0.05) | wtRAGE transfection increased invasion (p<0.05); rS100A8/A9 and wtRAGE transfection somewhat increased invasion (p<0.01; >0.05); CT-RAGE decreased invasion with and without rS100A8/A9 (p<0.001) | N/A | wtRAGE transfection didn't affect NF-kB expression (p>0.05); rS100A8/A9 and wtRAGE transfection increased NF-kB expression (p<0.05; <0.01) |

| | | | | | | | | | |
|---|---|---|---|---|---|---|---|---|---|
| Ryan, 2019 [214] | Gem240, Gem257 | Breast | Standard | S100A4 + Gas6, FPS-ZM1 | Confirmed RAGE expression in cell lines, treated with RAGE ligand and inhibitor | N/A | S100A4 + Gas-6 increased invasion ($p<0.05$; 0.001); FPS-ZM1 negated S100A4/Gas6-induced invasion in Gem240 ($p<0.001$) but not in Gem257 ($p>0.05$) | N/A | N/A |
| Saha, 2010 [148] | B16F10 | Skin | Standard | S100A8, S100A9, anti-RAGE ab | Confirmed RAGE expression in B16F10 | N/A | Inhibition of RAGE by antibody negated S100A protein-induced migration ($p<0.05$) | N/A | N/A |
| Sajithlal, 2002 [182] | Neuro2a, SH-SY5Y | Brain | Standard | dnRAGE, anti-sense hRAGE, anti-sRAGE antibody | Confirmed RAGE expression, decrease with anti-sense hRAGE | dnRAGE, anti-sense-hRAGE, and anti-sRAGE decreased cell viability ($p<0.05$; 0.01) | N/A | N/A | N/A |
| Sakamoto, 2020 [114] | SAS, HSC-2, HSC-3 | Oral | Standard | FPS-ZM1 | Confirmed RAGE expression in SAS cell line, treated with RAGE inhibitor | FPS-ZM1 decreased cell viability in SAS cell line ($p<0.05$) but not HSC-2 or HSC-3 ($p>0.05$) | N/A | N/A | N/A |
| Sakurai, 2017 [144] | MCF-7, SK-BR-3, T47D. ZR-75-1 | Breast | Standard | SiS#1,#2 | Confirmed RAGE expression, decreased with SiS#1 and #2 except in SK-BR-3 | Silencing of RAGE ligands mostly decreased cell proliferation for all cell line ($p<0.05$; >0.05) except SK-BR-3 ($p>0.05$) | N/A | N/A | N/A |
| Santolla, 2022 [199] | MDA-MB-231 | Breast | CAF conditioned media | AGEs (100 µg/mL); FPS-ZM1 (1µM) | Confirmed RAGE expression, treated with RAGE ligand and inhibitor | N/A | AGEs induce migration and invasion ($p<0.05$); RAGE inhibition negated AGEs-induced migration and invasion ($p<0.05$) | N/A | N/A |
| Seguella, 2019 [195] | Caco-2 | Colorectum | Standard | S100B (5µM); RAGE mAb | Confirmed RAGE expression, increased with S100B, decreased with RAGE mAb | S100B increased cell proliferation ($p<0.001$); RAGE inhibition decreased S100B-induced cell proliferation ($p<0.05$) | S100B increased migration and invasion ($p<0.001$); RAGE inhibition decreased S100B-induced migration and invasion ($p<0.05$) | N/A | N/A |
| Seki, 2024 [145] | A172 | Brain | Standard | FPS-ZM1 (20µM); TTP488 (1µM); RAGE-KD | Confirmed RAGE expression, decreased with RAGE-KD, treated with RAGE inhibitors | RAGE inhibitors didn't affect cell viability ($p>0.05$) | RAGE inhibition and knockdown decreased migration ($p<0.05$; 0.01; 0.0001) | N/A | N/A |
| Sharaf, 2015 [115] | MDA-MB-231 | Breast | Standard | MG-BSA-AGEs (100 µg/mL; 200 µg/mL); anti-RAGE ab | Confirmed RAGE expression, increased with MG-BSA-AGEs | MG-BSA-AGEs didn't affect cell proliferation and viability ($p>0.05$); anti-RAGE ab decreased cell proliferation ($p<0.001$) | MG-BSA-AGEs didn't affect migratio and invasion ($p>0.05$); anti-RAGE ab decreased migration and invasion ($p<0.05$) | N/A | N/A |

| | | | | | | | | | |
|---|---|---|---|---|---|---|---|---|---|
| Shen, 2015 [139] | HCT116 | Colorectum | Standard | SOX9 KO, SOX+, SOX9+/S100P- | Confirmed RAGE expression, decreased with SOX9 KO and SOX9+/S100P-, increased with SOX9+ | N/A | SOX9 KO decreased migration and invasion (p<0.001); SOX9+ increased migration and invasion (p<0.001; =0.028); SOX9+/S100P- decreased migration and invasion (p<0.001; =0.003) | N/A | N/A |
| Shen, 2016 [116] | HCT116, LS174T | Colorectum | Standard | S100P-KD | Confirmed RAGE expression, decreased with S100P-KD | N/A | S100P knockdown reduced migration and invasion (p<0.05; 0.01) | N/A | N/A |
| Shu, 2025 [117] | HEC-1-B, Ishikawa | Endometrium | Standard | AGEs (200 µg/mL; 400 µg/mL); siRAGE | Confirmed RAGE expression, decreased with siRAGE, increased with AGEs | AGEs increased cell viability (p<0.001); RAGE silencing decreased AGEs-induced cell viability (p<0.05) | AGEs induced migration and invasion (p<0.001) | N/A | N/A |
| Siddique, 2013 [210] | LNCaP | Prostate | Standard | rhS100A4 (2 µg/mL); siRAGE | Confirmed RAGE expression, decreased with siRAGE, confirmed S100A4 bound to RAGE | RAGE silencing decreased cell proliferation (p<0.05) | N/A | N/A | RAGE silencing decreased NF-kB expression (p<0.05) |
| Swami, 2020 [178] | Panc-1 | Pancreas | Standard | oeRAGE, siRAGE, FPS-ZM1 (10µM) | Confirmed RAGE expression, decreased with siRAGE, treated with RAGE inhibitors and transfected with RAGE-overexpression | oeRAGE increased cell proliferation (p<0.001); RAGE silencing and inhibition decreased cell proliferation (p<0.001) | oeRAGE decreased migration and invasion (p<0.001) | N/A | oeRAGE decreased NF-kB expression (p<0.001) |
| Swanner, 2023 [118] | GBM6, GBM12, GBM28, GBM39, GBM59 | Brain | Standard | OVesRAGE | Confirmed OVesRAGE (decoy receptor) transfection | OVesRAGE mostly decreased cell viability (p<0.05; >0.05) | N/A | N/A | N/A |
| Taguchi, 2000 [119] | C6 | Brain | Standard | Amphoterin (10 µg/mL); flRAGE, tail deletion RAGE, sRAGE (100 µg/mL), anti-RAGE ab (100 µg/mL) | Confirmed RAGE expression in C6 cell line, treated with RAGE ligand, antibody, inhibitor, and RAGE-variants | flRAGE transfection with amphoterin increased cell proliferation (p<0.01); tail deletion RAGE and sRAGE decreased amphoterin-induced cell proliferation (p<0.0001; 0.00001) | flRAGE transfection with amphoterin increased invasion (p<0.00001); tail deletion RAGE, sRAGE, and anti-RAGE ab decreased amphoterin-induced invasion (p<0.00001) | N/A | N/A |
| Takamatsu, 2019 [120] | PK-8, Panc-1, AsPC-1, BxPC-3 | Pancreas | Standard | S100A11 (10 µg/mL) | Confirmed RAGE expression in all cell lines | S100A11 didn't affect cell proliferation (p>0.05) | N/A | N/A | N/A |
| Takeuchi, 2013 [150] | C6 | Brain | Standard | HMGB1 (0.1 µg/mL), flRAGE | Treated with RAGE ligand, transfected with RAGE | N/A | N/A | N/A | HMGB1 synergized with flRAGE to increase NF-kB expression (p<0.01) |

| | | | | | | | | | |
|---|---|---|---|---|---|---|---|---|---|
| Takeuchi, 2013 [150] | HT1080 | Musculoskeletal | Standard | HMGB1 (0.1 µg/mL), flRAGE, heparin (1 IU/mL) | Transfected RAGE to cell lines | RAGE transfection increased cell proliferation (p<0.01) | RAGE transfection increased migration and invasion (p<0.01); HMGB1 didn't impact RAGE-induced migration and invasion (p>0.05); heparin inhibited RAGE-induced migration and invasion (p<0.01) | N/A | N/A |
| Takino, 2010 [121] | A549 | Lungs | Standard | Glycer-AGEs (100 µg/mL) | Confirmed RAGE expression, treated with RAGE ligand | Glycer-AGEs decreased cell viability (p<0.01) | Glycer-AGEs increased migration and invasion (p<0.05; 0.01) | N/A | N/A |
| Talia, 2023 [201] | MCF-7, T47D | Breast | Standard | oeRAGE | Confirmed RAGE expression, increased with oeRAGE | oeRAGE increased cell proliferation (p<0.05) | oeRAGE increased migration and invasion (p<0.05) | N/A | N/A |
| Tang, 2012 [134] | JJ012, SW1353 | Musculoskeletal | Standard | HMGB1 (100 ng/mL), siRAGE | Confirmed RAGE expression, decreased with siRAGE | N/A | HMGB1 increased migration and invasion (p<0.05); siRAGE negated HMGB1-induced migration and invasion (p<0.05) | N/A | N/A |
| Tsuruhisa, 2021 [163] | MCF-7 | Breast | Standard | AGEs-BSA (100 µg/mL) | Confirmed RAGE expression, increased with AGEs-BSA | AGEs-BSA increased cell proliferation (p<0.01) | N/A | N/A | N/A |
| Wang, 2012 [138] | 95D | Lungs | Standard | siRAGE, anti-RAGE ab, CpG ODN | Confirmed RAGE expression, decreased with siRAGE, treated with RAGE antibody | RAGE silencing and anti-RAGE ab decreased cell proliferation (p<0.05) | RAGE silencing and anti-RAGE ab decreased invasion (p<0.05) | N/A | N/A |
| Wang, 2013 [194] | GL261 | Brain | Standard | oeS100B | Confirmed RAGE expression, increased with oeS100B | oeS100B didn't affect cell proliferation (p>0.05) | N/A | N/A | N/A |
| Wang, 2020 [69] | H1299 | Lungs | Standard | oeRAGE, siRAGE | Confirmed RAGE expression, increased with oeRAGE, decreased siRAGE | oeRAGE decreased cell proliferation (p<0.05); siRAGE increased cell proliferation (p<0.05) | oeRAGE decreased migration and invasion (p<0.05); siRAGE increased migration and invasion (p<0.05) | oeRAGE decreased apoptosis evasion (p<0.05); siRAGE increased apoptosis evasion (p<0.05) | n/A |
| Wang, 2021 [122] | LLC | Lungs | Standard | Dipyridamole (20 µM; 50 µM) | Confirmed RAGE expresion, decreased with dipyridamole | Dipyridamole decreased cell viability (p<0.05) | Dipyridamole decreased migration (p<0.05) | N/A | N/A |
| Wang, 2021 [70] | C666-1 | Nasopharynx | Standard | S100P (1000 ng/mL; 5000 ng/mL); FPS-ZM1 (1000 ng/mL; 5000 ng/mL) | Treated cell line with RAGE ligands and inhibitor | S100P increased cell proliferation (p<0.001); S100P didn't affect cell viability (p>0.05); FPS-ZM1 decreased cell proliferation and viability (p<0.001) | S100P increased migration (p<0.001); FPS-ZM1 decreased migration (p<0.001) | N/A | S100P increased NF-kB expression (p<0.05); FPS-ZM1 decreased NF-kB expression (p<0.001) |
| Wu, 2015 [192] | HepG2, SMMC-7721, Huh7 | Liver | Standard | GST-S100A9 (20 µg/mL); anti-RAGE ab (160 µg/mL) | Confirmed RAGE expression, increased with GST-S100A9, treated with anti-RAGE ab | Anti-RAGE ab decreased S100A9-induced cell proliferation and viability (p<0.05) | Anti-RAGE ab decreased S100A9-induced invasion (p<0.05; 0.01) | N/A | N/A |

| | | | | | | | | | |
|---|---|---|---|---|---|---|---|---|---|
| Wu, 2018 [223] | A498, ACHN | Kidneys | Standard | HMGB1-Vec, siHMGB1, sRAGE | Confirmed RAGE expression, decreased with siHMGB1 and sRAGE, increased with HMGB1-Vec | siHMGB1 and sRAGE decreased cell proliferation (p<0.01); HMGB1-Vec increased cell proliferation (p<0.05); HMGB1-Vec reduced sRAGE-induced cell proliferation decrease (p<0.05) | siHMGB1 and sRAGE decreased invasion (p<0.05; 0.01); HMGB1-Vec increased invasion (p<0.01); HMGB1-Vec reversed sRAGE-induced invasion decreased (p>0.05; 0.01) | siHMGB1 and sRAGE increased apoptosis (p<0.01); HMGB1-Vec didn't affect apoptosis (p>0.05); HMGB1-Vec reduced sRAGE-induced apoptosis (p<0.05) | N/A |
| Xu, 2013 [73] | SGC-7901 | Stomach | Standard | shRAGE | Confirmed RAGE expression, decreased with shRAGE | shRAGE decreased cell proliferation (p<0.01) | shRAGE decreased invasion (p<0.01) | shRAGE increased apoptosis (p<0.01) | N/A |
| Xu, 2014 [123] | Lewis | Lungs | Standard | HMGB1 (10 µg/mL); anti-RAGE ab (100 µg/mL) | Confirmed RAGE expression, increased with HMGB1, treated with antibody | HMGB1 increased cell proliferation (p<0.005); anti-RAGE ab negated HMGB1-induced cell proliferation (p<0.005) | N/A | N/A | N/A |
| Xu, 2019 [219] | HP(+) 7901, HP(+) MKN45 | Stomach | Standard | MiR-1915 mimic, pcDNA-RAGE | Confirmed RAGE expression, decreased with MiR-1915 mimic, regained expression with pcDNA-RAGE | MiR-1915 mimic decreased cell proliferation (p<0.05); pcDNA-RAGE negated MiR-1915 mimic-induced cell proliferation suppression (p<0.05) | MiR-1915 mimic decreased migration and invasion (p<0.05); pcDNA-RAGE mostly negated MiR-1915 mimic-induced migration and invasion (p<0.05; >0.05) | N/A | N/A |
| Yamamoto, 1996 [124] | MiaPaCa-2 | Pancreas | Standard | AGE-BSA (1 µg/mL) | Confirmed RAGE expression, increased with AGE-BSA | AGE-BSA increased cell viability (p<0.05) | N/A | N/A | N/A |
| Yamamoto, 2013 [125] | U87MG | Brain | Standard | siRAGE | Confirmed RAGE expression | N/A | RAGE silencing decreased migration (p<0.05) | N/A | N/A |
| Yamamoto, 2023 [165] | DLD-1 | Colorectum | Standard | Maple syrup protein fraction (MSpf; 10 µg/mL) | Confirmed RAGE expression, decreased with MSpf | MSpf decreased cell proliferation and viability (p<0.01; 0.001) | MSpf decreased migration and invasion (p<0.001) | N/A | N/A |
| Yang, 2015 [74] | SMMC-7721, HepG2 | Liver | Standard | siRAGE | Confirmed RAGE expression, treated with siRAGE | RAGE silencing reduced cell viability (p<0.01) | RAGE silencing reduced invasion (p<0.01) | N/A | N/A |
| Yang, 2024 [205] | 786-O, A498 | Kidneys | Standard | corylin (5µM; 40µM); 5-FU; oeRAGE | Confirmed RAGE expression, decreased with corylin, increased with oeRAGE | Corylin had no effect on cell viability (p>0.05); corylin increased cell proliferation (p<0.01); oeRAGE reversed corylin-induced cell proliferation (p<0.05) | Corylin decreased migration and invasion (p<0.01); oeRAGE reversed corylin-induced migration and invasion (p<0.05) | Corylin increased 5-FU-induced apoptosis (p<0.05) | N/A |
| Yaser, 2012 [76] | Huh7 | Liver | Standard | siRAGE | Confirmed RAGE expression, decreased with siRAGE | siRAGE decreased cell viability (p<0.01); siRAGE decreased cell proliferation (p<0.01) | N/A | N/A | RAGE silencing decreased NF-kB expression (p<0.01) |
| Ye, 2025 [126] | Panc-1, KPC-A548 | Pancreas | Standard | RAGEi, Sublethal PEF | Confirmed RAGE expression, increased with PEF, treated with RAGE inhibitor | RAGE inhibition decreased cell viability in KPC-A548 (p<0.05) and didn't affect cell viability in Panc-1 (p>0.05) | N/A | N/A | N/A |

| | | | | | | | | | |
|---|---|---|---|---|---|---|---|---|---|
| Yin, 2018 [77] | MDA-MB-231, MCF-7 | Breast | Standard | miR-185-5p, anti-miR-185-5p | Confirmed RAGE expression, decreased with miR-185-5p, increased with anti-miR-185-5p | N/A | miR-185-5p reduced invasion ($p<0.05$); anti-miR-185-5p increased migration ($p<0.05$) | N/A | N/A |
| Yu, 2017 [213] | H1975 | Lungs | Standard | siRAGE | Confirmed RAGE expression, decreased with siRAGE | siRAGE decreased cell viability and proliferation ($p<0.01$) | siRAGE decreased migration and invasion ($p<0.01$) | N/A | N/A |
| Yusein-Myashkova, 2025 [155] | MDA-MH-468, MDA-MB-231 | Breast | Standard | HMGB1 (800 ng/mL); metformin (1mM); esiRAGE | Confirmed RAGE expression, increased with HMGB1, decreased with esiRAGE, metformin negated HMGB1-induced increase | N/A | HMGB1 increased migration ($p<0.01$); metformin and esiRAGE negates HMGB1-induced migration ($p<0.05$; 0.01) | N/A | HMGB1 increased NF-kB expression ($p<0.01$); metformin negates HMGB1-induced NF-kB expression ($p<0.01$) |
| Zhan, 2012 [127] | BGC-823 | Stomach | Standard | siRAGE | Confirmed RAGE expression, decreased with siRAGE | N/A | N/A | RAGE silencing increased apoptosis ($p<0.01$) | N/A |
| Zhang, 2014 [212] | MG-63 | Musculoskeletal | Standard | SGP-2 (125nM; 250nM) | Confirmed RAGE expression, decreased with SGP-2 | SGP-2 decreased cell proliferation and viability ($p<0.01$) | SGP-2 decreased migration ($p<0.01$) | SGP-2 increased apoptosis ($p<0.01$) | N/A |
| Zhang, 2015 [128] | BGC-823 | Stomach | Normal media; gefitinib-conditioned media | HMGB1 (100 ng/mL), siRAGE | Confirmed RAGE expression, decreased with siRAGE, treated with RAGE ligand | RAGE silencing negated HMGB1-induced and gefitinib-conditioned media-induced cell proliferation ($p<0.01$) | N/A | N/A | N/A |
| Zhang, 2016 [78] | U2OS | Musculoskeletal | Standard | flRAGE transfection | Confirmed RAGE expression, increased with flRAGE transfection | flRAGE increased cell proliferation ($p<0.01$) | N/A | N/A | N/A |
| Zhang, 2018 [132] | PC-3 | Prostate | Standard | rHMGB1 (1 µg/mL); siRAGE | Confirmed RAGE expression, increased with rHMGB1, treated with RAGE inhibitor | RAGE silencing negated HMGB1-induced invasion ($p<0.0001$) | N/A | N/A | RAGE silencing negated HMGB1-induced NF-kB expression ($p<0.0001$) |
| Zhao, 2017 [181] | Tca-8113 | Oral | Standard | RAGE-ab (150 ng/mL); cisplatin (15µM) | Treated with anti-RAGE antibody | anti-RAGE ab decreased cell viability ($p<0.05$); anti-RAGE ab synergistically decreased cell viability with cisplatin ($p<0.05$) | N/A | N/A | N/A |
| Zheng, 2021 [129] | HCT116 | Colorectum | Standard | S100B (2 µg/mL); Apt-RAGE (100nM) | Confirmed RAGE expression, decreased with Apt-RAGE, increased with S100B | S100B increased cell proliferation ($p<0.01$); Apt-RAGE negated S100B-induced cell proliferation ($p<0.01$) | S100B increased migration ($p<0.01$); Apt-RAGE negated S100B-induced migration ($p<0.01$) | N/A | N/A |
| Zhou, 2024[130] | MDA-MB-231 | Breast | Standard | AGEs (100 µg/mL; 200 µg/mL); siRAGE | Confirmed RAGE expression, increased with AGEs, decreased with siRAGE | AGEs didn't affect cell viability ($p>0.05$) | AGEs increased migration ($p<0.01$); siRAGE decreased migration ($p<0.05$; 0.01); siRAGE negated AGEs-induced migration ($p<0.01$) | AGEs didn't affect apoptosis ($p>0.05$) | N/A |

| | | | | | | | | | |
|---|---|---|---|---|---|---|---|---|---|
| Zhu, 2015 [133] | HCT116, LoVo | Colorectum | Standard | rHMGB1 (2 µg/mL); siHMGB1, siRAGE | Confirmed RAGE expression, decreased with siRAGE, treated with RAGE ligand and RAGE ligand-silencing | N/A | rHMGB1 increased migration and invasion (p<0.05); siHMGB1 and siRAGE decreased rHMGB1-induced migration and invasion (p<0.05) | N/A | rHMGB1 increased NF-kB expression (p<0.05); siRAGE decreased rHMGB1-induced NF-kB expression (p<0.05) |
| Zhu, 2018 [131] | SiHa, CaSki | Cervix | Standard | LV-RAGE, siRNA1/2 | Confirmed RAGE expression, decreased with siRNA1 and 2, increased with LV-RAGE | LV-RAGE increased cell proliferation (p<0.05); siRNA1 and 2 decreased cell proliferation (p<0.05) | LV-RAGE increased migration (p<0.05); siRNA1 and 2 decreased migration (p<0.05) | LV-RAGE decreased apoptosis (p<0.05); siRNA1 and 2 increased apoptosis (p<0.05) | N/A |

*All studies reported standard incubator conditions (5% CO2, 37 °C) unless otherwise stated.

Abbreviations: advanced glycation end-products (AGE), receptor for advanced glycation end-products (RAGE), full length RAGE (flRAGE), RAGE-overexpression (RAGE-OE), soluble RAGE (sRAGE), human RAGE (hRAGE), wild type RAGE (wtRAGE), short hairpin RAGE (shRAGE), dominant negative RAGE (dnRAGE), silencing RAGE (siRAGE), long non-coding RAGE (lncRAGE), pLenti-C-mGFP-RAGE (LV-RAGE), endogenous secretory RAGE (esRAGE), high mobility group box 1 (HMGB1), RAGE-antagonistic peptide (RAP), N(6)-carboxymethyllysine (CML), bovine serum albumin (BSA), 4-acetylantroquinonol B (4-AAQB), aspirin (ASA), recombinant human APE1/Ref-1 (rhAPE1/Ref-1), gemcitabine (GEM), glyceraldehyde-AGEs (GA-AGEs), green fluorescent protein (GFP), notoginsenoside R1 (NGR1), methylgloxyal-BSA-AGEs (MG-BSA-AGEs), human serum albumin (HSA), lysophosphatidic acid (LPA), pulsed electric field (PEF), *S. glabra* polysaccharide 2 (SGP-2)
