## Supplemental Table 4 for "The Role of the Receptor for Advanced Glycation End-Products in Cancer: Evidence from a Systematic Review and Meta-Analysis"

| Author, Year | Source[1] | Experimental Conditions | | | | | Selective Reporting[7] | Statistics[8] | Total[9] |
|---|---|---|---|---|---|---|---|---|---|
| | | Culture Conditions[2] | Replicates[3] | Positive Controls[4] | Negative Controls[5] | Multiple Cell Lines[6] | | | |
| Abe, 2004 [184] | 1 | 1 | 1 | 0 | 1 | 0 | 1 | 1 | 6 |
| Alka, 2024 [207] | 1 | 1 | 1 | 1 | 1 | 1 | 1 | 1 | 8 |
| Amornsupak, 2022 [135] | 1 | 1 | 1 | 0 | 1 | 0 | 1 | 1 | 6 |
| Arrè, 2024 [141] | 1 | 1 | 1 | 0 | 1 | 1 | 1 | 1 | 7 |
| Arumugam, 2005 [196] | 1 | 1 | 1 | 0 | 1 | 0 | 1 | 1 | 6 |
| Arumugam, 2006 [81] | 1 | 1 | 1 | 1 | 1 | 1 | 1 | 1 | 8 |
| Arumugam, 2012 [197] | 1 | 1 | 1 | 1 | 0 | 1 | 1 | 1 | 7 |
| Bao, 2015 [82] | 1 | 1 | 1 | 1 | 0 | 0 | 0 | 1 | 5 |
| Bartling, 2005 [83] | 1 | 1 | 1 | 1 | 0 | 0 | 1 | 1 | 6 |
| Bassi, 2008 [84] | 1 | 1 | 1 | 1 | 0 | 0 | 1 | 1 | 6 |
| Bhawal, 2005 [85] | 1 | 1 | 1 | 1 | 0 | 1 | 0 | 1 | 6 |
| Chang, 2023 [161] | 1 | 1 | 1 | 0 | 1 | 1 | 1 | 1 | 7 |
| Chang, 2024 [162] | 1 | 1 | 1 | 1 | 1 | 1 | 1 | 1 | 8 |
| Chen, 2014 [86] | 1 | 1 | 1 | 1 | 1 | 0 | 1 | 1 | 7 |
| Chen, 2020 [170] | 1 | 1 | 1 | 0 | 1 | 0 | 1 | 1 | 6 |
| Chen, 2022 [87] | 1 | 1 | 1 | 0 | 1 | 0 | 1 | 1 | 6 |
| Chen, 2022 [151] | 1 | 1 | 1 | 0 | 1 | 0 | 1 | 1 | 6 |
| Chen, 2024 [186] | 1 | 1 | 1 | 0 | 1 | 1 | 1 | 1 | 7 |
| Cho, 2024 [191] | 1 | 1 | 1 | 1 | 0 | 0 | 1 | 1 | 6 |
| Choi, 2011 [142] | 1 | 1 | 1 | 0 | 1 | 0 | 1 | 1 | 6 |
| Chung, 2015 [88] | 1 | 1 | 1 | 0 | 1 | 0 | 1 | 1 | 6 |
| Dahlmann, 2014 [175] | 1 | 1 | 1 | 1 | 1 | 1 | 1 | 1 | 8 |
| Dai, 2023 [32] | 1 | 1 | 1 | 0 | 1 | 1 | 1 | 1 | 7 |
| Deng, 2017 [33] | 1 | 1 | 1 | 1 | 1 | 0 | 1 | 1 | 7 |
| Deng, 2017 [34] | 1 | 1 | 1 | 1 | 1 | 1 | 1 | 1 | 8 |
| Dhumale, 2015 [168] | 1 | 1 | 1 | 0 | 1 | 0 | 1 | 1 | 6 |
| Elangovan, 2012 [216] | 1 | 1 | 1 | 1 | 0 | 1 | 0 | 1 | 6 |
| El-Far, 2018 [140] | 1 | 1 | 1 | 0 | 1 | 1 | 1 | 1 | 7 |
| Fuentes, 2007 [169] | 1 | 1 | 1 | 1 | 1 | 0 | 1 | 1 | 7 |
| Gao, 2024 [89] | 1 | 1 | 1 | 1 | 0 | 1 | 1 | 1 | 7 |

| | | | | | | | | | |
|---|---|---|---|---|---|---|---|---|---|
| Geicu, 2020 [90] | 1 | 1 | 1 | 0 | 1 | 0 | 1 | 1 | 6 |
| Ghavami, 2008 [190] | 1 | 1 | 1 | 1 | 1 | 1 | 1 | 1 | 8 |
| Gurumayum, 2024 [159] | 1 | 1 | 1 | 0 | 1 | 0 | 1 | 1 | 6 |
| Guzmán, 2019 [206] | 1 | 1 | 1 | 0 | 1 | 1 | 1 | 1 | 7 |
| Han, 2023 [204] | 1 | 1 | 1 | 0 | 1 | 0 | 1 | 1 | 6 |
| He, 2018 [137] | 1 | 1 | 1 | 0 | 1 | 1 | 1 | 1 | 7 |
| Herwig, 2016 [91] | 1 | 1 | 1 | 1 | 1 | 0 | 1 | 1 | 7 |
| Herwig, 2016 [143] | 1 | 1 | 1 | 1 | 1 | 0 | 1 | 1 | 7 |
| Hou, 2018 [185] | 1 | 1 | 1 | 1 | 1 | 1 | 1 | 1 | 8 |
| Hsu, 2020 [166] | 1 | 1 | 1 | 0 | 1 | 0 | 1 | 1 | 6 |
| Huang, 2018 [136] | 1 | 1 | 1 | 0 | 1 | 0 | 1 | 1 | 6 |
| Huang, 2018 [218] | 1 | 1 | 1 | 0 | 1 | 0 | 1 | 1 | 6 |
| Huttunen, 2002 [183] | 1 | 1 | 1 | 0 | 1 | 0 | 1 | 1 | 6 |
| Ichikawa, 2011 [189] | 1 | 1 | 1 | 1 | 1 | 1 | 1 | 1 | 8 |
| Inada, 2019 [92] | 1 | 1 | 1 | 1 | 0 | 0 | 1 | 1 | 6 |
| Inada, 2019 [93] | 1 | 1 | 1 | 0 | 1 | 1 | 1 | 1 | 7 |
| Ishibashi, 2013 [154] | 1 | 1 | 1 | 1 | 1 | 0 | 1 | 1 | 7 |
| Ji, 2022 [222] | 1 | 1 | 1 | 1 | 1 | 1 | 1 | 1 | 8 |
| Jia, 2020 [157] | 1 | 1 | 1 | 0 | 1 | 0 | 1 | 1 | 6 |
| Jiang, 2025 [160] | 1 | 1 | 1 | 1 | 0 | 0 | 1 | 1 | 6 |
| Jin, 2011 [193] | 1 | 1 | 1 | 1 | 1 | 0 | 1 | 1 | 7 |
| Jin, 2025 [94] | 1 | 1 | 1 | 1 | 1 | 0 | 1 | 1 | 7 |
| Jing, 2015 [164] | 1 | 1 | 1 | 0 | 1 | 1 | 1 | 1 | 7 |
| Jube, 2012 [95] | 1 | 1 | 1 | 1 | 1 | 1 | 1 | 1 | 8 |
| Kang, 2010 [208] | 1 | 1 | 1 | 0 | 1 | 1 | 1 | 1 | 7 |
| Kang, 2011 [209] | 1 | 1 | 1 | 0 | 1 | 1 | 1 | 1 | 7 |
| Kang, 2012 [202] | 1 | 1 | 1 | 0 | 1 | 0 | 1 | 1 | 6 |
| Kang, 2014 [174] | 1 | 1 | 1 | 0 | 1 | 0 | 1 | 1 | 6 |
| Kang, 2014 [203] | 1 | 1 | 1 | 0 | 1 | 1 | 1 | 1 | 7 |
| Kataoka, 2012 [187] | 1 | 1 | 1 | 1 | 1 | 1 | 1 | 1 | 8 |
| Khoo, 2023 [45] | 1 | 1 | 1 | 1 | 0 | 1 | 1 | 1 | 7 |
| Kim, 2025 [96] | 1 | 1 | 1 | 0 | 1 | 1 | 1 | 1 | 7 |
| Kinoshita, 2019 [97] | 1 | 1 | 1 | 1 | 0 | 0 | 1 | 1 | 6 |
| Ko, 2014 [98] | 1 | 1 | 1 | 0 | 1 | 0 | 0 | 1 | 5 |

| | | | | | | | | | |
|---|---|---|---|---|---|---|---|---|---|
| Kobayashi, 2007 [47] | 1 | 1 | 1 | 0 | 1 | 1 | 1 | 1 | 7 |
| Kuniyasu, 2002 [99] | 1 | 1 | 1 | 1 | 0 | 0 | 1 | 1 | 6 |
| Kuniyasu, 2003 [100] | 1 | 1 | 1 | 1 | 0 | 1 | 0 | 1 | 6 |
| Kwak, 2017 [215] | 1 | 1 | 1 | 0 | 1 | 1 | 1 | 0 | 6 |
| Lai, 2021 [101] | 1 | 1 | 1 | 0 | 1 | 1 | 1 | 1 | 7 |
| Lan, 2019 [167] | 1 | 1 | 1 | 0 | 1 | 0 | 1 | 1 | 6 |
| Lata, 2014 [147] | 1 | 1 | 1 | 1 | 1 | 0 | 0 | 0 | 5 |
| Lee, 2015 [102] | 1 | 1 | 1 | 1 | 0 | 0 | 1 | 1 | 6 |
| Lee, 2018 [217] | 1 | 1 | 1 | 1 | 0 | 0 | 1 | 1 | 6 |
| Li, 2012 [103] | 1 | 1 | 1 | 0 | 1 | 1 | 1 | 1 | 7 |
| Li, 2018 [172] | 1 | 1 | 1 | 0 | 1 | 1 | 0 | 1 | 6 |
| Li, 2020 [104] | 1 | 1 | 1 | 0 | 1 | 1 | 1 | 1 | 7 |
| Li, 2020 [211] | 1 | 1 | 1 | 0 | 1 | 1 | 1 | 1 | 7 |
| Li, 2024 [51] | 1 | 1 | 1 | 0 | 1 | 0 | 1 | 1 | 6 |
| Li, 2024 [224] | 1 | 1 | 1 | 0 | 1 | 1 | 1 | 1 | 7 |
| Liang, 2011 [146] | 1 | 1 | 1 | 0 | 1 | 0 | 1 | 1 | 6 |
| Liao, 2018 [105] | 1 | 1 | 1 | 1 | 0 | 0 | 1 | 1 | 6 |
| Liao, 2023 [117] | 1 | 1 | 1 | 0 | 1 | 0 | 1 | 1 | 6 |
| Lin, 2012 [53] | 1 | 1 | 1 | 0 | 1 | 0 | 1 | 1 | 6 |
| Lin, 2020 [221] | 1 | 1 | 1 | 0 | 1 | 0 | 1 | 1 | 6 |
| Liu, 2025 [106] | 1 | 1 | 1 | 0 | 1 | 0 | 1 | 1 | 6 |
| Madhavan, 2021 [107] | 1 | 1 | 1 | 0 | 1 | 0 | 1 | 1 | 6 |
| Magna, 2023 [173] | 1 | 1 | 1 | 0 | 1 | 1 | 1 | 1 | 7 |
| Matou-Nasri, 2017 [108] | 1 | 1 | 1 | 0 | 1 | 0 | 1 | 1 | 6 |
| Medapati, 2015 [56] | 1 | 1 | 1 | 0 | 1 | 0 | 1 | 1 | 6 |
| Meghnani, 2014 [176] | 1 | 1 | 1 | 0 | 1 | 0 | 1 | 1 | 6 |
| Méndez, 2018 [109] | 1 | 1 | 1 | 1 | 1 | 1 | 0 | 1 | 7 |
| Menini, 2018 [198] | 1 | 1 | 1 | 1 | 0 | 0 | 1 | 1 | 6 |
| Mitsui, 2019 [220] | 1 | 1 | 1 | 0 | 1 | 0 | 1 | 1 | 6 |
| Nakamara, 2017 [179] | 1 | 1 | 1 | 1 | 1 | 0 | 1 | 1 | 7 |
| Nam, 2021 [156] | 1 | 1 | 1 | 0 | 1 | 1 | 1 | 1 | 7 |
| Ojima, 2014 [110] | 1 | 1 | 1 | 1 | 1 | 0 | 1 | 1 | 7 |
| Pan, 2018 [149] | 1 | 1 | 1 | 0 | 1 | 1 | 1 | 1 | 7 |
| Pan, 2022 [111] | 1 | 1 | 1 | 0 | 1 | 0 | 1 | 1 | 6 |

| | | | | | | | | | |
|---|---|---|---|---|---|---|---|---|---|
| Popa, 2014 [112] | 1 | 1 | 1 | 0 | 1 | 1 | 1 | 0 | 6 |
| Pujals, 2023 [180] | 1 | 1 | 1 | 0 | 1 | 1 | 1 | 1 | 7 |
| Qian, 2019 [66] | 1 | 1 | 0 | 1 | 1 | 1 | 0 | 0 | 5 |
| Qiao, 2016 [113] | 1 | 1 | 1 | 0 | 1 | 1 | 1 | 1 | 7 |
| Ray, 2020 [152] | 1 | 1 | 1 | 1 | 1 | 1 | 1 | 1 | 8 |
| Reeb, 2015 [188] | 1 | 1 | 1 | 0 | 1 | 1 | 1 | 1 | 7 |
| Rehbein, 2008 [158] | 1 | 1 | 1 | 1 | 0 | 0 | 1 | 1 | 6 |
| Ren, 2021 [200] | 1 | 1 | 1 | 0 | 1 | 0 | 0 | 1 | 5 |
| Riuzzi, 2007 [171] | 1 | 1 | 1 | 0 | 1 | 0 | 1 | 1 | 6 |
| Ruma, 2016 [153] | 1 | 1 | 1 | 1 | 1 | 0 | 1 | 1 | 7 |
| Ryan, 2019 [214] | 0 | 1 | 1 | 1 | 1 | 1 | 1 | 1 | 7 |
| Saha, 2010 [148] | 1 | 1 | 1 | 1 | 1 | 0 | 1 | 0 | 6 |
| Sajithlal, 2002 [182] | 1 | 1 | 1 | 0 | 1 | 1 | 1 | 0 | 6 |
| Sakamoto, 2020 [114] | 1 | 1 | 1 | 0 | 1 | 1 | 0 | 1 | 6 |
| Sakurai, 2017 [144] | 1 | 1 | 1 | 0 | 1 | 1 | 1 | 1 | 7 |
| Santolla, 2022 [199] | 1 | 1 | 1 | 1 | 1 | 0 | 1 | 1 | 7 |
| Seguella, 2019 [195] | 1 | 1 | 1 | 1 | 1 | 0 | 1 | 1 | 7 |
| Seki, 2024 [145] | 1 | 1 | 1 | 0 | 1 | 0 | 1 | 1 | 6 |
| Sharaf, 2015 [115] | 1 | 1 | 1 | 1 | 1 | 0 | 1 | 1 | 7 |
| Shen, 2015 [139] | 1 | 1 | 1 | 0 | 1 | 0 | 0 | 1 | 5 |
| Shen, 2016 [116] | 1 | 1 | 1 | 0 | 1 | 1 | 0 | 1 | 6 |
| Shu, 2025 [117] | 1 | 1 | 1 | 1 | 1 | 1 | 1 | 1 | 8 |
| Siddique, 2013 [210] | 1 | 1 | 1 | 1 | 0 | 0 | 1 | 1 | 6 |
| Swami, 2020 [178] | 1 | 1 | 1 | 0 | 1 | 0 | 1 | 1 | 6 |
| Swanner, 2023 [118] | 1 | 1 | 1 | 0 | 1 | 1 | 1 | 1 | 7 |
| Taguchi, 2000 [119] | 1 | 1 | 1 | 1 | 0 | 0 | 1 | 1 | 6 |
| Takamatsu, 2019 [120] | 1 | 1 | 1 | 0 | 1 | 1 | 1 | 1 | 7 |
| Takeuchi, 2013 [150] | 1 | 1 | 1 | 1 | 1 | 1 | 1 | 1 | 8 |
| Takino, 2010 [121] | 1 | 1 | 1 | 0 | 1 | 0 | 1 | 1 | 6 |
| Talia, 2023 [201] | 1 | 1 | 1 | 0 | 1 | 1 | 1 | 1 | 7 |
| Tang, 2012 [134] | 1 | 1 | 1 | 1 | 1 | 0 | 1 | 1 | 7 |
| Tsuruhisa, 2021 [163] | 1 | 1 | 1 | 0 | 1 | 0 | 1 | 1 | 6 |
| Wang, 2012 [138] | 1 | 1 | 1 | 0 | 1 | 0 | 1 | 1 | 6 |
| Wang, 2013 [194] | 1 | 1 | 1 | 0 | 1 | 0 | 1 | 1 | 6 |

| | | | | | | | | | |
|---|---|---|---|---|---|---|---|---|---|
| Wang, 2020 [69] | 1 | 1 | 1 | 0 | 1 | 0 | 1 | 1 | 6 |
| Wang, 2021 [122] | 1 | 1 | 1 | 0 | 1 | 0 | 1 | 1 | 6 |
| Wang, 2021 [70] | 1 | 1 | 1 | 0 | 1 | 0 | 1 | 1 | 6 |
| Wu, 2015 [192] | 1 | 1 | 1 | 1 | 0 | 1 | 0 | 1 | 6 |
| Wu, 2018 [223] | 1 | 1 | 1 | 1 | 1 | 1 | 1 | 1 | 8 |
| Xu, 2013 [73] | 1 | 1 | 1 | 0 | 1 | 0 | 1 | 1 | 6 |
| Xu, 2014 [123] | 1 | 1 | 1 | 1 | 1 | 0 | 1 | 1 | 7 |
| Xu, 2019 [219] | 1 | 1 | 1 | 1 | 1 | 1 | 1 | 1 | 8 |
| Yamamoto, 1996 [124] | 1 | 1 | 1 | 0 | 1 | 0 | 1 | 1 | 6 |
| Yamamoto, 2013 [125] | 1 | 1 | 1 | 0 | 1 | 0 | 1 | 1 | 6 |
| Yamamoto, 2023 [165] | 1 | 1 | 1 | 0 | 1 | 0 | 1 | 1 | 6 |
| Yang, 2015 [74] | 1 | 1 | 1 | 0 | 1 | 1 | 1 | 1 | 7 |
| Yang, 2024 [205] | 1 | 1 | 1 | 1 | 1 | 1 | 1 | 1 | 8 |
| Yaser, 2012 [76] | 1 | 1 | 1 | 0 | 1 | 0 | 1 | 1 | 6 |
| Ye, 2025 [126] | 1 | 1 | 1 | 0 | 1 | 1 | 1 | 1 | 7 |
| Yin, 2018 [77] | 1 | 1 | 1 | 0 | 1 | 1 | 1 | 1 | 7 |
| Yu, 2017 [213] | 1 | 1 | 1 | 0 | 1 | 0 | 1 | 1 | 6 |
| Yusein-Myashkova, 2025 [155] | 1 | 1 | 1 | 1 | 1 | 1 | 1 | 1 | 8 |
| Zhan, 2012 [127] | 1 | 1 | 1 | 0 | 1 | 0 | 1 | 1 | 6 |
| Zhang, 2014 [212] | 1 | 1 | 1 | 0 | 1 | 0 | 1 | 1 | 6 |
| Zhang, 2015 [128] | 1 | 1 | 1 | 1 | 0 | 0 | 1 | 1 | 6 |
| Zhang, 2016 [78] | 1 | 1 | 1 | 0 | 1 | 0 | 1 | 1 | 6 |
| Zhang, 2018 [132] | 1 | 1 | 1 | 1 | 0 | 0 | 1 | 1 | 6 |
| Zhao, 2017 [181] | 1 | 1 | 1 | 0 | 1 | 0 | 0 | 1 | 5 |
| Zheng, 2021 [129] | 1 | 1 | 1 | 1 | 1 | 0 | 0 | 1 | 6 |
| Zhou, 2024 [130] | 1 | 1 | 1 | 1 | 1 | 0 | 1 | 1 | 7 |
| Zhu, 2015 [133] | 1 | 1 | 1 | 1 | 1 | 1 | 1 | 1 | 8 |
| Zhu, 2018 [131] | 1 | 1 | 1 | 0 | 1 | 1 | 1 | 1 | 7 |

Studies are given a score of 0 or 1 for each of the following parameters (a score of 0 was given for a lack of criteria fulfillment or failure to report): 1cell lines are independently validated; 2comparable culture conditions to other studies; 3experiment performed in replicate(s); 4appropriate positive controls included; 5appropriate negative controls included; 6more than one cell line used; 7all experimental results are reported (a score of 0 was given for missing statistical data); 8appropriate statistical test used.
