## Supplemental Table 5 for "The Role of the Receptor for Advanced Glycation End-Products in Cancer: Evidence from a Systematic Review and Meta-Analysis"

| Author, Year | Animal Model | Organ of Origin | Cell line(s) used | Established Cell Line? | Treatment/ Dose(s) | RAGE Expression | Length of study/ intervention | Tumor Growth | Metastatic Growth |
|---|---|---|---|---|---|---|---|---|---|
| Abe, 2004 [184] | Subcutaneous xenograft on Balb/c-nu/nu mice | Skin | G361 | Y | anti-RAGE ab (IP; 0.5 mg/5d; 0.5mg/3d) | Confirmed RAGE expression in cell line, treated with anti-RAGE therapy | 30d for tumor growth, 3w for metastasis | RAGE inhibition reduced tumor volume (p<0.01) | RAGE inhibition reduced lung metastasis (p<0.01) |
| Alka, 2024 [207] | Subcutaneous xenograft on female mice (C57BL/6NJ or NU/J) | Pancreas | Panc1, Panc02 | Y | Azeliragon (IP; 1mg/kg/d) | Confirmed RAGE expression in cell lines, treated with RAGE inhibitor | 34d | RAGE inhibition reduced tumor volume (p<0.05) | N/A |
| Arumugam, 2005 [196] | Subcutaneous xenograft on male athymic nu/nu mice | Pancreas | Panc1, BxPC-3 | Y | S100P-OV, siS100P | Confirmed S100P binding to RAGE | 4w | S100P-OV increased tumor volume (p<0.05); siS100P decreased tumor bioluminescence (p<0.05) | N/A |
| Arumugam, 2006 [81] | Orthotopic xenograft on male CB17 SCID mice | Pancreas | BxPC-3, Mpanc-96, Panc1 | Y | Cromolyn (IP; 5mg/kg/d) | Confirmed RAGE expression in cell lines, cromolyn decreased RAGE expression | 6w | Cromolyn reduced tumor bioluminescence in BxPC-3 and Mpanc-96 (p=0.009; <0.001) but Panc1 didn't affect tumor bioluminescence (p>0.05) | Cromolyn didn't impact liver metastasis in BxPC-3 and Panc1 and lung metastasis in Panc1 (p>0.05); cromolyn reduced lung metastasis in BxPC-3 and Mpanc-96 and reduced liver metastasis in Mpanc-96 (p<0.01; =0.013; =0.017) |
| Arumugam, 2012 [197] | Subcutaneous xenograft on male athymic nu/nu mice | Brain | C6 | Y | RAGE-antagonistic peptide (RAP) (IP; 100 µg/d) | Treated with RAGE inhibitor | 3w | RAP reduced tumor bioluminescence (p<0.05) | N/A |
| Arumugam, 2012 [197] | Orthotopic xenograft on male athymic nu/nu mice | Pancreas | MPanc96 | Y | RAP (IP; 100 µg/d) | Treated with RAGE inhibitor | 3w | RAP reduced tumor bioluminescence (p<0.05) | RAP reduced liver metastasis bioluminescence (p<0.05) |
| Arumugam, 2013 [230] | Orthotopic xenograft on male C57BL/6J mice | Pancreas | k-ras-p53KO transgenic mouse tumor-derived cells | N | C5OH (IP; 5mg/d) | C5OH blocks RAGE bindining | 5w | C5OH inhibited tumor weight (p<0.05) | C5OH inhibited liver metastasis (p<0.05) |
| Arumugam, 2013 [230] | Orthotopic xenograft on male athymic nude mice | Pancreas | Mpanc-96 | Y | C5OH (IP; 5mg/d) | C5OH blocks RAGE bindining | 5w | C5OH inhibited tumor volume (p<0.05) | N/A |
| Bartling, 2005 [83] | Subcutaneous xenograft on female athymic NMRI-nu mice | Lungs | NCI-H358 | Y | dnRAGE transfection | Confirmed RAGE expression in cell line, decreased with dnRAGE | 29d | RAGE inhibition increased tumor volume and weight (p<0.05) | N/A |

| | | | | | | | | | |
|---|---|---|---|---|---|---|---|---|---|
| Chen, 2014 [244] | Orthotopic xenograft on C57BL/6J AGER-KO mice | Brain | GL261, K-Luc | Y | RAGE-KO transfection | Confirmed RAGE expression in cell lines, decreased with RAGE KO | 18d | RAGE knockdown reduced tumor bioluminescence in GL261 xenograft ($p<0.05$); RAGE knockdown didn't affect the tumor area in either cell line and the tumor bioluminescence in K-Luc ($p>0.05$) | N/A |
| Chen, 2020 [170] | Subcutaneous xenograft on male BALB/c-nu/nu mice | Lungs | A549 | Y | oeRAGE transfection | Confirmed no RAGE expression in parental cell line and increased with oeRAGE | 10w | oeRAGE didn't increase tumor volume ($p>0.05$) | oeRAGE didn't affect lung metastasis ($p>0.05$) |
| Chen, 2021 [227] | Orthotopic xenograft on male BALB/c-nu/nu mice | Pancreas | MiaPaCa-2-Luc | Y | Pterostilbene (Oral gavage; 500mg/kg/d) + Chloroquine (oral gavage; 10mg/kg/d) | Confirmed RAGE expression, decreased with pterostilbene/chloroquine treatment | 28d | PT+CQ decreased tumor weight and bioluminescence ($p<0.05$) | N/A |
| Chen, 2024 [186] | Popliteal lymph node metastasis model on BALB/c-nu/nu mice | Liver | H22 | Y | oeS100A6; shS100A6 | Treated with RAGE ligand or silencing RAGE ligand | Not specified | N/A | shS100A6 reduced lymph node weight ($p<0.01$); oeS100A6 increased lymph node weight ($p<0.001$) |
| Chen, 2024 [251] | Induced oral carcinogenesis by 4NQO on male C57BL/6J mice | Oral | N/A | N/A | anti-RAGE ab (IP; 100 µg/inj, 4 times) | Treated with anti-RAGE antibody | 15d | anti-RAGE antibody decreased number of small tumors <1 mm ($p<0.05$) but not number of larger tumors >1 mm ($p>0.05$) | N/A |
| Cheng, 2014 [232] | Orthotopic xenograft on male BALB/c-nu/nu mice | Liver | HCC-LM3 | Y | Ethyl pyruvate (IP; 80 mg/kg/d) | Confirmed RAGE expression in cell line, decreased with ethyl pyruvate treatment | 4w | Ethyl pyruvate decreased tumor diameter and volume ($p<0.05$) | N/A |
| Chiappalupi, 2014 [229] | Intraperitoneal xenograft on female athymic nu/nu mice | Musculoskeletal | TE671 | Y | flRAGE transfection | Confirmed low RAGE expression in cell line, increased with flRAGE transfection | 6w | flRAGE decreased tumorigenesis ($p<0.01$) | flRAGE decreased metastasis ($p<0.05$) and lung metastasis ($p<0.01$; 0.001) |
| Cho, 2024 [228] | AOM/DSS-induced colorectal tumors in female C57BL/6 mice | Colorectum | N/A | N/A | rCT-S100A8/9 (IP; 50 µg/kg; 3x 10w) | Confirmed RAGE expression, rCT-S100A8/9 blocked S100A8/9-RAGE binding | 10w | rCT-S100A8/9 decreased number of tumors <2mm ($p<0.05$) | N/A |
| Cho, 2024 [228] | Subcutaneous xenograft on female athymic nu/nu mice | Colorectum | HCT116 | Y | rCT-S100A8/9 (IP; 50 µg/kg; 30d) | Confirmed RAGE expression, rCT-S100A8/9 blocked S100A8/9-RAGE binding | 30d | rCT-S100A8/9 decreased tumor volume ($p<0.001$) | N/A |
| Dai, 2023 [32] | Subcutaneous xenograft on female BALB/c nu/nu mice | Breast | MDA-MB-231; BT549 | Y | RP7 (IP; 30 mg/kg) | Confirmed RAGE expression in cell lines, decreased with RAGE inhibitor | 22d | RP7 decreased tumor volume and weight ($p<0.001$) | N/A |

| | | | | | | | | | |
|---|---|---|---|---|---|---|---|---|---|
| DiNorcia, 2010 [245] | Apc-1638N/+ transgenic colorectal mouse model | Intestinal | N/A | N/A | RAGE-/- strain | Crossed mice with RAGE -/- strain | 30w | RAGE -/- mice didn't affect mean and median tumor count (p=0.31; >0.05), decreased mean/median tumor diameter and incidence (p<0.001; 0.01; 0.03) | n/A |
| DiNorcia, 2010 [245] | Intrahepatic injection in BALB/c or C57BL/6 mice | Colorectum | CT26; MC38 | Y | sRAGE (100 µg/d); RAGE-/- strain; flRAGE transfection | Crossed mice with RAGE -/- strain, transfected with flRAGE, treated with RAGE inhibitor | 28d | N/A | sRAGE mostly reduced liver metastasis (p=0.45; <0.05; 0.01); RAGE -/- decreased liver metastasis (p<0.03; 0.01); flRAGE increased liver metastasis (p<0.02) |
| Elangovan, 2012 [216] | Subcutaneous xenograft in male nude mice | Prostate | LNCaP, DU145 | Y | shRAGE (IP; 100 µg/5x 6w) | Confirmed RAGE expression in cell lines, decreased with shRAGE | 6w | shRAGE decreased tumor volume (p<0.05) | N/A |
| Fan, 2024 [239] | Oncogenic plasmid by hydrodynamic tail vein injection | Liver | N/A | N/A | Hi-AGE diet; ALT-711 (1 mg/kg/d) | Treated mice with RAGE ligand diet and RAGE inhibition | 7w | Hi AGE diet increased tumor count (p<0.0001); ALT-711 negated Hi AGE diet decreased tumor count (p<0.0001) | N/A |
| Heijmans, 2013 [246] | Sporadic intestinal tumorigenic model with APC-min/+ mice | Intestinal | N/A | N/A | RAGE-/- strain | Crossed mice with RAGE -/- strain | 17w | RAGE -/- mice decreased adenoma polyp number (p<0.01) | N/A |
| Hernandez, 2013 [253] | Subcutaneous xenograft on Hsd:athymic nude Foxn1nu mice | Skin | M21 | Y | S100A4 transfection; 5C3 (IP; 25 mg/kg/w) | Confirmed RAGE expression in cell line, blocked S100A4-RAGE with 5C3 | 44d or 17d | S100A4 increased tumor weight (p<0.05); 5C3 decreased tumor weight (p<0.05) | N/A |
| Hernandez, 2013 [253] | Subcutaneous xenograft on Hsd:athymic nude Foxn1nu mice | Pancreas | MiaPaCa-2 | Y | siS100A4; 5C3 (IP; 25 mg/kg/w) | Confirmed RAGE expression in cell line, blocked S100A4-RAGE with 5C3 | 62d or 17d | siS100A4 and 5C3 decreased tumor weight (p<0.05) | N/A |
| Hiramoto, 2021 [234] | Tail vein injection on male C57BL/6J mice | Skin | B16 | Y | Glycyrrhizin (IP; 15 mg/kg/2d) | Confirmed RAGE expression in lungs with melanoma metastasis, decreased with Glycyrrhizin | 2w | N/A | Glycyrrhizin decreased lung metastases area (p<0.05) |
| Hou, 2025 [225] | Orthotopic xenograft on male C57BL/6 mice | Liver | Hepa1-6 | Y | 5OMV (IP; 50 mg/kg/d) | Confirmed RAGE expression in cell line, decreased with 5OMV | 14d | 5OMV decreased tumor weight (p<0.01) | N/A |
| Huttunen, 2002 [183] | Tail vein injection on female C57BL/6-scid-beige mice | Skin | B16-F1 | Y | dnRAGE transfection, amphoterin (IP; 500µM) | Confirmed RAGE expression, transfected with dnRAGE, treated with RAGE inhibitor | 2w | N/A | dnRAGE and amphoterin decrreased lung metastasis (p<0.0001) |

| | | | | | | | | | |
|---|---|---|---|---|---|---|---|---|---|
| Ichikawa, 2011 [189] | AOM/DSS-induced colorectal tumors in female C57BL/6 mice | Colorectum | N/A | N/A | S100A9 KO | Confirmed RAGE-S100A9 interaction | 12; 20w | S100A9 KO decreased tumor count (p<0.05) | N/A |
| Ichikawa, 2011 [189] | Subcutaneous xenograft on C57BL/6 mice | Colorectum | MC38 | Y | S100A9 KO | Confirmed RAGE expression in cell line and RAGE-S100A9 interaction | 3w | S100A9 KO decreased tumor volume (p<0.01) | N/A |
| Ichikawa, 2011 [189] | Under spleen injection on C57BL/6 mice | Colorectum | MC38 | Y | S100A9 KO | Confirmed RAGE expression in cell line and RAGE-S100A9 interaction | 2w | N/A | S100A9 KO decreased metastasis count and tumor burden (p<0.05) |
| Jiang, 2025 [160] | Subcutaneous xenograft on female BALB/c mice | Breast | 4T1 | Y | NanoPt @ Cas9-PAR2 | Confirmed RAGE expression, decreased with NanoPt @ Cas9-PAR2 | 15d | NanoPt @ Cas9-PAR2 decreased tumor volume (p<0.05) | N/A |
| Jing, 2015 [164] | Subcutaneous xenograft or tail vein injection on Balb/c-nu/nu mice | Esophagus | Eca-109 | Y | miR-185 | Confirmed RAGE expression in cell line, decreased with miR-185 | 3w | miR-185 didn't impact tumor volume and weight (p=0.8; 0.097) | miR-185 decreased lung metastasis (p<0.003) |
| Kang, 2010 [208] | Subcutaneous xenograft on C57/Bl6 mice | Pancreas | Panc02 | Y | shRAGE | Confirmed RAGE expression in cell line, decreased with shRAGE | 38d | RAGE knockdown didn't affect tumor volume (p>0.05) | N/A |
| Kang, 2014 [203] | Subcutaneous allograft on C57/Bl6 mice | Pancreas | Panc02 | Y | shRAGE, RAGE -/- strain | Confirmed RAGE expression in cell line, decreased with shRAGE, crossed mice with RAGE -/- strain | 42d | shRAGE and RAGE KO decreased tumor volume (p<0.001) | N/A |
| Khoo, 2023 [45] | Subcutaneous xenograft on NOD/SCID mice | Prostate | DU145 | Y | shRAGE | Confirmed RAGE expression, decreased with shRAGE | 4w | shRAGE decreased tumor bioluminescence (p<0.05) | N/A |
| Kim, 2025 [96] | Xenograft on NOD/SCID mice | Lungs | NCI-H596 | Y | HMGB1 (IP; 200 mg/kg/18x) | Confirmed RAGE expression in cell line, decreased with HMGB1 | 38d | HMGB1 decreased tumor volume and weight (p<0.0001) | N/A |
| Krisanits, 2022 [242] | Subcutaneous allograft on C57/Bl6 mice | Prostate | MYC-CaP | Y | Hi Glycated diet, AGE pre-treatment, RAGE KO | Confirmed RAGE expression, increased Hi-Glyc diet and AGE pretreatment, decreased with RAGE KO, | 24d, 60d | AGE pretreatment and hi-glycated diet increased tumor volume (p<0.0031); RAGE KO decreased tumor volume (p<0.0001) | N/A |
| Kwak, 2017 [215] | Orthotopic xenograft on female NSG, BALBc, or C57BL6 mice | Breast | MDA-MD-231, MDA-MB-4175, 4T1, AT3 | Y | oeRAGE, RAGE (sh10, sh12, sh66), FPS-ZM1 (IP; 1 mg/kg/2x w) | Confirmed RAGE expression in cell lines, decreased with shRAGEs, increased with oeRAGE, treated with RAGE inhibitor | 35d | RAGE modulation in MDA-MB-231 xenografts didn't affect tumor growth (p>0.05); RAGE reduction in all other xenografts decreased tumor weight and volume (p<0.05) | N/A |
| Lee, 2018 [217] | Orthotopic xenograft on female BALB/c mice | Breast | MDA-MB-231 | Y | ASA (oral; 20 mg/kg/d); TSA (SC; 0.5 mg/kg/3d); RAGE-KD transfection | Confirmed RAGE expression in cell line, decreased with RAGE KD | 13w | RAGE knockdown decreased tumor volume and weight (p<0.01) | N/A |

| | | | | | | | | | |
|---|---|---|---|---|---|---|---|---|---|
| Li, 2018 [172] | Subcutaneous xenograft on male BALB/c-nu/nu mice | Liver | HCCLM3 | Y | sorafenib (IP; 50 mg/kg/d), shRAGE | Confirmed RAGE expression in cell line, treated with RAGE silencing | 4w | RAGE silencing increased sorafenib reduction of tumor volume (p<0.05) | N/A |
| Li, 2021 [254] | Subcutaneous xenograft on female BALB/c nu/nu mice | Pancreas | MiaPaCa-2 | Y | Ursolic acid (IP; 40 mg/kg/8x) | Confirmed RAGE expression in cell line, decreased with UA | 22d | UA decreased tumor volume (p<0.05) | N/A |
| Li, 2024 [241] | Subcutaneous xenograft on nude or NSG mice | Colon | HCT116, DLD1, RKO | Y | RAGE KO 1/2 | Confirmed RAGE expression in cell line, decreased with knockout transfection | 28d | RAGE KOs didn't affect tumor volume (p>0.05) | N/A |
| Mark, 2013 [249] | Induced oral and esophageal carcinogenesis by 4NQO on male C57BL/6 mice | Oral and Esophageal | N/A | N/A | RAGE-/- strain | Confirmed RAGE expression, crossed with RAGE-/- strain | 25w | RAGE KO didn't affect tumor count (p=0.66; 0.6) | N/A |
| Meghnani, 2014 [248] | Subcutaneous xenograft on female SCID mice | Skin | WM115 | Y | flRAGE transfection, anti-RAGE IgG2A11 (0.5 mg/5d) | Used flRAGE transfection and anti-RAGE antibody | 28d | flRAGE increased tumor volume (p<0.001); anti-RAGE IgG2A11 decreased tumor volume (p<0.01) | N/A |
| Mitsui, 2019 [220] | Subcutaneous xenograft on Balb/c-nu/nu mice | Pancreas | PK-8-GFP and OUMS-24 | Y | exRAGE-Fc (SC; 100 µg/100 µL/6x) | Treated with decoy RAGE | 44d | Decoy RAGE reduced tumor volume but not weight (p<0.01; >0.05) | N/A |
| Muoio, 2023 [57] | Orthotopic xenograft on female athymic nu/nu Swiss mice | Breast | 4T1 | Y | FPS-ZM1 (IP; 1 mg/kg/2x w) | Treated with anti-RAGE therapy | 28d | FPS-ZM1 didn't affect tumor volume (p>0.05) | N/A |
| Nakamura, 2017 [179] | Intradermal xenograft on female athymic nude mice | Skin | G361 | Y | RAGE aptamer (IP; 38.4 pmol/g/d) | Confirmed RAGE expression in tumor tissue, decreased with RAGE-aptamer | 42d | RAGE-aptamer decreased tumor volume (p<0.05) | N/A |
| Nakamura, 2019 [237] | Intradermal xenograft on female athymic nude mice | Skin | G361 | Y | RAGE aptamer (IP; 38.4 pmol/g/d) | Confirmed RAGE expression in tumor tissue, decreased with RAGE-aptamer | 113d | RAGE-aptamer decreased tumor volume (p<0.05) | N/A |
| Nasser, 2015 [59] | Intracardiac injection on nude mice | Breast | SCP2 | Y | naRAGE (20 µg pretreatment) | Treated with neutralizing antibody against RAGE | 60d | N/A | naRAGE decreased metastatic bioluminescence (p<0.05) |
| Nasser, 2015 [59] | Orthotopic xenograft on female C57B/6 RAGE -/- mice | Breast | PyMT | Y | RAGE-/- strain | Confirmed RAGE expression in cell line, crossed with RAGE-/- strain | 5w | RAGE KO decreased tumor volume and weight (p<0.05) | N/A |
| Nasser, 2015 [59] | Orthotopic xenograft on female bitransgenic MMTV-mS100a7a15 mice | Breast | MVT-1 | Y | naRAGE (IP; 20 µg/2d); sRAGE (IP; 2 µg/2d) | Treated with neutralizing antibody against RAGE or sRAGE | 28d | naRAGE and sRAGE decreased tumor volume and weight (p<0.05) | naRAGE and sRAGE decreased number of lung metastasis (p<0.05) |

| | | | | | | | | | |
|---|---|---|---|---|---|---|---|---|---|
| Ojima, 2014 [110] | Intradermal xenograft on female athymic nude mice | Skin | G361 | Y | AGE-aptamer (IP; 0.136 µg) | Confirmed RAGE expression in tumor tissue, decreased with AGE-aptamer | 43d | AGE-aptamer decreased tumor volume (p<0.05) | N/A |
| Pan, 2018 [149] | Subcutaneous xenograft on Balb/c-nu/nu mice | Lungs | PC9, L78 | Y | lncRAGE transfection | Confirmed RAGE expression in cell lines, increased with lncRAGE (no signalling capacity) | 24d | lncRAGE decreased tumor volume (p<0.05) | N/A |
| Pujals, 2023 [180] | Orthotopic xenograft on female NOD-SCID or NMRI mice | Breast | MDA-MB-231 | Y | Azeliragon (IP; 5mg/kg/d) | Confirmed RAGE expression in tumor tissue, decreased with azeliragon | Not specified | Azeliragon didn't affect tumor volume (p>0.05) | Azeliragon didn't affect number of metastases in NOD mice (p=0.126) but decreased in NMRI mice (p=0.033) |
| Pusterla, 2013 [247] | Diethylnitrosamine-induced liver tumors in male Mdr2-/- mice | Liver | N/A | N/A | RAGE-/- strain | Crossed with RAGE-/- strain | 6-12 mo | RAGE KO decreased tumor multiplicity (p<0.001) | N/A |
| Qiao, 2016 [113] | Subcutaneous xenograft on male Balb/c mice | Liver | Bel-7402 | Y | RAGE sh11, sh12 | Confirmed RAGE expression in cell line, decreased with shRAGEs | 8w | RAGE silencing decreased tumor volume (p<0.01) | N/A |
| Rai, 2012 [238] | Intraperitoneal xenograft on female C57/BL6 mice | Ovarian | ID8 | Y | sRAGE (IP; 10 nmol/d), crossed with RAGE-/- strain | Confirmed RAGE expression in cell line, treated with sRAGE, crossed with RAGE-/- strain | 4w | sRAGE and RAGE KO decreased tumor count (p<0.05) | N/A |
| Ray, 2020 [152] | Intraperitoneal xenograft on female BALB/c nu/nu mice | Lungs | A549 | Y | shRAGE; LPA (20 µg/2d) | Confirmed RAGE expression in cell line, decreased with shRAGE | 3w | shRAGE didn't induce tumor colonization (p>0.05) and decreased number of tumor foci (p<0.001) | N/A |
| Ray, 2020 [152] | Intraperitoneal xenograft on female BALB/c nu/nu mice | Breast | MDA-MB-231, MCF-7 | Y | shRAGE; LPA (20 µg/2d) | Confirmed RAGE expression in cell lines, decreased with shRAGE | 3w | shRAGE didn't induce tumor colonization (p>0.05) and decreased number of tumor foci (p<0.001) | N/A |
| Reeb, 2015 [188] | Orthotopic xenograft or tail vein injection on female NOD/SCIDIl2rg-/- mice | Thyroid | THJ-11T, THJ-16T | Y | shS100A8, shS100A9 | Confirmed RAGE expression in tumor tissue and S100A8/RAGE interaction, treated with RAGE ligands | 3-6w | shS100A8 decreased tumor bioluminescence (p<0.05) and shS100A9 didn't affect tumor bioluminescence (p>0.05) | shS100A8 decreased metastatic bioluminescence (p<0.05) and shS100A9 didn't affect metastatic bioluminescence (p=0.305) |
| Ren, 2021 [200] | Subcutaneous xenograft on BALb/c nude mice | Oral | HSC-4 | Y | Evodiamine (Gavage; 3 mg/kg/d); Ov-RAGE transfection | Confirmed RAGE expression in cell line, decreased with EVO, increased with Ov-RAGE | 21d | Evodiamine decreased tumor volume and weight (p<0.001) and Ov-RAGE negated EVO-induced tumor size reduction (p<0.01; 0.001) | N/A |
| Riuzzi, 2007 [171] | Subcutaneous xenograft on female NOD/SCID mice | Musculoskeletal | TE671 | Y | dnRAGE, flRAGE transfection | Confirmed lack of RAGE expression of cell line, transfected with dnRAGE and flRAGE | 6w | dnRAGE and flRAGE decreased tumor volume (p<0.05) | N/A |

| | | | | | | | | | |
|---|---|---|---|---|---|---|---|---|---|
| Ryan, 2019 [214] | Orthotopic xenograft on female immunocompromised nu/nu mice | Breast | Gem240 | N | FPS-ZM1 (IP; 1 mg/kg/d) | Confirmed RAGE expression in cell line, decreased with RAGE inhibitor | 6w | FPS-ZM1 decreased tumor volume (p<0.001) | N/A |
| Shen, 2015 [139] | Subcutaneous xenograft or intraperitoneal injection on nude mice | Colorectum | HCT-116 | Y | SOX9-, SOX9+, SOX9+/S100P- | Confirmed RAGE expression in cell line, decreased with SOX9- and SOX9+/S100P-, increased with SOX9+ | 31d | SOX9- and SOX9+/S100P- decreased tumor volume (p<0.001; 0.005), SOX9+ increased tumor volume (p=0.012) | SOX9- and SOX9+/S100P- decreased metastases count (p=0.037; 0.003), SOX9+ increased metastases count (p=0.013) |
| Shen, 2016 [116] | Subcutaneous xenograft or intraperitoneal injection on nude mice | Colorectum | HCT116, LS174T | Y | S100P KD | Confirmed RAGE expression in cell line, decreased with S100P KD | 28d | S100P KD didn't affect tumor volume (p>0.05) | S100P KD decreased metastases count (p<0.01; 0.001) |
| Siddique, 2013 [210] | Spontaneous prostate tumor in TRAMP mice | Prostate | N/A | N/A | S100A4-/- strain | Confirmed RAGE expression in tumor tissue, decreased with S100A4-/- strain cross | 28w | S100A4-/- decreased tumor volume (p<0.05) | S100A4-/- decreased liver metastasis (p<0.05) |
| Siddique, 2013 [210] | Subcutaneous xenograft on male athymic nu/nu mice | Prostate | LNCaP | Y | S100A4 KO | Treated with RAGE ligand KO | 5w | S100A4 KO decreased tumor volume (p<0.05) | N/A |
| Swami, 2021 [236] | Orthotopic xenograft on C57Bl/6 mice | Pancreas | KPC #5508 | N | IgG 2A11 (IV; 100 µg/5d); gemcitabine (IP; 100 mg/kg/5d) | Confirmed RAGE expression in tumor tissue, treated with anti-RAGE ab | 3w | IgG2A11 increased gemcitabine-induced tumor reduction (p=0.076) | N/A |
| Taguchi, 2000 [119] | Subcutaneous xenograft on female NCR or SCID mice | Brain | C6 | Y | sRAGE (IP; 20 µg/d); anti-RAGE ab (IP; 200 µg/d) | Confirmed RAGE expression in cell line, treated with sRAGE and anti-RAGE antibody | 3w | sRAGE decreased tumor volume (p<0.03) and anti-RAGE ab (p<0.001) | N/A |
| Taguchi, 2000 [119] | Subcutaneous xenograft on male C57BL/6 mice before excision at 1,500 mm3 or intravenous injection | Lungs | LLC | Y | sRAGE (IP; 100 µg/d) | Confirmed RAGE expression in cell line, treated with sRAGE | Not specified (xenograft); 14d (IV) | N/A | sRAGE decreased lung metastasis (p<0.001; 0.00001) and metastatic tumor burden (p<0.001; 0.00001) |
| Takeuchi, 2013 [150] | Subcutaneous xenograft or tail vein injection on female BALB/c-nu/nu mice | Musculoskeletal | HT1080 | Y | flRAGE, dnRAGE transfection | Confirmed RAGE transfection in cell lines | 4w | flRAGE increased tumor volume (p<0.05) and dnRAGE didn't affect tumor volume (p>0.05) | flRAGE increased number of lung metastases (p<0.01) and dnRAGE decreased number of lung metastases (p<0.01) |
| Wang, 2021 [122] | Subcutaneous xenograft on male C57BL/6 mice | Lungs | LLC | Y | Dipyridamole (PO; 10 mg/kg/d) | Confirmed RAGE expression in cell line, decreased with dipyridamole | 3w | Dipyridamole decreased tumor volume (p<0.05) | N/A |
| Wang, 2025 [235] | Subcutaneous xenograft on female NOD-SCID mice | Lungs | NCI-H322 | Y | oeRAGE transfection | Transfected cell line with oeRAGE | 30d | oeRAGE didn't increase tumor volume (p>0.05) | N/A |

| | | | | | | | | | |
|---|---|---|---|---|---|---|---|---|---|
| Wu, 2015 [192] | Subcutaneous xenograft on female nude mice | Liver | HepG2 | Y | anti-RAGE ab (intratumor; 160 µg/mL); GST-S100A9 (intratumor; 20 µg/mL) | Confirmed RAGE expression in cell line, increased with S100A9, treated with anti-RAGE ab | 20d | Anti-RAGE antibody didn't affect tumor volume (p>0.05) | N/A |
| Wuren, 2021 [250] | Tail vein injection on C57BL/6 mice | Skin | B16-F10 | Y | RAGE-/- strain | Confirmed RAGE expression in cell line, crossed with RAGE-/- strain | 21d | N/A | RAGE KO decreased metastatic tumor burden (p<0.05) |
| Wuren, 2021 [250] | Tail vein injection on C57BL/6 mice | Lungs | LLC | Y | RAGE-/- strain | Confirmed RAGE expression in cell line, crossed with RAGE-/- strain | 21d | N/A | RAGE kO decreased metastatic tumor burden (p<0.01) |
| Yang, 2024 [205] | Subcutaneous xenograft on NSG BALB/c or intravenous injection on BALB/c nu/nu mice | Kidneys | A498 | Y | Corylin (Gavage; 60 mg/kg/d) | Confirmed RAGE expression in cell line, decreased with corylin | 2m | Corylin decreased tumor bioluminescence (p<0.01) | Corylin decreased lung metastatic bioluminescence (p<0.01) |
| You, 2024 [226] | Subcutaneous xenograft on male BALB/c nu/nu mice | Prostate | PC-3 | Y | Astragaloside (gavage; 40 mg/kg); Scorpion venom polypeptide (PESV) (IP; 1.2 mg/kg); Fecal microbiota transplantation (FMT)-astragaloside IV-PESV (gavage; 0.1 mL/d); oeRAGE | Confirmed RAGE expression in cell line; decreased with PESV, astragaloside IV, astra-PESV, FMT-astra-PESV; increased with oeRAGE | 27d | All drug combinations and fecal microbiota transplantation decreased tumor weight and volume (p<0.05); oeRAGE negated fecal microbiota transplantation-induced tumor size reduction (p<0.05) | N/A |
| Yu, 2017 [213] | Subcutaneous xenograft on female BALB/c nu/nu mice | Lungs | H1975 | Y | siRAGE | Confirmed RAGE expression in cell line, decreased with siRAGE | 21d | siRAGE decreased tumor volume and weight (p<0.05) | N/A |
| Zhang, 2012 [252] | Orthotopic implantation of human tumor tissue on female SCID mice | Stomach | Human gastric carcinoma | N | Ethyl pyruvate (IP; 80 mg/kg/d) | Confirmed RAGE expression in tumor tissue, decreased with ethyl pyruvate | 3w | Ethyl pyruvate decreased tumor weight and volume (p<0.01) | Ethyl pyruvate decreased liver metastases number (p<0.01) |
| Zhang, 2023 [243] | Stereotactic intracranial implantation on female C57BL/6J mice | Brain | GL261, K-Luc | Y | RAGE-KD, oeRAGE | Confirmed RAGE expression in cell line, decreased with RAGE KD, increased with oeRAGE | 2w | RAGE KD decreased tumor volume (p<0.05) and oeRAGE increased tumor volume (p<0.05) | RAGE KD decreased invasion scores (p<0.01) |
| Zhang, 2025 [233] | Orthotopic xenograft on male C57BL/6J mice | Liver | Hepa1-6 | Y | shS100A9, RAGE KO (TLR4KO, Lyz2Cre, ETV5), S100A9 | Confirmed RAGE expression in mice, decreased with RAGE KO, treated with RAGE ligand silencing or RAGE ligand | 10w, 9w | shS100A9 and RAGE KO decreased tumor bioluminescence (p=0.001) | shS100A9 and RAGE KO decreased lung metastasese (p=0.026; 0.002) and S100A9 increased lung metastases (p=0.008) |
| Zheng, 2016 [231] | Subcutaneous xenograft on female BALB/c nude mice | Endometrium | HEC-1A | Y | siRAGE | Confirmed RAGE expression in tumor tissue, decreased with siRAGE | 20d | siRAGE decreased tumor weight (p<0.05) but didn't affect tumor volume (p>0.05) | N/A |

| | | | | | | | | | |
|---|---|---|---|---|---|---|---|---|---|
| Zhou, 2025 [240] | Subcutaneous xenograft on male C57BL/6J mice | Colorectum | APCmin/+ derived organoids | N | oeS100A11, FPS-ZM1 (IP; 5 mg/kg/3x d) | Treated with RAGE ligand overexpression and RAGE inhibitor | 2w | S100A11 increased tumor growth (p<0.001) and FPS-ZM1 negated S100A11-induced tumor growth (p=0.003) | N/A |
| Zhou, 2025 [240] | Subcutaneous xenograft on male NSG mice | Colorectum | CT26 | Y | FPS-ZM1 (IP; 5 mg/kg/3x d), Azeliragon (Gavage; 4 mg/kg/d) | Treated with RAGE inhibitors | 2w; 19d | FPS-ZM1 decreased tumor growth (p<0.001; =0.037) and Azeliragon decreased tumor growth (p=0.012; 0.029) | N/A |

Abbreviations: advanced glycation end-products (AGE), receptor for advanced glycation end-products (RAGE), full length RAGE (flRAGE), RAGE-overexpression (oeRAGE; RAGE-OV), soluble RAGE (sRAGE), short hairpin RAGE (shRAGE), dominant negative RAGE (dnRAGE), RAGE-knockout (RAGE-KO), RAGE-knockdown (RAGE-KD), neutralizing antibody RAGE (naRAGE), long non-coding (lncRAGE), high mobility group box 1 (HMGB1), intraperitoneal (IP), subcutaneous (SC), per os (PO), severe combined immunodeficiency (SCID), 4-nitroquinoline 1-oxide (4NQO), pterostilbene (PT), chloroquine (CQ), ursolic acid (UA), lysophosphatidic acid (LPA), acetylsalicylic acid (ASA), trichostatin A (TSA), azoxymethane/dextran sodium sulfate (AOM/DSS)
