## Supplemental Table 6 for "The Role of the Receptor for Advanced Glycation End-Products in Cancer: Evidence from a Systematic Review and Meta-Analysis"

| | Selection Bias | | | Performance Bias | | Detection Bias | | | | |
|---|---|---|---|---|---|---|---|---|---|---|
| Authors | Sequence Generation[1] | Baseline[2] | Allocation Concealment[3] | Random Housing[4] | Blinding[5] | Random Outcome Assessment[6] | Blinding[7] | Attrition Bias[8] | Reporting Bias[9] | Other Bias[10] |
| Abe, 2004 [184] | Unclear | Low | Unclear | Low | Unclear | Unclear | Unclear | Low | Low | Low |
| Alka, 2024 [207] | Unclear | Low | Low | Low | Unclear | Unclear | Unclear | Low | Low | Low |
| Arumugam, 2005 [196] | Unclear | Low | Unclear | Low | Unclear | Unclear | Unclear | Low | Low | Low |
| Arumugam, 2006 [81] | Unclear | Low | Unclear | Low | Unclear | Unclear | Unclear | Low | Low | Low |
| Arumugam, 2012 [197] | Unclear | Low | Unclear | Low | Unclear | Unclear | Unclear | Low | Low | Low |
| Arumugam, 2013 [230] | Unclear | Low | Unclear | Low | Unclear | Unclear | Unclear | Low | Low | Low |
| Bartling, 2005 [83] | Unclear | Low | Unclear | Low | Unclear | Unclear | Unclear | Low | Low | Low |
| Chen, 2014 [244] | Unclear | Low | Unclear | Low | Unclear | Unclear | Unclear | Low | Low | Low |
| Chen, 2020 [170] | Low | Low | Unclear | Low | Unclear | Unclear | Low | Low | Low | Low |
| Chen, 2021 [227] | Unclear | Low | Unclear | Low | Unclear | Unclear | Unclear | Low | Low | Low |
| Chen, 2024 [186] | Unclear | Low | Unclear | Low | Unclear | Unclear | Unclear | Low | Low | Low |
| Chen, 2024 [251] | Unclear | Low | Unclear | Low | Unclear | Unclear | Unclear | Low | Low | Low |
| Cheng, 2014 [232] | Low | Low | Unclear | Low | Unclear | Unclear | Unclear | Low | Low | Low |
| Chiappalupi, 2014 [229] | Unclear | Low | Unclear | Low | Unclear | Unclear | Unclear | Low | Low | High |
| Cho, 2024 [228] | Unclear | Low | Unclear | Low | Unclear | Unclear | Unclear | Low | Low | Low |
| Dai, 2023 [32] | Low | Low | Unclear | Low | Unclear | Unclear | Unclear | Low | Low | Low |
| DiNorcia, 2010 [245] | Low | Low | Unclear | Low | Unclear | Unclear | Low | Low | Low | Low |
| Elangovan, 2012 [216] | Unclear | Low | Unclear | Low | Unclear | Unclear | Unclear | Low | Low | Low |
| Fan, 2024 [239] | Low | Low | Unclear | Low | Unclear | Unclear | Unclear | Low | Low | Low |
| Heijmans, 2013 [246] | Low | Low | Unclear | Unclear | Unclear | Unclear | Unclear | Unclear | Unclear | Low |
| Hernandez, 2013 [253] | Unclear | Low | Unclear | Low | Unclear | Unclear | Unclear | Low | Low | Low |
| Hiramoto, 2021 [234] | Low | Low | Unclear | Low | Unclear | Unclear | Unclear | Low | Low | Low |
| Hou, 2025 [225] | Low | Low | Unclear | Low | Unclear | Unclear | Unclear | Low | Low | Low |
| Huttunen, 2002 [183] | Unclear | Low | Unclear | Low | Unclear | Unclear | Unclear | Low | Low | Low |
| Ichikawa, 2011 [189] | Unclear | Low | Unclear | Low | Unclear | Unclear | Unclear | Low | Low | Low |
| Jiang, 2025 [160] | Low | Low | Unclear | Low | Unclear | Unclear | Unclear | Low | Low | Low |
| Jing, 2015 [164] | Unclear | Low | Unclear | Low | Unclear | Unclear | Unclear | Low | Low | Low |
| Kang, 2010 [208] | Unclear | Low | Unclear | Low | Unclear | Unclear | Unclear | Low | Low | Low |
| Kang, 2014 [203] | Unclear | Low | Unclear | Low | Unclear | Unclear | Unclear | Low | Low | Low |
| Khoo, 2023 [45] | Low | Low | Unclear | Low | Low | Unclear | Low | Low | Low | Low |
| Kim, 2025 [96] | Unclear | Low | Unclear | Low | Unclear | Unclear | Unclear | Low | Low | Low |
| Krisanits, 2022 [242] | Unclear | Low | Unclear | Low | Unclear | Unclear | Unclear | Low | Low | Low |
| Kwak, 2017 [215] | Low | Low | Low | Low | Low | Unclear | Low | Low | Low | Low |
| Lee, 2018 [217] | Unclear | Low | Unclear | Low | Unclear | Unclear | Unclear | Low | Low | Low |
| Li, 2018 [172] | Unclear | Low | Unclear | Low | Unclear | Unclear | Unclear | Low | Low | Low |
| Li, 2021 [254] | Low | Low | Unclear | Low | Unclear | Unclear | Unclear | Low | Low | Low |
| Li, 2024 [241] | Low | Low | Unclear | Low | Unclear | Unclear | Unclear | Low | Low | Low |
| Mark, 2013 [249] | Unclear | Low | Unclear | Low | Unclear | Unclear | Unclear | Low | Low | Low |
| Meghnani, 2014 [248] | Low | Low | Unclear | Low | Unclear | Unclear | Unclear | Low | Low | Low |

| | | | | | | | | | | |
|---|---|---|---|---|---|---|---|---|---|---|
| Mitsui, 2019 [220] | Unclear | Low | Unclear | Low | Unclear | Unclear | Unclear | Low | Low | Low |
| Muoio, 2023 [57] | Low | Low | Unclear | Low | Unclear | Unclear | Unclear | Low | Low | Low |
| Nakamara, 2017 [179] | Unclear | Low | Unclear | Low | Unclear | Unclear | Unclear | Low | Low | Low |
| Nakamura, 2019 [237] | Unclear | Low | Unclear | Low | Unclear | Unclear | Unclear | Low | Low | Low |
| Nasser, 2015 [59] | Unclear | Low | Unclear | Low | Unclear | Unclear | Unclear | Low | Low | Low |
| Ojima, 2014 [110] | Unclear | Low | Unclear | Low | Unclear | Unclear | Unclear | Low | Low | Low |
| Pan, 2018 [149] | Unclear | Low | Unclear | Low | Unclear | Unclear | Unclear | Low | Low | Low |
| Pujals, 2023 [180] | Low | Low | Unclear | Low | Unclear | Unclear | Unclear | Low | Low | Low |
| Pusterla, 2013 [247] | Low | Low | Unclear | Low | Unclear | Unclear | Unclear | Low | Low | Low |
| Qiao, 2016 [113] | Unclear | Low | Unclear | Low | Unclear | Unclear | Unclear | Low | Low | Low |
| Rai, 2012 [238] | Unclear | Low | Unclear | Low | Unclear | Unclear | Unclear | Low | Low | Low |
| Ray, 2020 [152] | Unclear | Low | Unclear | Low | Unclear | Unclear | Unclear | Low | Low | Low |
| Reeb, 2015 [188] | Unclear | Low | Unclear | Low | Unclear | Unclear | Unclear | Low | Low | Low |
| Ren, 2021 [200] | Unclear | Low | Unclear | Low | Unclear | Unclear | Unclear | Low | Low | Low |
| Riuzzi, 2007 [171] | Unclear | Low | Unclear | Low | Unclear | Unclear | Unclear | Low | Low | Low |
| Ryan, 2019 [214] | Low | Low | Unclear | Low | Unclear | Unclear | Unclear | Low | Low | Low |
| Shen, 2015 [139] | Unclear | Low | Unclear | Low | Unclear | Unclear | Unclear | Low | Low | Low |
| Shen, 2016 [116] | Unclear | Low | Unclear | Low | Unclear | Unclear | Unclear | Low | Low | Low |
| Siddique, 2013 [210] | Unclear | Low | Unclear | Low | Unclear | Unclear | Unclear | Low | Low | Low |
| Swami, 2021 [236] | Unclear | Low | Unclear | Low | Unclear | Unclear | Unclear | Low | Low | Low |
| Taguchi, 2000 [119] | Unclear | Low | Unclear | Low | Unclear | Unclear | Unclear | Low | Low | Low |
| Takeuchi, 2013 [150] | Unclear | Low | Unclear | Low | Unclear | Unclear | Unclear | Low | Low | Low |
| Wang, 2021 [122] | Unclear | Low | Unclear | Low | Unclear | Unclear | Unclear | Low | Low | Low |
| Wang, 2025 [235] | Unclear | Low | Unclear | Low | Unclear | Unclear | Unclear | Low | Low | Low |
| Wu, 2015 [192] | Low | Low | Unclear | Low | Unclear | Unclear | Unclear | Low | Low | Low |
| Wuren, 2021 [250] | Unclear | Low | Unclear | Low | Unclear | Unclear | Unclear | Low | Low | Low |
| Yang, 2024 [205] | Low | Low | Unclear | Low | Unclear | Unclear | Unclear | Low | Low | Low |
| You, 2024 [226] | Unclear | Low | Unclear | Low | Unclear | Unclear | Unclear | Low | Low | Low |
| Yu, 2017 [213] | Unclear | Low | Unclear | Low | Unclear | Unclear | Unclear | Low | Low | Low |
| Zhang, 2012 [252] | Low | Low | Unclear | Low | Unclear | Unclear | Unclear | Low | Low | Low |
| Zhang, 2023 [243] | Unclear | Low | Unclear | Low | Unclear | Unclear | Unclear | Low | Low | Low |
| Zhang, 2025 [233] | Unclear | Low | Unclear | Low | Unclear | Unclear | Unclear | Low | Low | Low |
| Zheng, 2016 [231] | Unclear | Low | Unclear | Low | Unclear | Unclear | Lowow | Low | Low | Low |
| Zhou, 2025 [240] | Low | Low | Unclear | Low | Unclear | Unclear | Unclear | Low | Low | Low |

Studies are given a risk of bias of either “high” (disagreement with parameters), “low” (agreement with parameters), or “unclear” (unclear is parameters were met/unmet) based on the following parameters: 1random allocation of animals; 2similarity of baseline characteristics; 3allocation blinding; 4random housing distribution within the room; 5investigator blinding; 6random animal selection for outcome assessment; 7outcome assessor blinding; 8incomplete outcome data addressed; 9 free from selective outcome reporting; 10 free from any other potential sources of bias (e.g., contamination, funding sources, unit of analysis errors). No summary score is given to avoid assigning weights to each category.
