## Supplemental Table 7 for "The Role of the Receptor for Advanced Glycation End-Products in Cancer: Evidence from a Systematic Review and Meta-Analysis"

| | Cancer | Type | p-value | n | Cancer | Type | p-value | n |
|---|---|---|---|---|---|---|---|---|
| RAGE-Positive | Breast | Growth Potential | 6.45E-05 | 464 | Brain | Growth Potential | 0.12† | 314 |
| | | Metastatic Potential | 7.01E-07 | 862 | | Metastatic Potential | 1.16E-17 | 140 |
| | | Apoptosis | 0.34† | 18 | | NF-kB | 9.95E-03 | 30 |
| | | NF-kB | 3.69E-03 | 12 | Kidney | Growth Potential | 0.36† | 102 |
| | Colorectal | Growth Potential | 3.49E-05 | 408 | | Metastatic Potential | 8.05E-11 | 132 |
| | | Metastatic Potential | 7.67E-39 | 512 | | Apoptosis | 8.39E-04 | 72 |
| | | Apoptosis | 1.84E-06 | 96 | Cervical | Growth Potential | 1.15E-06 | 162 |
| | | NF-kB | 6.80E-04 | 40 | | Metastatic Potential | 4.69E-10 | 54 |
| | Pancreatic | Growth Potential | 5.46E-05 | 510 | | Apoptosis | 1.10E-03 | 54 |
| | | Metastatic Potential | 0.11† | 102 | Thyroid | Growth Potential | 7.01E-05 | 40 |
| | | Apoptosis | 2.73E-03 | 60 | | Metastatic Potential | 1.26E-03 | 24 |
| | | NF-kB | 1.18E-12 | 212 | | Apoptosis | 0.01 | 12 |
| | Musculoskeletal | Growth Potential | 0.08* | 76 | Gall Bladder | Growth Potential | 0.04 | 12 |
| | | Metastatic Potential | 1.25E-05 | 258 | | Metastatic Potential | 7.80E-04 | 12 |
| | | Apoptosis | 0.02* | 12 | | Apoptosis | 5.02E-04 | 12 |
| | | NF-kB | 3.89E-03 | 36 | Ovarian | Growth Potential | 0.70† | 36 |
| | Liver | Growth Potential | 1.36E-05 | 102 | | Apoptosis | 1.00E-03* | 6 |
| | | Metastatic Potential | 0.05 | 36 | Blood | Growth Potential | 7.01E-04 | 12 |
| | | NF-kB | 0.01 | 6 | | Apoptosis | 5.13E-04 | 12 |
| | Prostate | Growth Potential | 8.16E-03 | 72 | Endometrial | Growth Potential | 7.36E-06 | 72 |
| | | Metastatic Potential | 1.00E-04 | 6 | | Metastatic Potential | 3.71E-10 | 72 |
| | | NF-kB | 9.19E-05 | 12 | Mesothelioma | Growth Potential | 2.62E-03 | 64 |
| | Skin | Growth Potential | 1.60E-04 | 116 | | Metastatic Potential | 4.12E-03 | 30 |
| | | Metastatic Potential | 2.75E-12 | 396 | Head and Neck | Metastatic Potential | 2.71E-11 | 72 |
| | | NF-kB | 0.071*† | 32 | Bile Duct | Metastatic Potential | 0.05 | 6 |
| | Combined | Growth Potential | 2.95E-05 | 2562 | | | | |
| | | Metastatic Potential | 4.24E-15 | 2714 | | | | |
| | | Apoptosis | 4.26E-13 | 354 | | | | |
| | | NF-kB | 1.53E-04 | 380 | | | | |
| RAGE-Negative | Lung | Growth Potential | 0.86† | 292 | Oral | Growth Potential | 0.20† | 60 |
| | | Metastatic Potential | 0.78† | 178 | | Metastatic Potential | 9.23E-04 | 66 |
| | | Apoptosis | 2.34E-03 | 42 | | Apoptosis | 1.00E-03 | 6 |
| | | NF-kB | 0.10† | 18 | Gastric | Growth Potential | 0.12† | 100 |
| | Nasoesophageal | Growth Potential | 5.13E-04 | 84 | | Apoptosis | 4.85E-03 | 12 |
| | | Metastatic Potential | 1.64E-17 | 108 | | | | |
| | | Apoptosis | 6.52E-04 | 24 | | | | |
| | | NF-kB | 6.68E-04 | 24 | | | | |
| | Combined | Growth Potential | 2.12E-03 | 536 | | | | |
| | | Metastatic Potential | 0.11† | 352 | | | | |
| | | Apoptosis | 3.53E-05 | 84 | | | | |
| | | NF-kB | 4.80E-04 | 42 | | | | |
| Full Meta-Analysis | | Growth Potential | 4.80E-06 | 3098 | | | | |
| | | Metastatic Potential | 7.62E-12 | 3066 | | | | |
| | | Apoptosis | 1.84E-14 | 438 | | | | |
| | | NF-kB | 1.21E-05 | 422 | | | | |
