## Supplemental Table 8 for "The Role of the Receptor for Advanced Glycation End-Products in Cancer: Evidence from a Systematic Review and Meta-Analysis"

| | Cancer | Type | p-value | n | Cancer | Type | p-value | n |
|---|---|---|---|---|---|---|---|---|
| RAGE-Positive | Breast | Tumor Growth | 5.9455E-05 | 382 | Skin | Tumor Growth | 9.04E-03 | 124 |
| | | Metastasis | 1.39E-04 | 154 | | Metastasis | 5.82E-08 | 52 |
| | Colorectal | Tumor Growth | 4.26E-06 | 258 | Brain | Tumor Growth | 2.20E-03 | 104 |
| | | Metastasis | 1.19E-05 | 148 | | Metastasis | 8.05E-04 | 24 |
| | Pancreatic | Tumor Growth | 8.48E-06 | 266 | Kidney | Tumor Growth | 0.01 | 30 |
| | | Metastasis | 1.93E-03 | 90 | | Metastasis | 0.01 | 30 |
| | Musculosk-eletal | Tumor Growth | 0.57† | 72 | Thyroid | Tumor Growth | 0.10† | 32 |
| | | Metastasis | 5.76E-04* | 129 | | Metastasis | 0.043 | 16 |
| | Liver | Tumor Growth | 1.86E-04 | 144 | Ovarian | Tumor Growth | 0.01 | 20 |
| | | Metastasis | 2.22E-04 | 50 | Intestinal | Tumor Growth | 0.01 | 16 |
| | Prostate | Tumor Growth | 1.23E-04 | 184 | Endome-trial | Tumor Growth | 0.05 | 48 |
| | | Metastasis | 0.05 | 6 | | | | |
| | Combined | Tumor Growth | 9.59E-09 | 1664 | | | | |
| | | Metastasis | 1.09E-11 | 699 | | | | |
| RAGE-Negative | Lung | Tumor Growth | 3.19E-04 | 144 | Gastric | Tumor Growth | 9.92E-03 | 24 |
| | | Metastasis | 1.59E-07 | 44 | | Metastasis | 0.01 | 12 |
| | Nasoesop-hageal | Tumor Growth | 0.90† | 70 | Oral | Tumor Growth | 1.19E-04 | 66 |
| | | Metastasis | 0.003 | 6 | | | | |
| | Combined | Tumor Growth | 1.60E-07 | 320 | | | | |
| | | Metastasis | 2.42E-08 | 62 | | | | |
| Full Meta-Analysis | | Tumor Growth | 9.59E-09 | 1984 | | | | |
| | | Metastasis | 1.46E-13 | 761 | | | | |
