## Supplemental Table 9 for "The Role of the Receptor for Advanced Glycation End-Products in Cancer: Evidence from a Systematic Review and Meta-Analysis"

| | Cancer | Type | p-value | n | Cancer | Type | p-value | n |
|---|---|---|---|---|---|---|---|---|
| RAGE-Positive | Breast | Control vs Cancer | 3.33E-04 | 149 | Musculoskeletal | Control vs Cancer | 1.00E-03 | 54 |
| | | Staging | 9.52E-04 | 1499 | Kidney | Control vs Cancer | 8.44E-03 | 38 |
| | Prostate | Control vs Cancer | 7.58E-03 | 172 | Ovarian | Control vs Cancer | 1.00E-03 | 444 |
| | | Staging | 0.5† | 78 | Bladder | Control vs Cancer | 0.05 | 34 |
| | Liver | Control vs Cancer | 4.90E-03 | 72 | | | | |
| | Combined | Control vs Cancer | 1.60E-05 | 963 | | | | |
| | | Staging | 9.96E-04 | 1577 | | | | |
| RAGE-Negative | Lung | Control vs Cancer | 0.70† | 2309 | Oral | Control vs Cancer | 0.05 | 33 |
| | Gastric | Control vs Cancer | 1.00E-03* | 60 | | Staging | 0.5† | 20 |
| | Nasoesophageal | Control vs Cancer | 1.08E-03* | 434 | | | | |
| | Combined | Control vs Cancer | 0.046* | 2836 | | | | |
| | | Staging | 0.5† | 20 | | | | |
| Full Meta-Analysis | | Control vs Cancer | 0.62† | 3799 | | | | |
| | | Staging | 1.00E-03 | 1597 | | | | |
