## Supplemental Figure Legends for "The Role of the Receptor for Advanced Glycation End-Products in Cancer: Evidence from a Systematic Review and Meta-Analysis"

- **Supplemental Figure S1**. Histogram plot of included publications over time, stratified by study type.
- **Supplemental Figure S2**. Funnel plots of RAGE expression in (**A**) cancer, (**B**) pro-RAGE cancers, and (**C**) nulli-RAGE cancers. Used for visual assessment of publication bias or study heterogeneity. Each dot represents a single study, the gray lines indicate 95% CI, and the red vertical line is the overall effect. Asymmetry in the funnel plot, indicated by greater numbers on one side of the overall effect, suggests publication bias or increased study heterogeneity.
- **Supplemental Figure S3**. Leave-one-out plots of RAGE expression in (**A**) cancer, (**B**) pro-RAGE cancers, and (**C**) nulli-RAGE cancers. Used to determine if one study is over-weighing the effect. Each line omits a single study, with the grey dot representing the subsequent OR and the grey line representing the subsequent 95% CI. The vertical dashed red line indicates the effect from the correlated forest plot.
- **Supplemental Figure S4**. Funnel plots of RAGE expression in (**A**) cancer grading, (**B**) pro-RAGE cancer grading, and (**C**) nulli-RAGE cancer grading. Used for visual assessment of publication bias or study heterogeneity. Each dot represents a single study, the gray lines indicate 95% CI, and the red vertical line is the overall effect. Asymmetry in the funnel plot, indicated by greater numbers on one side of the overall effect, suggests publication bias or increased study heterogeneity.
- **Supplemental Figure S5**. Leave-one-out plots of RAGE expression in (**A**) cancer grading, (**B**) pro-RAGE cancer grading, and (**C**) nulli-RAGE cancer grading. Used to determine if one study is over-weighing the effect. Each line omits a single study, with the grey dot representing the subsequent OR and the grey line representing the subsequent 95% CI. The vertical dashed red line indicates the effect from the correlated forest plot.
- **Supplemental Figure S6**. Funnel plots of RAGE expression in (**A**) cancer invasion of regional lymph nodes, (**B**) pro-RAGE cancer invasion, and (**C**) nulli-RAGE cancer invasion. Used for visual assessment of publication bias or study heterogeneity. Each dot represents a single study, the gray lines indicate 95% CI, and the red vertical line is the overall effect. Asymmetry in the funnel plot, indicated by greater numbers on one side of the overall effect, suggests publication bias or increased study heterogeneity.
- **Supplemental Figure S7**. Leave-one-out plots of RAGE expression in (**A**) cancer invasion of regional lymph nodes, (**B**) pro-RAGE cancer invasion, and (**C**) nulli-RAGE cancer invasion. Used to determine if one study is over-weighing the effect. Each line omits a single study, with the grey dot representing the subsequent OR and the grey line representing the subsequent 95% CI. The vertical dashed red line indicates the effect from the correlated forest plot.
- **Supplemental Figure S8**. Forest plot of RAGE expression in patient survival. These associations were indicated as odds ratio (OR) estimates with a corresponding 95% confidence interval (CI).  Gray line indicates OR of 1 or null association. Results are indicated by red dots, indicating OR estimate, and horizontal red lines, indicating 95% CI. Vertical red line and green diamond indicates the overall OR estimate. Stratified results for the nulli-RAGE and pro-RAGE groups are also indicated.

- **Supplemental Figure S9**. Funnel plots of RAGE expression in (**A**) cancer patient survival, (**B**) pro-RAGE cancer patient survival, and (**C**) nulli-RAGE cancer patient survival. Used for visual assessment of publication bias or study heterogeneity. Each dot represents a single study, the gray lines indicate 95% CI, and the red vertical line is the overall effect. Asymmetry in the funnel plot, indicated by greater numbers on one side of the overall effect, suggests publication bias or increased study heterogeneity.

- **Supplemental Figure S10**. Leave-one-out plots of RAGE expression in (**A**) cancer patient survival, (**B**) pro-RAGE cancer patient survival, and (**C**) nulli-RAGE cancer patient survival. Used to determine if one study is over-weighing the effect. Each line omits a single study, with the grey dot representing the subsequent OR and the grey line representing the subsequent 95% CI. The vertical dashed red line indicates the effect from the correlated forest plot.

- **Supplementary Figure S11.** Albatross plot of clinical studies measuring RAGE expression with (**A**) cancer, measured by molecular assays; and (**B**) cancer grading, measured by molecular assays. Each point represents data and its size is correlated to number of overlapping points, with the effect estimate (represented as a p-value), plotted against the total given sample size (n) included within each study. The directionality of the p-values correlate to negative associations with RAGE for the left and positive associations with RAGE for the right. Contour lines are standardized mean differences (SMD). Stratification for nulli-RAGE and pro-RAGE cancers was done by color, as described in the legends of each panel. Non-exact p-values reported were plotted as stated in the manuscript (e.g., if $p < 0.05$, plotted $p = 0.05$; if $p > 0.05$, plotted $p = 0.5$) as an estimate.
