## Supplementary figures and images for "The Role of the Receptor for Advanced Glycation End-Products in Cancer: Evidence from a Systematic Review and Meta-Analysis"

### Supplemental Figure S1

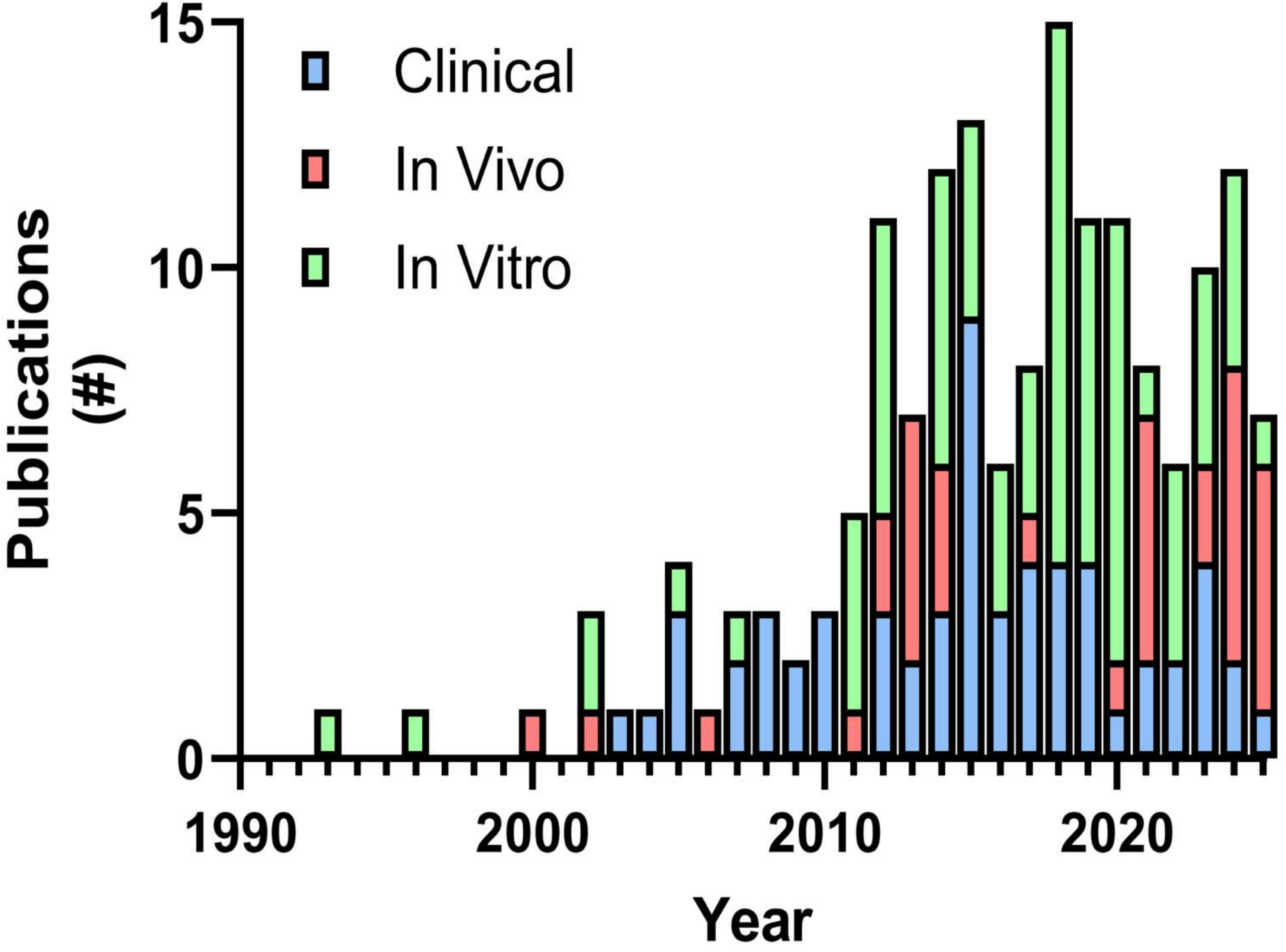

### Supplemental Figure S2

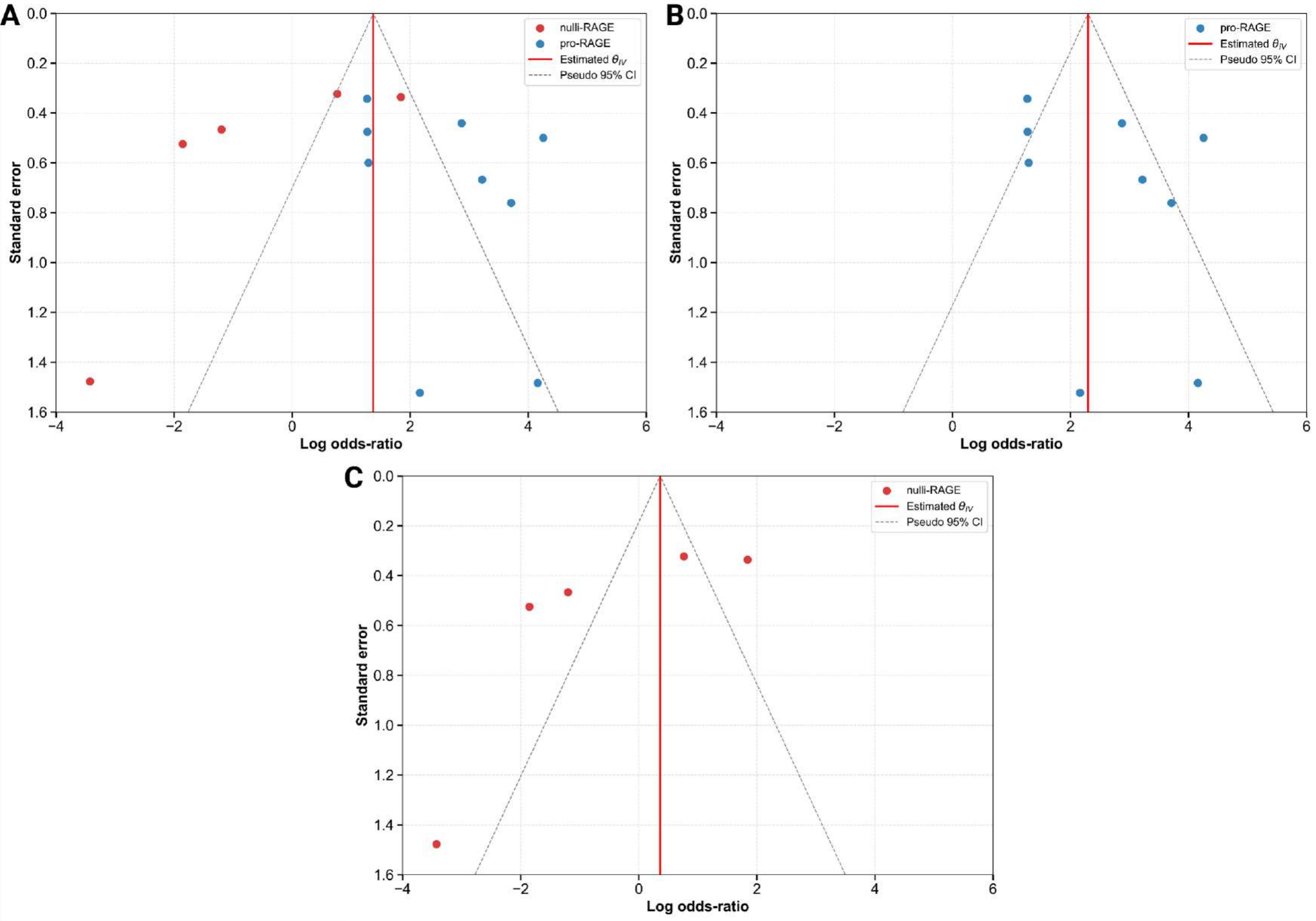

### Supplemental Figure S3

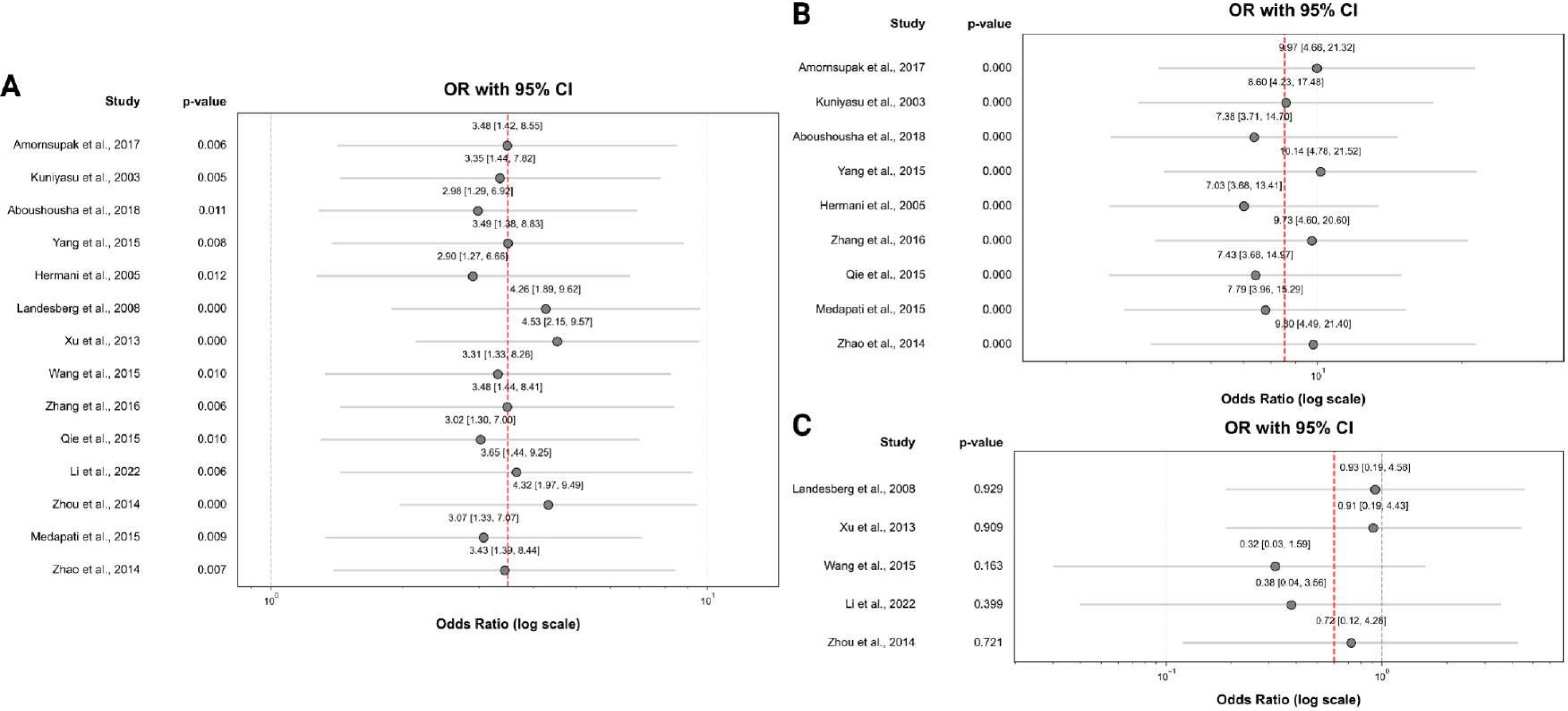

### Supplemental Figure S4

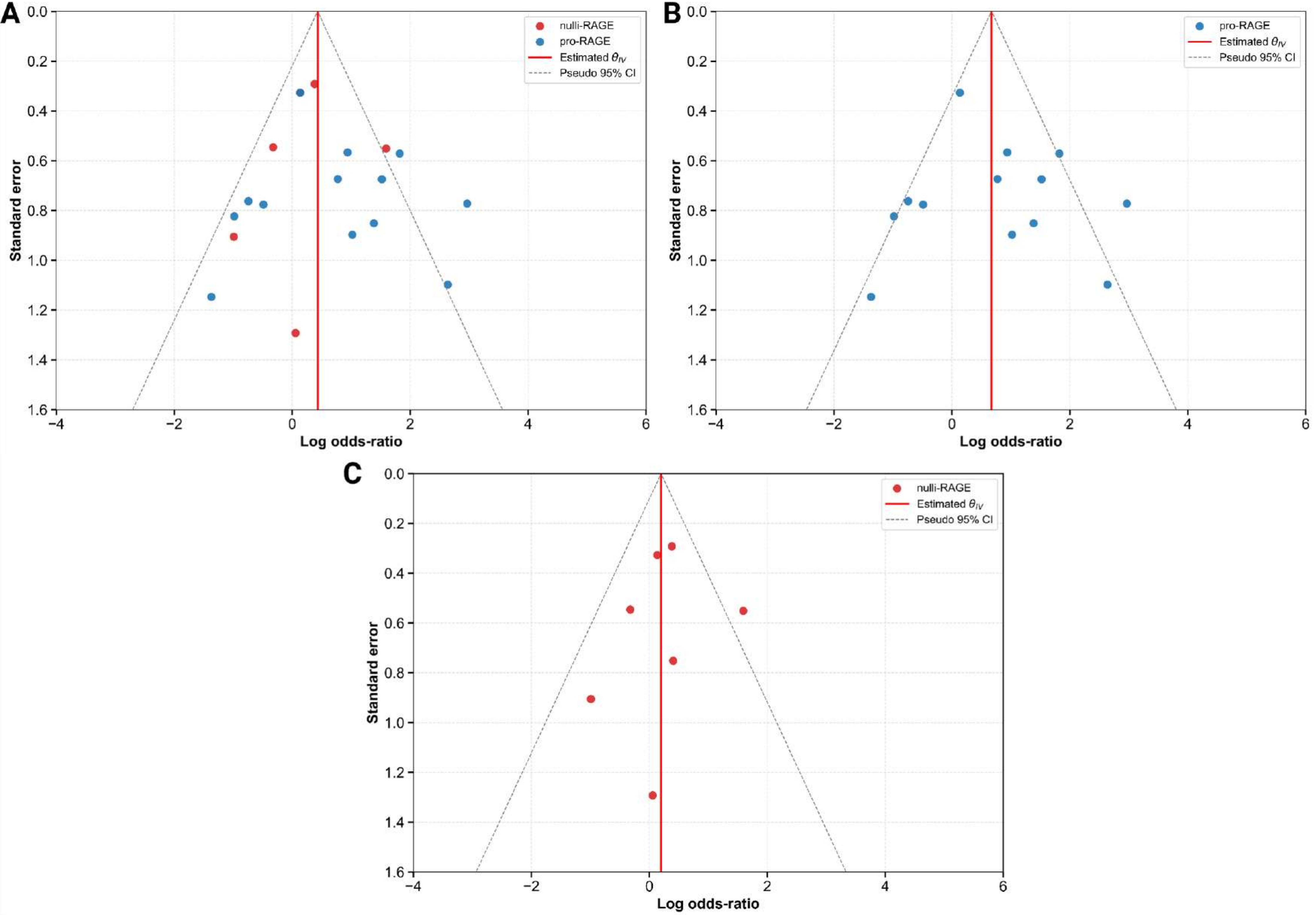

### Supplemental Figure S5

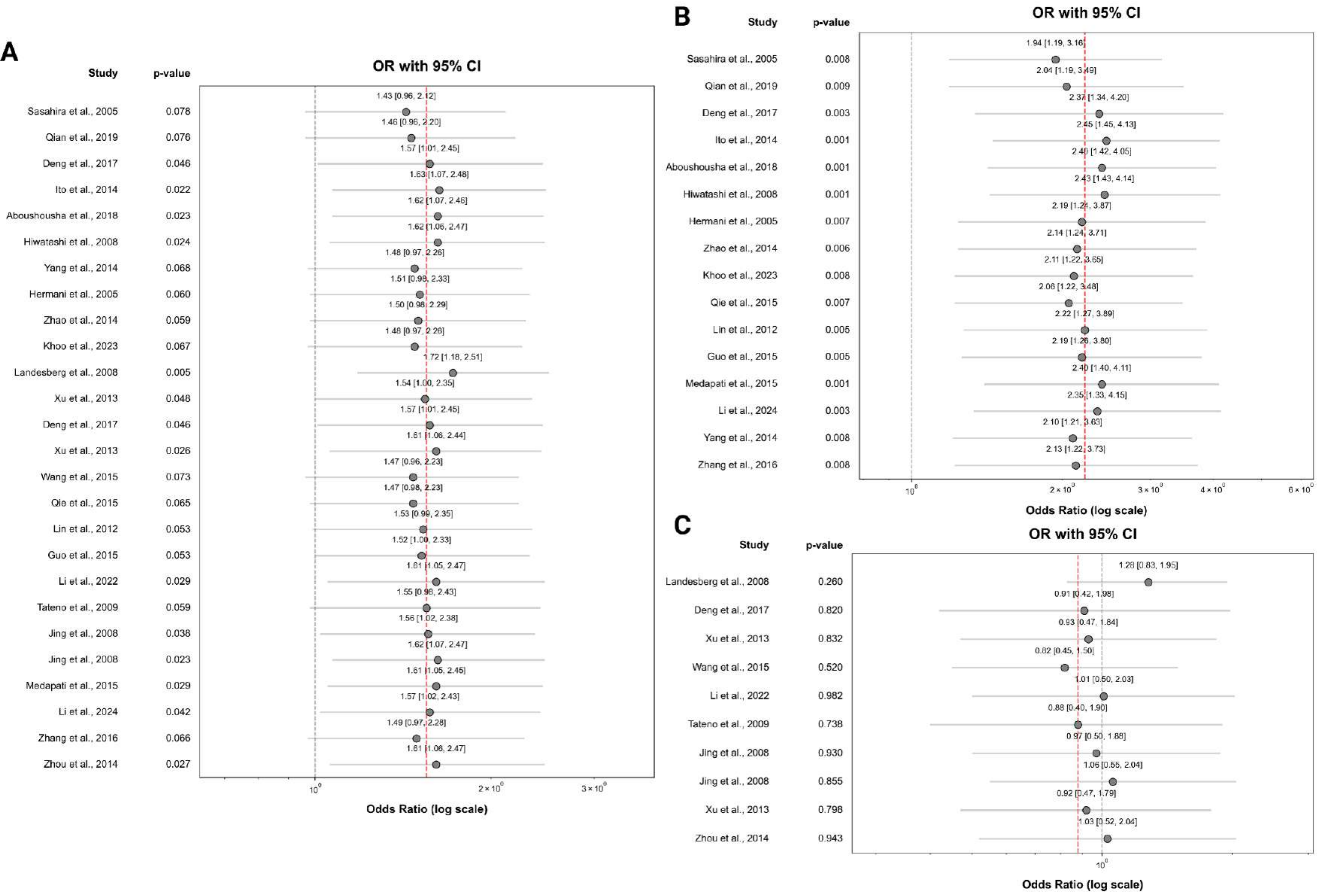

### Supplemental Figure S6

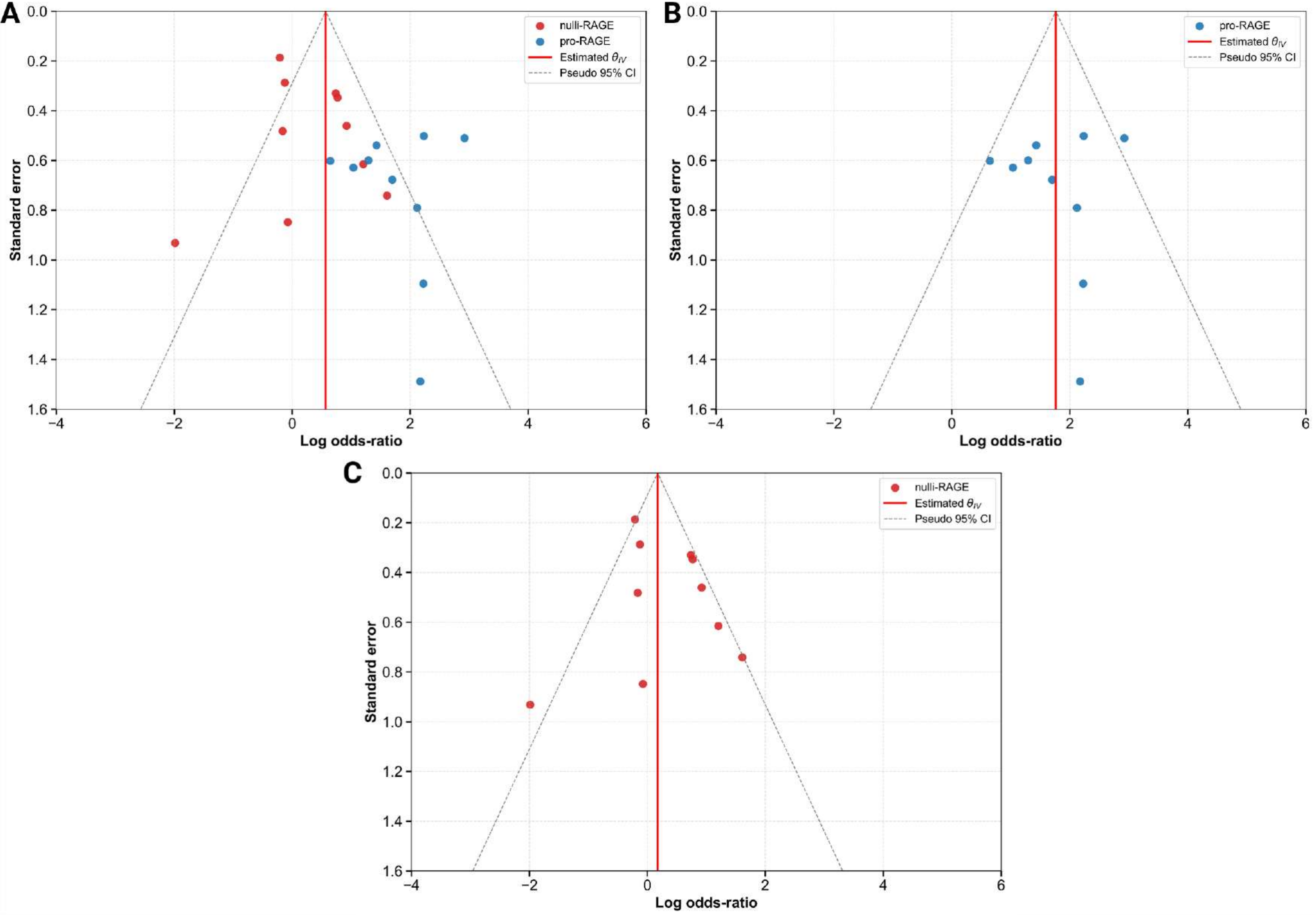

### Supplemental Figure S7

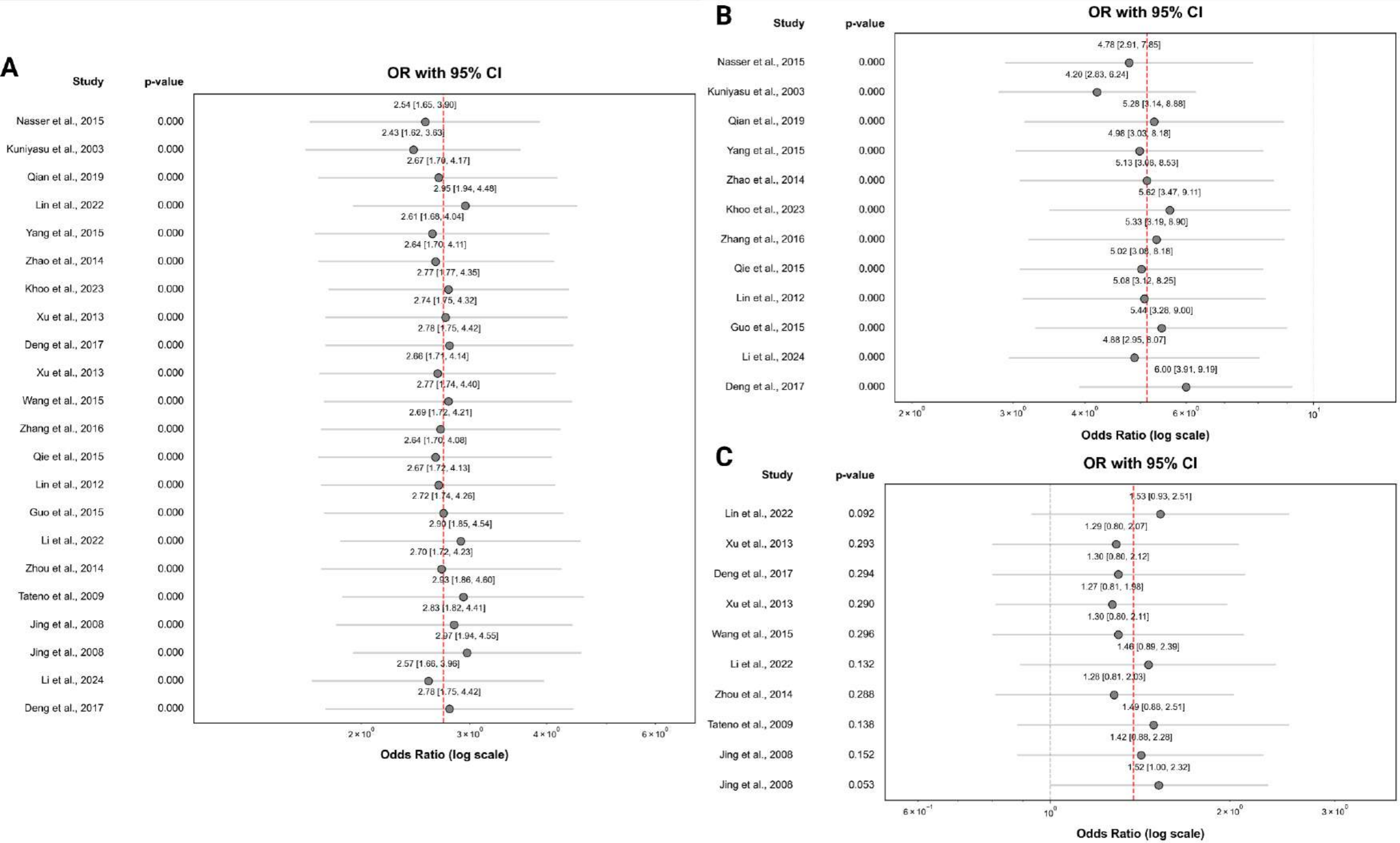

### Supplemental Figure S8

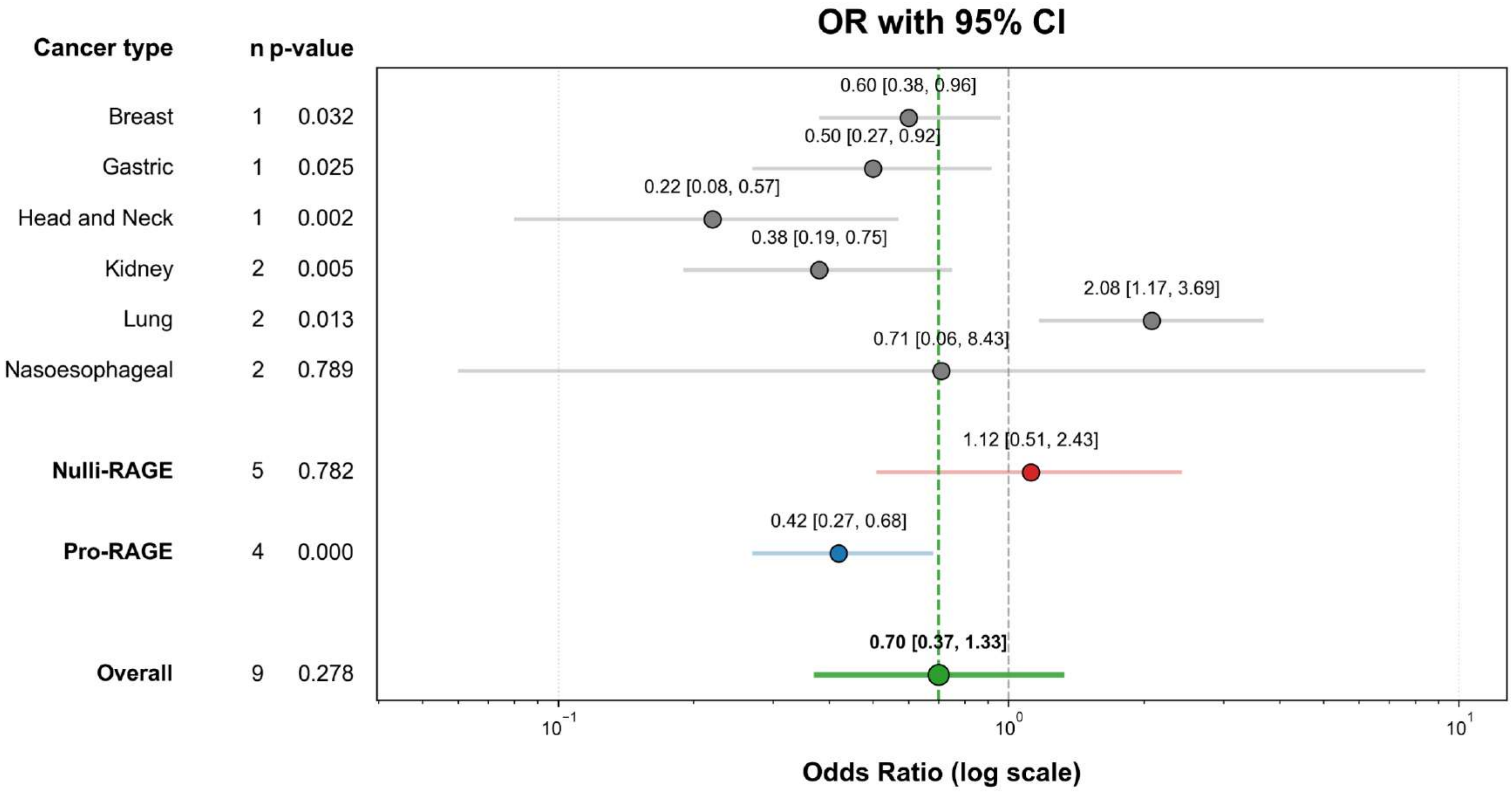

### Supplemental Figure S9

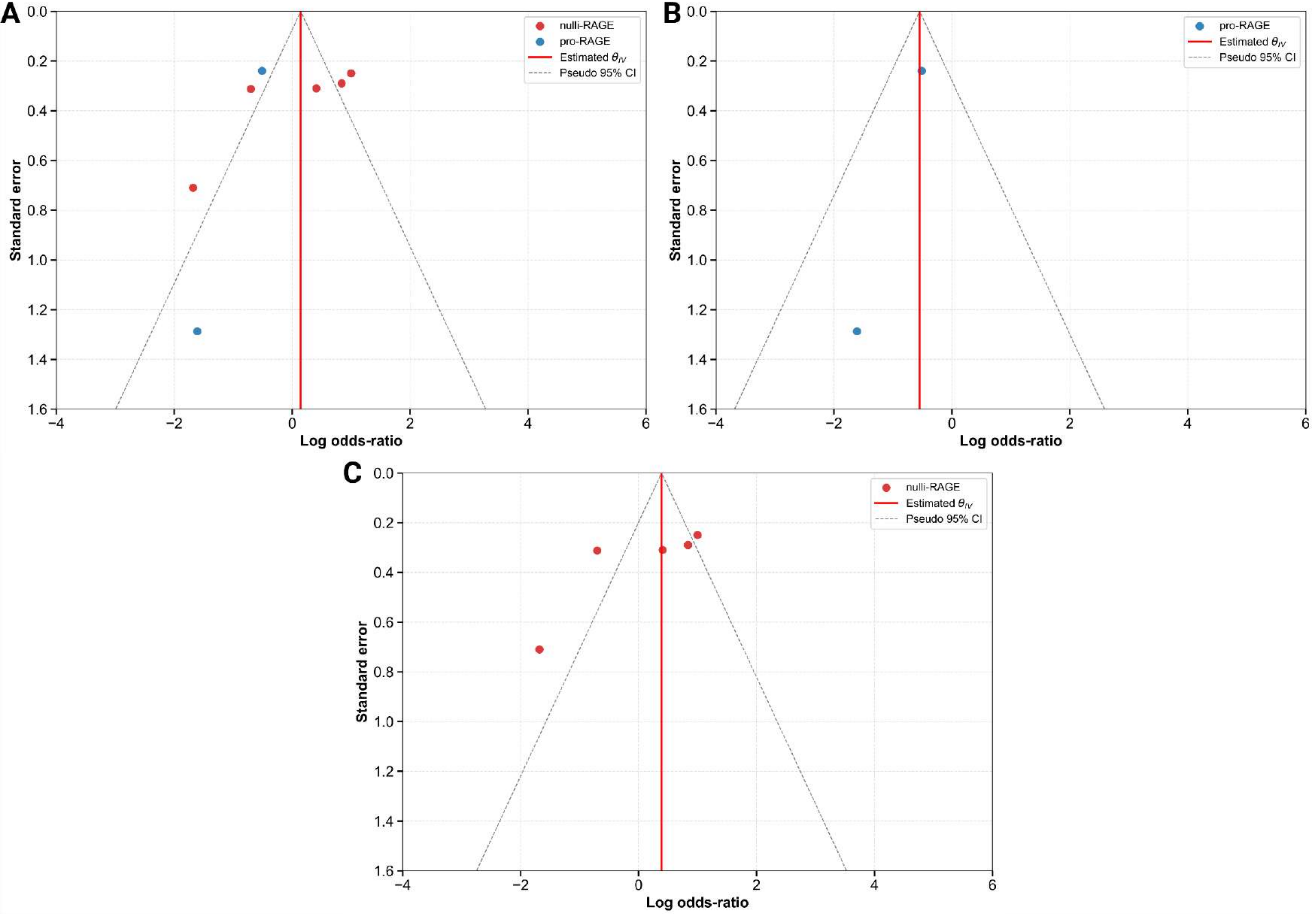

### Supplemental Figure S10

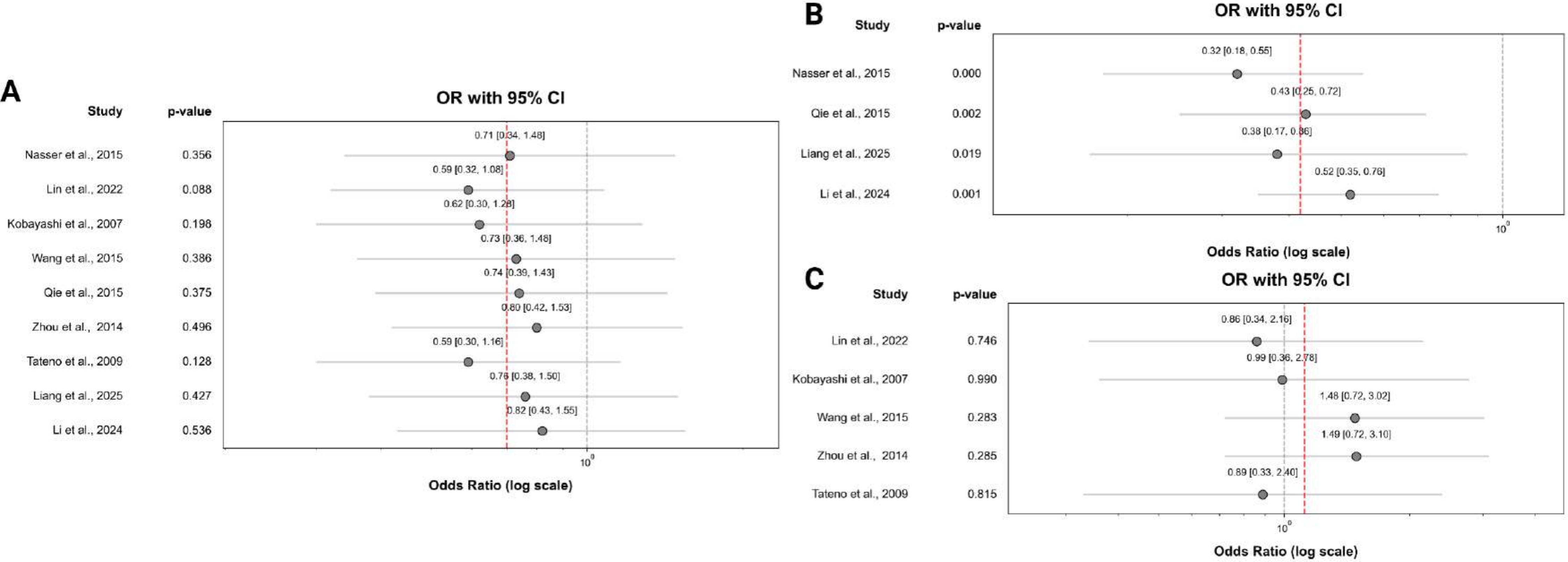

### Supplemental Figure S11

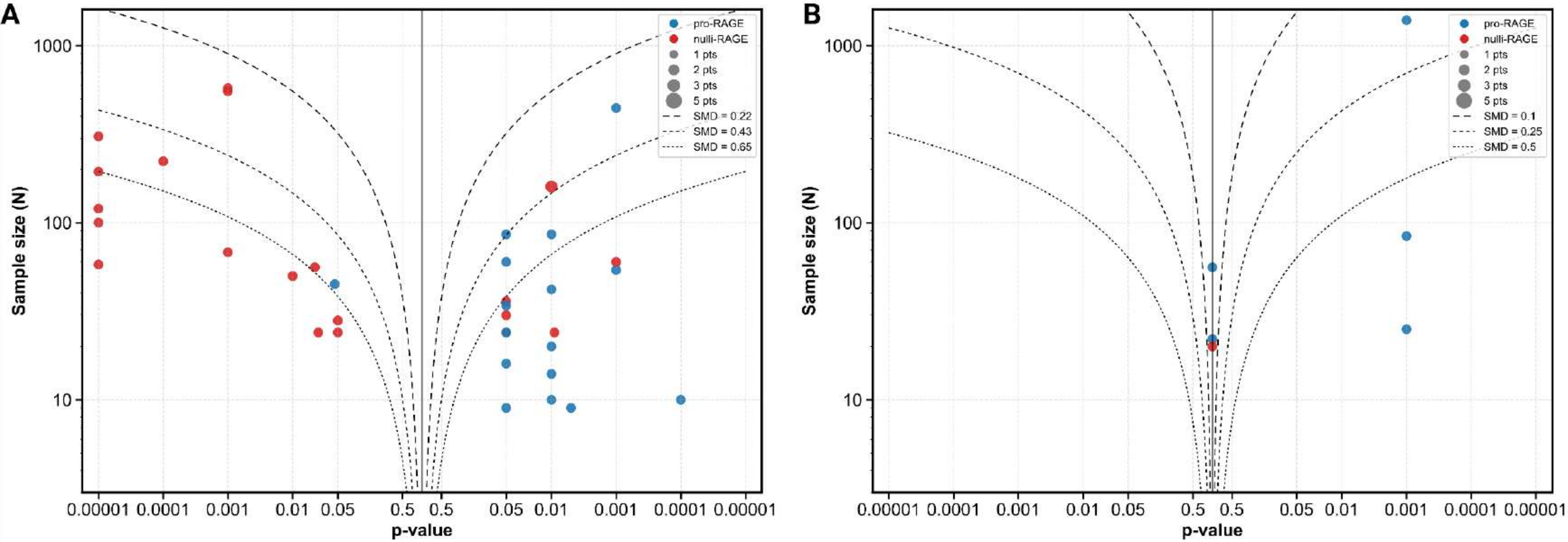
